# Accelerating iterative deconvolution and multiview fusion by orders of magnitude

**DOI:** 10.1101/647370

**Authors:** Min Guo, Yue Li, Yijun Su, Talley Lambert, Damian Dalle Nogare, Mark W. Moyle, Leighton H. Duncan, Richard Ikegami, Anthony Santella, Ivan Rey-Suarez, Daniel Green, Jiji Chen, Harshad Vishwasrao, Sundar Ganesan, Jennifer C. Waters, Christina M. Annunziata, Markus Hafner, William A. Mohler, Ajay B. Chitnis, Arpita Upadhyaya, Ted B. Usdin, Zhirong Bao, Daniel Colón-Ramos, Patrick La Riviere, Huafeng Liu, Yicong Wu, Hari Shroff

## Abstract

We describe theoretical and practical advances in algorithm and software design, resulting in ten to several thousand-fold faster deconvolution and multiview fusion than previous methods. First, we adapt methods from medical imaging, showing that an unmatched back projector accelerates Richardson-Lucy deconvolution by at least 10-fold, in most cases requiring only a single iteration. Second, we show that improvements in 3D image-based registration with GPU processing result in speedups of 10-100-fold over CPU processing. Third, we show that deep learning can provide further accelerations, particularly for deconvolution with a spatially varying point spread function. We illustrate the power of our methods from the subcellular to millimeter spatial scale, on diverse samples including single cells, nematode and zebrafish embryos, and cleared mouse tissue. Finally, we show that our methods facilitate the use of new microscopes that improve spatial resolution, including dual-view cleared tissue light-sheet microscopy and reflective lattice light-sheet microscopy.

## Introduction

Fluorescence microscopy enables imaging with submicron spatial resolution, molecular specificity and high contrast. These attributes allow direct interrogation of biological structure and function, yet intrinsic blurring and noise degrade fluorescence data, yielding an imperfect estimate of the underlying sample. Provided the imaging process can be characterized, such degradation can be partially reversed using deconvolution^1,2^, resulting in improved resolution and contrast. For example, given the point spread function (PSF) and data corrupted by Poisson noise (often dominant in fluorescence microscopy), the Richardson-Lucy deconvolution (RLD)^3,4^ procedure deblurs the estimate of the sample density with each iteration. In addition to deblurring, deconvolution can be used to combine multiple independent measurements taken on the same sample to produce an improved overall estimate of the sample^5^. This approach is especially useful in reconstructing super-resolution images in structured illumination microscopy^6,7^ or in performing joint deconvolution to improve spatial resolution in multiview light-sheet microscopy^8-12^.

Iterative deconvolution has been useful in these applications, but still suffers multiple theoretical and practical drawbacks. A known but unsolved issue is the lack of a well-defined ‘stopping criterion’. There is no general method for determining the optimal number of iterations – if too few, the improvement obtained is less than ideal; if too many, spatial frequencies are artificially enhanced beyond the resolution limit and noise is amplified, resulting in obvious image artifacts. A related practical problem is the computational burden associated with iterative deconvolution, which scales with the number of iterations. While manageable for single-view microscopes, deconvolving large multiview datasets can take days^12,13^, in many cases drastically exceeding the time for data acquisition.

Here we develop tools that address these problems. First, we adapt methods from medical imaging^14^ to fluorescence microscopy data, showing that in most cases the number of iterations in RLD can be reduced to 1, fundamentally speeding iterative deconvolution. Second, we optimize 3D image-based registration methods for efficient multiview fusion and deconvolution on graphics processing unit (GPU) cards. Finally, we show that computationally intensive deconvolution with a spatially varying PSF can be accelerated by using convolutional neural networks to ‘learn’ the relevant operations, provided that suitable training data can be assembled. These advances result in a speedup factor of ten to several thousand-fold over previous efforts. We illustrate the advantages on subcellular to macroscopic length scales, using samples that include single cells, zebrafish and nematode embryos, and mouse tissue. In addition to demonstrating improvements on super-resolution and large multiview datasets acquired with state-of-the-art microscopes, we also show that our methods enable the use of new microscopes, including dual-view, cleared tissue light-sheet microscopy and reflective lattice light-sheet microscopy.

## Results

### Drastically reducing the number of iterations in RLD

Iterative deconvolution algorithms attempt to estimate the underlying sample density from noisy, blurred images. Important components of such algorithms are a ‘forward projector’, which describes the mapping from the desired image of the object to the noisy, blurred image measured by the microscope; and a ‘back projector’, which maps the measured image back onto the desired object image. For example, in RLD,

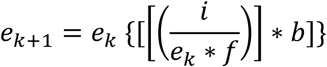

where *e*_*k*_ is the *k*-th (current) estimate of the desired object image *o, e*_*k+1*_ is the *(k+1)*-th (future) estimate, *i* the measured image, *f* the forward projector, *b* the back projector, and * denotes convolution. The PSF is typically used for *f*, since *f* must accurately account for the blurring imparted by the band-limited microscope. *b* is traditionally ‘matched’ to *f* as its transpose (i.e. by flipping the PSF), but this is not the only possible choice. The field of radiology^14^ suggests that using an ‘unmatched’ back projector can accelerate this procedure, but to our knowledge this result has not been exploited in fluorescence microscopy.

We found that the number of iterations in RLD can be greatly reduced if *b* is chosen so that *f* * *b* tends toward a delta function (or equivalently, if the product of the magnitude of the Fourier Transforms (FT) of *f* and *b* approximates a constant in spatial frequency space, **Fig. 1, Supplementary Notes 1, 2**). To study this effect, we began with images acquired with instant structured illumination microscopy (iSIM)^15^, a super-resolution technique. The iSIM PSF, or *f*, resembles a confocal PSF but with smaller spatial extent (**Fig. 1a**). Although *b* is typically chosen to be identical to *f* given the transpose symmetry of the iSIM PSF, we considered other choices with progressively smaller spatial extent (or equivalently, greater amplitude in the spatial frequency passband of the microscope, **Fig. 1b, Methods**). The last of these was a Butterworth filter designed specifically to ‘invert’ the native iSIM frequency response up to the resolution limit, resulting in a much flatter frequency response of |FT(*f*) × FT(*b*)| (**Fig. 1c**). Given its conceptual similarity to a Wiener filter, we termed this choice the ‘Wiener-Butterworth (WB) filter’.

**Fig. 1.**
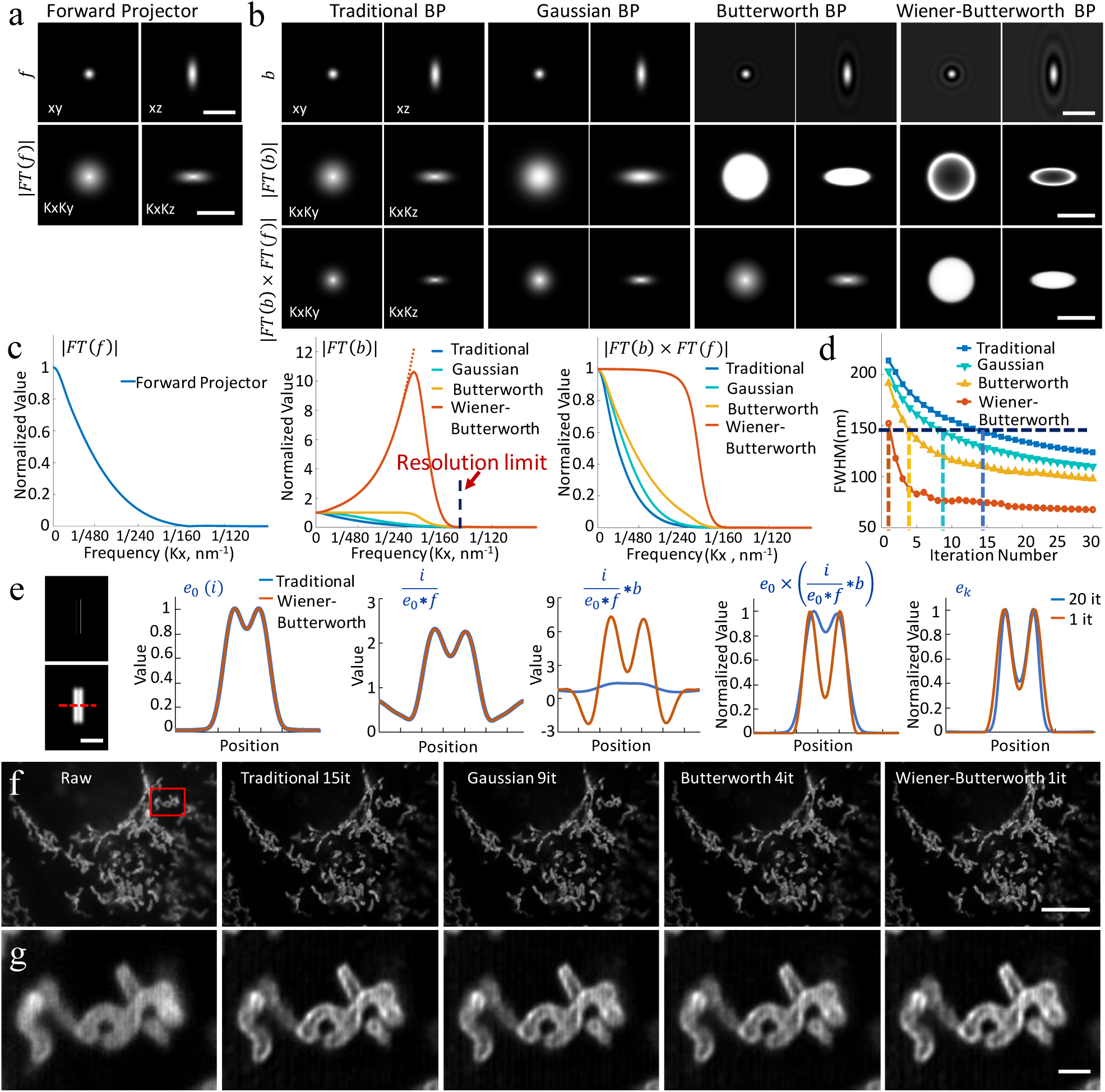
An unmatched back projector reduces the number of iterations required for Richardson-Lucy deconvolution. **a)** Lateral (left) and axial (right) slices through the forward projector for instant structured illumination microscopy (iSIM), shown in real space (top row; PSF) or Fourier space (bottom row, FT(f)). **b**) Different back projectors, including the traditional back projector (transpose PSF) usually employed in RLD, a Gaussian back projector, a Butterworth back projector, and a Wiener-Butterworth back projector. First two rows are as in **a**), the last row shows the product of forward and backward projectors in Fourier space. Note that the colormap for Butterworth and Wiener-Butterworth PSFs have been adjusted to show the negative values (black ringing) that result with these choices, and the colormap for the Wiener-Butterworth Fourier transforms has been adjusted to better show the increase in amplitude at high spatial frequencies. **c)** Line profiles through the Fourier transforms in **a, b**, comparing forward projector (left), back projector (middle), and product of forward and backwards projectors (right). The resolution limit of iSIM is indicated by a vertical dotted line in the middle panel. **d)** The apparent size of a 100 nm bead (vertical axis, average FWHM of 10 beads after deconvolution) as a function of iteration number (horizontal axis) is compared for different back projectors. The resolution limit of iSIM is indicated with a horizontal dotted line. See also **Supplementary Fig. 1. e)** Left: Simulated object consisting of two parallel lines in 3D space (top) and object blurred by the iSIM (bottom). For clarity only a transverse XY plane through the object is shown. Right panels: Line profiles corresponding to red dotted line at left, comparing the effect of original (blue) and Wiener-Butterworth (orange) back projectors in RL deconvolution. The estimate after 20 iterations using the original back projector and only 1 iteration using the Wiener-Butterworth filter is shown in the rightmost graph. **f)** U2OS cells were fixed and immunolabeled to highlight Tomm 20, imaged with iSIM, and deconvolved. Single planes from imaging stacks are shown, with iteration number (it) and back projector as indicated. **g)** Higher magnification views, corresponding to the red rectangular region in **f**). See also **Supplementary Video 1**. Scale bars: **a, b)** 1 *μ*m in top row, 1/100 nm^−1^ in middle, bottom rows; **e)** 1 *μ*m; **f)** 10 µm; **g)** 1 µm.

When deconvolving images of 100 nm beads captured with a homebuilt iSIM, we found that our alternative *b* choices produced a resolution-limited result faster than the traditional back projector (**Fig. 1d, Supplementary Fig. 1**), with speedup factor correlating with the constancy of |FT(*f*) × FT(*b*)|. For example, the WB filter recovered the object’s resolution-limited size with only one iteration, whereas the traditional back projector required 15 iterations. The improved performance of the WB filter does not rely on an improved signal-to-noise ratio (SNR) in the input data (**Supplementary Fig. 2**), nor does it amplify noise more than other methods (**Supplementary Fig. 3**). We also compared the WB back projector to the classic Wiener filter employed in noniterative deconvolution. Here too we found that using the WB filter in RLD outperformed the classic Wiener filter (**Supplementary Figs. 3, 4**). Butterworth and WB back projectors both introduce unphysical negative values into the deconvolved reconstructions (**Fig. 1b, Supplementary Fig. 5**). However, since these values were small and typically located within the noise floor of each image, we set them to zero to yield reconstructions that were nearly identical to the conventional RLD results for these and other datasets presented in the paper (**Supplementary Table 1**).

**Figure 2.**
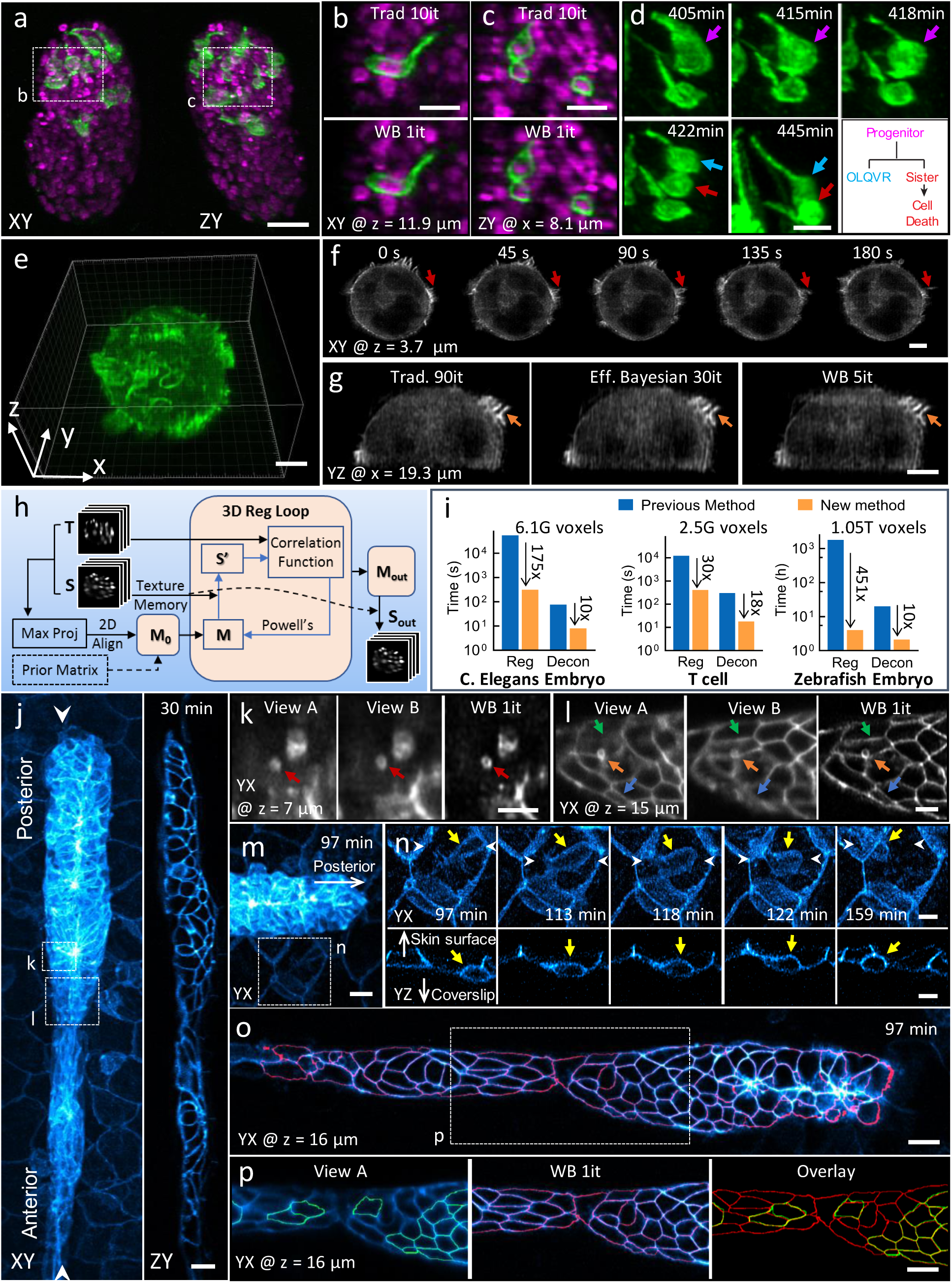
Improvements in deconvolution and registration accelerate the processing of multiview light-sheet datasets. **a**) Lateral (left) and axial (right) maximum intensity projections demonstrate isotropic reconstructions of *C. elegans* embryos expressing neuronal (green, GFP-membrane marker) and pannuclear (magenta, mCherry-histone) markers. Images were captured with diSPIM, and deconvolution was performed using the Wiener-Butterworth (WB) filter. See also **Supplementary Video 4. b, c)** Higher magnification single slices from dotted rectangular regions in **a**), emphasizing similarity between reconstructions obtained with traditional Richardson-Lucy deconvolution (‘trad’) and WB deconvolution. Iterations (it) for each method are displayed. **d)** Higher magnification maximum intensity projection view of neuronal dynamics, indicating neurite extension and terminal cell division for progenitor (purple arrow), OLQVR (blue arrow) and apoptotic sister cell (red arrow). See also lower right schematic and **Supplementary Video 5. e)** WB reconstruction of Jurkat T cell expressing EGFP-actin, raw data captured in a quadruple-view light-sheet microscope. **f)** Selected slices 3.7 *μ*m from the coverslip surface. Indicated time points display fine actin dynamics at the cell periphery. See also **Supplementary Video 6. g**) Axial slice through sample, indicating close similarity between traditional, efficient Bayesian, and WB deconvolution with iteration number as indicated. **h**) Schematic of GPU-based 3D registration used for multiview fusion. Example inputs are two 3D images, referred as the source (S, image to be registered) and target image (T, fixed image). Maximum intensity projections of the input 3D images are used for preliminary alignment and to generate an initial transformation matrix (M_0_). Alternatively, a transformation matrix from a prior time point is used as M_0_. A 3D registration loop iteratively performs affine transfomations on S (which is kept in GPU texture memory for fast interpolation), using Powell’s method for updating the transformation matrix by minimizing the correlation ratio between the transformed source (S’) and T. **i**) Bar graphs showing time required to process the datasets in this figure (left, middle and right columns corresponding to datasets in **a, e** and **j**, respectively, with voxel count as indicated) conventionally and via our new methods. The conventional registration method was performed using an existing MIPAV plugin (see **Methods**) using CPUs while the new registration method was performed using GPUs. Both deconvolution methods were performed with GPUs. Note log scale on ordinate, and that the listed times apply for the entire time series in each case. **j)** Representative lateral (left, maximum intensity projection) and axial (right, single plane corresponding to white arrowheads in left panel) images showing 32-hour zebrafish embryo expressing Lyn-eGFP under the control of the ClaudinB promoter, marking cell boundaries within and outside the lateral line primordium. Images were captured with diSPIM, Wiener-Butterworth reconstructions are shown. Images are selected from the volume 30 minutes into the acquisition, see also **Supplementary Video 7. k, l)** Higher magnification views of dotted rectangles in **j)**, emphasizing improvement in resolving vesicles (red, orange arrows) and cell boundaries (green, blue arrows) with WB deconvolution compared to raw data. Note that **k, l** are rotated 90 degrees relative to **j. m**) Higher magnification view of leading edge of lateral line, 97 minute into the acquisition. **n)** Higher magnification view of dotted rectangular region in **m**), emphasizing immune cell (yellow arrow) migration between surrounding skin cells. White arrowheads are provided to give context and the white arrows point towards skin surface and coverslip. Top row: maximum intensity projection of lateral view, bottom row: single plane, axial view. See also **Supplementary Video 8. o**) Lateral slice through primordium, with automatically segmented cell boundaries marked in red. See also **Supplementary Video 9. p**) Higher magnification view of dotted rectangle in **o**), showing differential segmentation with raw single-view data (green, left) vs. deconvolved data (red, middle). Overlay at right shows common segmentations (yellow) vs. segmentations found only in the deconvolved data (red). Note that ‘z’ coordinate in **j-p** is defined normal to the coverslip surface. Scale bars: 10 *μ*m in **a), m), o)** and **p)**; 5 *μ*m in all other panels.

**Fig. 3.**
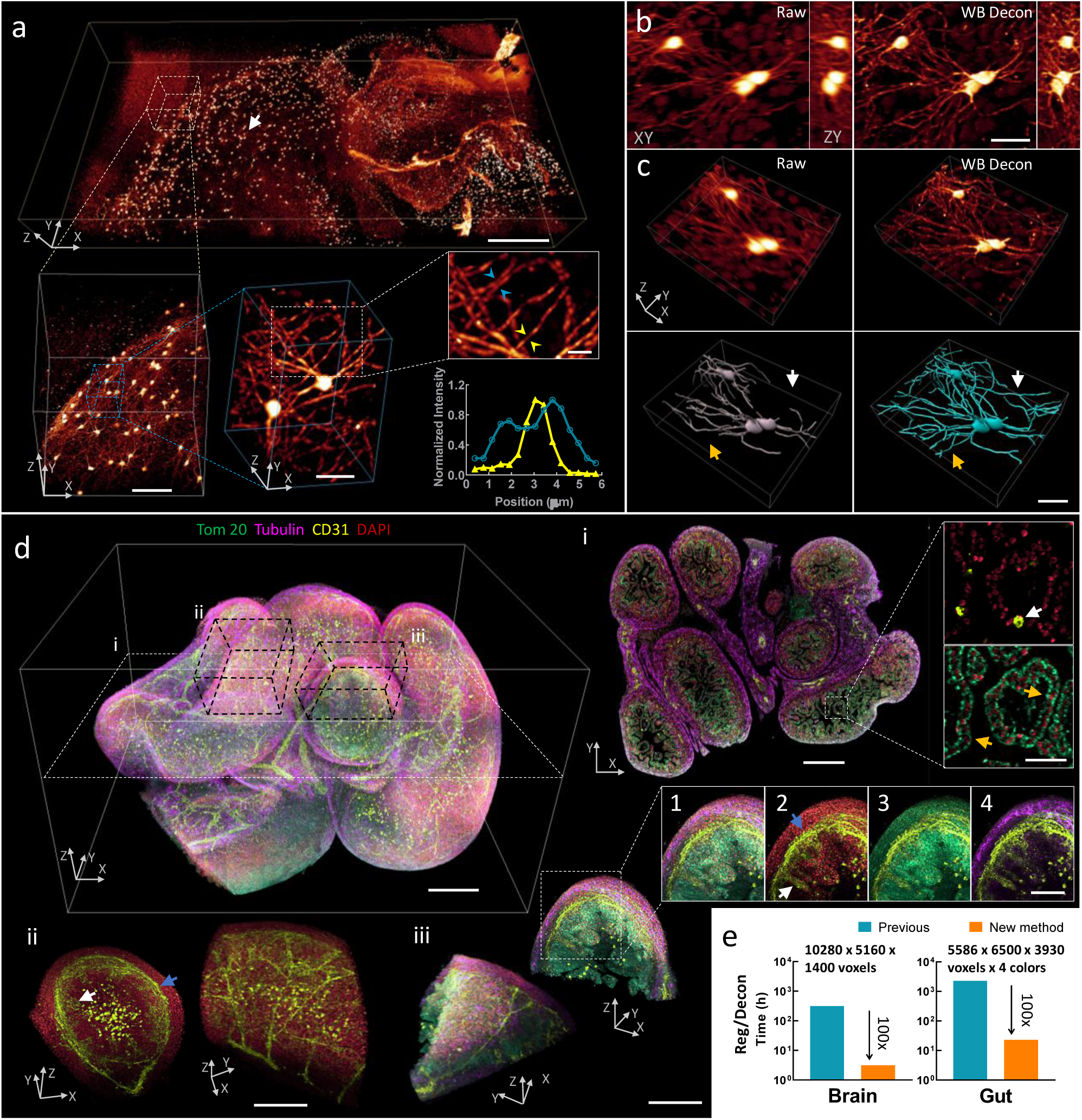
Imaging mm-scale cleared tissue volumes with isotropic micron-scale spatial resolution. **a)** 4 × 2 × 0.5 mm^3^ volume of brain from fixed and iDISCO+-cleared V1b mouse, immunolabeled with Alexa Fluor 555 secondary antibody against tdTomato primary antibody, imaged with cleared tissue diSPIM, and reconstructed after dual-view registration and Wiener-Butterworth (WB) deconvolution. Progressively higher resolution subvolumes are indicated, with line profiles indicating 1.3 *μ*m neurite FWHM (yellow arrowheads) and 1.9 *μ*m separation between neurites (blue arrowheads). See also **Supplementary Video 10. b**) Lateral/axial cross sections from region indicated with white arrow in **a**), emphasizing the higher resolution obtained with WB deconvolution compared to raw single-view data. **c**) Volume renderings of region displayed in **b**), again comparing raw data to deconvolution. Manually traced neurites are shown in bottom row; colored arrows indicate neurites traced in deconvolution that are obscured in raw single-view data. **d**) 2.1 × 2.5 × 1.5 mm^3^ intestinal volume from fixed and iDISCO-cleared E18.5 mouse; labeled with DAPI (red), Alexa-647 conjugated secondary antibody against Tomm20 primary antibody (green), Alexa-488 conjugated secondary antibody against CD31/PECAM-1 primary antibody (yellow), and Alexa-568 conjugated secondary antibody against *α*-Tubulin primary antibody (purple); imaged with cleared tissue diSPIM; and reconstructed after dual-view registration and WB deconvolution. See also **Supplementary Video 11**. i: Single plane demarcated by dotted white rectangular region at left, showing 4-color cross section and higher magnification dual-color views highlighting hollow blood vessel (white arrow) and mitochondria surrounding individual nuclei (orange arrows). ii: Subvolume demarcated by dotted black parallelepiped above, illustrating different perspectives of vascular plexus supplying submucosa (blue arrow) and mucosa (white arrow) of intestine. iii: Different perspectives of four-color subvolume demarcated by dotted black paralleped above and insets 1-4, highlighting hierarchical organization within intestine, e.g., submucosa (blue arrow) and mucosa (white arrow) (inset 2); mitochondrially-enriched regions that support the high energy demand and constant cellular renewal within the mucosa (inset 3); outer intestinal wall with dense alpha-tubulin staining (inset 4). See also **Supplementary Fig. 15. e**) Bar graphs showing the registration and deconvolution time required for post-processing datasets (image sizes in **a**) and **d**) as indicated), comparing previous (blue) and new (orange, 100-fold reduction in time) post-processing methods. Note that times for previous method are estimated (see **Methods** for further detail) and the log scale on the ordinate axes. Scale bars: **a)** 500 µm, 100 µm, 30 µm and 10 µm for progressively higher magnifications; **b**) and **c**) 30 µm; **d**) top left 300 µm, i: 300 µm and 30 µm for insets, ii: 200 µm, iii: 200 µm and 100 µm for insets. See also **Supplementary Videos 12-14**.

**Figure 4.**
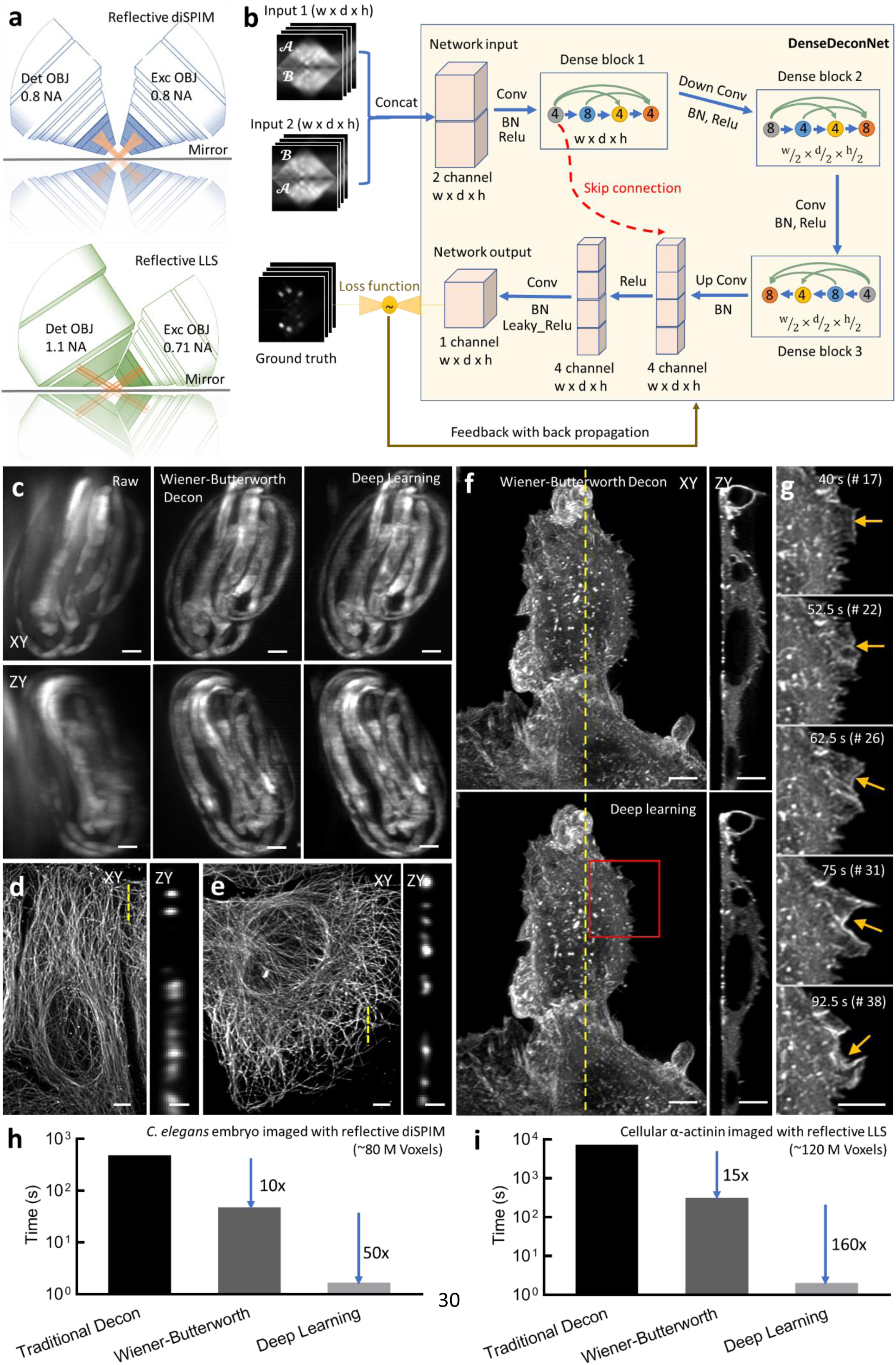
Deep learning massively accelerates deconvolution with a spatially varying PSF. **a)** Reflective imaging geometries for diSPIM (top) and lattice light-sheet (LLS, bottom) microscope. In both cases, the sample is deposited on a reflective coverslip (mirror), which produces additional views of the specimen. **b)** Schematic architecture of our convolutional neural network (‘DenseDeconNet’) used for deep learning. Inputs are concatenated (‘Concat’) image volumes (each containing width (w) × depth (d) × height (h) voxels) obtained from the microscope, which may contain multiple views (𝒜, ℬ) of the specimen. Three ‘dense blocks’ extract feature maps (circles) from the network input, eventually learning to reverse the spatially varying blurring imparted by the microscope by minimizing the difference (loss function) between the network output and the ground truth reconstruction via back propagation. Conv: convolution; BN: batch normalization; ReLu: rectified linear unit. Circles within each dense block unit show the number of feature maps after each convolutional layer, colored arrows within each dense block show the concatenation of successive layers in the network. See **Supplementary Note 3** for more details on the network architecture. **c**) Three-fold *C. elegans* embryos expressing GCaMP3 from a *myo-3* promoter were imaged in the reflective diSPIM (150 volumes, each acquired every 350 ms). Maximum intensity projections of raw data (left), Wiener-Butterworth deconvolution (middle), and deep learning (right) reconstruction are shown for lateral (top) and axial (bottom) views. See also **Supplementary Video 15. d**) U2OS cells were deposited on glass coverslips, fixed, the microtubules immunolabeled with anti-alpha tubulin conjugated with Alexa Fluor 488, and imaged with LLS microscopy. Lateral maximum intensity projection (left) and axial slice (corresponding to yellow dotted line at left) are shown. **e)** U2OS cells were deposited on reflective coverslips and fixed, immunolabeled, and imaged as in **d**). Lateral maximum intensity projection (left) and axial slice (corresponding to yellow dotted line at left) are shown. Reconstructions in **d, e)** were performed using traditional deconvolution with a spatially varying PSF. See also **Supplementary Fig. 16. f)** U2OS cells expressing mEmerald-*α*-Actinin were deposited on reflective coverslips and imaged (100 volumes, each acquired every 2.5 s) in the LLS microscope. Reconstructions were performed via Wiener-Butterworth deconvolution (top) and deep learning (bottom). Lateral maximum intensity projection (left) and axial slice (right, corresponding to yellow dotted line at left) are shown. See also **Supplementary Video 16. g)** Higher magnification view of red rectangular region, emphasizing the dynamics of *α*–actinin near cell boundary (yellow arrows). Bar graphs showing time required for processing a single volume traditional deconvolution with spatially varying PSF, deconvolution via the Wiener-Butterworth filter, and deep learning for **h)** dataset shown in **c**) and **i)** dataset shown in **f)**. Note log scale on ordinate. Scale bars: 5 *μ*m in all panels except 1 *μ*m in zy views in **d, e)**.

In a simulation, we examined the relative performance of traditional and WB back projectors in resolving two lines separated by 1.6× the iSIM resolution limit (**Fig. 1e, Supplementary Video 1**). Using the same forward operator *f* affects the RLD procedure equivalently in both cases, but inspection of the term 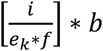 reveals that the WB filter applies a much larger ‘correction factor’ to *e*_*k*_, accelerating production of the final estimate.

We also applied these methods to images of fixed U2OS cells that were immunolabeled to highlight the outer mitochondrial membrane protein Tomm20 and acquired with iSIM (**Fig. 1f, g**). RLD with each of the back projectors improved signal-to-background and spatial resolution relative to the raw data, better revealing interior voids within the mitochondria. As before, however, using the unmatched back projectors also substantially reduced the number of iterations needed (**Supplementary Video 2**), a benefit that that also extended to time-lapse iSIM (**Supplementary Video 3**), confocal, widefield, and single-view light-sheet data (**Supplementary Fig. 6**).

### Accelerating multiview deconvolution and registration

The more than 10-fold improvement in processing speed obtained for single-view deconvolution prompted us to investigate whether our method could also be applied to the more computationally-intensive task of multiview deconvolution. We began by applying our method to dual-view light-sheet microscopy (diSPIM^9^), using the WB back projector instead of the traditional transpose PSF to perform joint deconvolution on the two registered input views (**Methods**). As before, the WB back projector produced nearly identical results to the more traditional method, but with only 1 iteration (**Supplementary Fig. 7**), a 10-fold improvement in speed.

We used our method to reconstruct neuronal dynamics in developing *C. elegans* embryos, obtaining clear images of a subgroup of neurons’ plasma membranes labeled by GFP in a pan-nuclear mCherry background^16^ (**Fig. 2a, Supplementary Videos 4-5**). Post deconvolution, morphologies of neurons and nuclei were sufficiently well-resolved (**Fig. 2b, c**) that we could perform semi-automated lineaging^17^ to identify neurons selectively labeled by the *fmi-1* promoter in this strain. The anterior neurons OLQV(L/R) are glutamatergic sensory neurons that facilitate head foraging and withdrawal reflexes. OLQVs are born after their progenitor cells (AB prpaaappa and AB plpaaappa) undergo a terminal cell division to produce OLQV(L or R) and sister cells (AB prpaaappap and AB plpaaappap) that undergo programmed cell death^18,19^. The progenitor cells first elaborate broad lamellipodial extensions towards the nose of the animal, which eventually become sensory dendrites (**Fig. 2d**). Concomitant with the terminal cell division, the lamellipodial extensions become thinner and longer neurites consistent with the final morphological features of the dendrites. Dendrite extension then continues through what appears to be retrograde extension^20^. Perhaps forces generated during the terminal mitotic division help to create the morphological changes in dendrite shape. Although further experiments are needed to validate this hypothesis, the form of asymmetric division in which the mother cell does not round up during division and one daughter inherits the shape and polarity of the mother has been described previously in fish^21^ and in *C. elegans*^22^. Importantly, our reconstructions allowed us to identify single cells in living embryos, and contextualize the morphological changes undergone by neurons during terminal cell divisions leading to dendrite biogenesis.

Our methods extend to imaging configurations with more views. For example, we acquired a quadruple-view dataset on a triple-objective light-sheet microscope^11^ (**Fig. 2e, Supplementary Fig. 8, Supplementary Video 6**). Stably transfected EGFP-Actin E6-1 Jurkat T cells were plated onto coverslips coated with anti-CD3 antibodies (mimicking antigen presenting cells). After the T cells spread on the coverslip, we imaged them for 30 time points (one time point every 15 s) spanning 7.5 minutes, acquiring 4 volumetric views at each time point. After adapting our deconvolution method for this acquisition scheme (**Methods**), dynamic changes in membrane ruffles and cell protrusions were obvious in the reconstructions (**Fig. 2f**), but obscured in the raw data (**Supplementary Fig. 9**). Using the WB back projector reduced the number of iterations from 90 to 5 (**Fig. 2g**). Importantly, our method also out-performed the state-of-the-art Efficient Bayesian Multiview Deconvolution^10^ (EBMD) method (which required 30 iterations to produce images of similar quality), which can be explained by the flatter frequency response of |FT(*f*) × FT(*b*)| using the WB filter compared to the EBMD result (**Supplementary Fig. 10**).

In processing these dual- and quad-view datasets, we noticed that the time for image registration considerably exceeded the time for deconvolution, usually by 75-120-fold. One approach to faster image registration encases the sample in a labeled matrix, using the multiple feature points from many fiducials to obtain the registration among different views^23^. We opted instead for the less invasive option of greatly accelerating the speed of our image-based registration software. First, we rewrote our CPU-based registration code^9^ in CUDA so that the procedure could be run entirely on our graphics processing unit (GPU). Second, we improved the underlying registration algorithm by incorporating an initial 2D registration and progressively more complex 3D registrations which resulted in faster and more robust performance (**Fig. 2h, Supplementary Fig. 11, Methods, Supplementary Software**). Collectively, these advances resulted in 175- and 30-fold speedups in registration (**Fig. 2i**), respectively, for the modestly sized *C. elegans* and T cell datasets presented in **Fig. 2a, 2e**, which enabled total processing times on par with the acquisition time (**Supplementary Table 2**).

Our improved registration method enabled an even more dramatic speedup (451×, **Fig. 2i**) for a an extended diSPIM acquisition spanning 900 volumes (7.5 hours, 1.05 Tvoxels, 2.1 TB), where we followed the migration of the lateral line primordium in a 32-hour zebrafish embryo expressing Lyn-eGFP under the control of the ClaudinB promoter^24^ (**Fig. 2j, Supplementary Video 7**). Following registration, joint WB deconvolution improved visualization of vesicular structures and cell boundaries compared to the raw data (**Fig. 2k, l**), and facilitated inspection of dynamic immune cells that appeared to migrate in between the skin and underlying somites (**Fig. 2m, n, Supplementary Video 8**). WB deconvolution also substantially improved automated segmentation of cells within the lateral line, as only 71/120 cells were accurately segmented in the raw data vs. 116/120 in the deconvolved data (**Fig. 2o, p, Supplementary Video 9**).

### Submicron isotropic imaging of large, cleared tissue

Other samples that benefit from improved multiview fusion and deconvolution are large volumes of clarified tissue, which can be rapidly imaged using light-sheet microscopes. To explore this possibility, we constructed a cleared tissue diSPIM (**Supplementary Fig. 12**), replacing our original water-immersion objectives with a pair of mixed-immersion 17.9×, 0.4 NA objectives (**Methods**). To estimate spatial resolution, we imaged 100 nm fluorescent beads in dibenzyl ether (Sigma, Cat. # 108014), obtaining single-view lateral full width at half maximum value (FWHM) 0.84 +/− 0.04 µm, and axial FWHM 4.6 +/− 0.4 µm (10 beads, mean+/− standard deviation, **Supplementary Fig. 13**). Registration and 1 iteration of WB deconvolution further improved spatial resolution, resulting in an isotropic 0.79 +/− 0.04 µm, offering a several-fold improvement in axial resolution over previous single-view experiments using the same lenses^25,26^. Next, we fixed, cleared, and immunolabeled mm-scale samples of mouse tissue (**Fig. 3a-d, Supplementary Videos 10-14**) with iDISCO+^27^ or iDISCO^28^, subsequently imaging them with the cleared-tissue diSPIM in stage-scanning mode^29^.

The resulting data span hundreds of gigavoxels – teravoxels, up to ∼2 TB in size. This size presents a major challenge, as such whole raw views do not fit within the memory of single GPU cards and must be subdivided prior to processing. To address this challenge, we created a processing pipeline that scales to arbitrarily large data: cropping the single-view data into subvolumes, registering and deconvolving the subvolumes, and finally stitching the resulting reconstructions back into a higher-resolution composite (**Supplementary Fig. 14**).

In a first example, we imaged a 4 × 2 × 0.5 mm^3^ slab of brain tissue derived from a V1b transgenic mouse, with sparse immunolabeling of neurons and neurites across the entire volume (**Fig. 3a**). The isotropic resolution of the deconvolved reconstruction enabled us to resolve individual neurites at the micron scale (**Fig. 3a**), and to observe fine detail laterally and axially that was not resolved in the raw data (**Fig. 3b**). Manual tracing of neurites was also significantly improved in the deconvolved data relative to the raw data (**Fig. 3c**). In a second example, we performed 4-color imaging on the gut of an E18.5 mouse, spanning a 2.1 × 2.5 × 1.5 mm^3^ volume (**Fig. 3d**). Our reconstruction highlights the organized and hierarchical structure of the intestine, including the interconnected vascular plexus feeding the submucosal and mucosal intestinal areas (PECAM-1, DAPI staining), mitochondrially-enriched regions within the mucosa (Tomm20, DAPI), and tubulin-dense regions within the outer intestinal wall (alpha-tubulin, PECAM-1). As with the brain sample, the isotropic submicron-scale resolution allowed us to visualize fine details that were otherwise obscured by diffraction, including hollow blood vessels and cytoplasmic mitochondria surrounding individual nuclei (**Supplementary Fig. 15**). Importantly, obtaining these as well as other large reconstructions of mouse intestine, stomach and ovary datasets (**Supplementary Videos 12-14, Supplementary Tables 3, 4**) is facilitated by our much faster post-processing methods. Collectively, the new registration (**Fig. 2h**) and deconvolution (**Fig. 1**) methods account for a 100-fold speed improvement over previous efforts, enabling post-processing in tens of hours rather than tens of days (**Fig. 3e**). We note that our method requires orders of magnitude less light dose than a recent technique with similar reported resolution^30^, as our technique confines the illumination to the vicinity of the focal plane.

### Accelerating deconvolution with a spatially varying PSF

Finally, we developed methods for accelerating the deconvolution of fluorescence microscopy data blurred with a spatially varying PSF, acquired by imaging samples deposited on reflective coverslips (**Fig. 4, Supplementary Table 5**). As we previously demonstrated^13^, reflective diSPIM enables the collection of additional specimen views (**Fig. 4a**), increasing information content and boosting spatiotemporal resolution. However, the raw reflective data are contaminated by substantial epifluorescence that varies over the imaging field (**Fig. 4c**). To remove the epifluorescence and fuse the views for optimal resolution enhancement, registration and subsequent deconvolution with a spatially varying PSF are needed. Unfortunately, spatially varying deconvolution carries a considerable computational burden. For example, deconvolving an imaging volume spanning 340 × 310 × 340 voxels with 20 iterations of traditional RLD with a spatially varying PSF requires 1020 3D convolutions (14 minutes per volume with a single GPU card), instead of the 4 3D convolutions required with a spatially invariant PSF (only 2.5 s per volume). Unlike in our previous examples (**Fig. 2, 3**), deconvolution rather than registration becomes the bottleneck in post-processing the raw data.

By modifying the spatially varying RLD update to incorporate the WB filter (**Methods**), we found that only 2 iterations were required to deconvolve a previously published^13^ dataset highlighting calcium waves (marked with GCaMP3) within muscles in 3-fold stage *C. elegans* embryos. As with traditional RLD, the WB modification improved contrast and resolution in the raw data (**Fig. 4c, Supplementary Video 15**), but with a ten-fold reduction in processing time (**Fig. 4h**). These gains also extended to a new form of reflective microscopy, using a higher NA lattice light-sheet (LLS) microscope instead of diSPIM (**Fig. 4a, d-f, i, Methods**).

LLS microscopy^31^ has garnered attention due to its combination of high detection NA and illumination structure; together these attributes result in a better compromise between field-of-view and light-sheet thickness than previous microscopes using pseudo non-diffracting beams. Nevertheless, the contrast and spatial resolution in raw LLS images still suffer from extraneous out-of-focus light due to illumination sidelobes, an effect that can be ameliorated with deconvolution. We found that the performance of the base LLS microscope could be further improved by imaging samples deposited on reflective coverslips (**Fig. 4d-f**), registering the two resulting high NA views oriented ∼113 degrees apart, and deconvolving them with a spatially varying PSF. As assayed with images of immunolabeled microtubules in U2OS cells captured on glass (**Fig. 4d**) and reflective (**Fig. 4e**) coverslips, axial resolution was improved 2-fold, from 750 +/− 39 nm to 379 +/− 23 nm (**Supplementary Fig. 16**). Deconvolving registered images of mEmerald α-actinin in live U2OS cells acquired in the reflective LLS microscope with the WB filter instead of traditional RLD resulted in a 15-fold reduction in processing time (**Fig. 4i, Supplementary Video 16**).

While a 10-15× reduction in processing time is substantial, the time associated with deconvolution still far exceeds data acquisition (3.5 hours to deconvolve the 150-volume *C. elegans* dataset imaged with reflective diSPIM; 13.3 hours to deconvolve the 100-volume α-actinin dataset imaged with reflective LLS microscopy). To obtain further speed enhancements, we turned to deep learning (DL^32^), which has resurged as a promising framework for image classification^33^, image recognition^34^, image segmentation^35^, denoising^36^, super-resolution^37^, and deconvolution^38^.

We constructed a convolutional neural network, terming it ‘DenseDeconNet’, as it is based on linking together dense network blocks^39^ in a memory efficient manner (**Fig. 4b, Supplementary Note 3, Supplementary Software**). These blocks use multiple dense connections to extract features from the raw image stacks, then learn to deblur the images. Unlike previous attempts that deblur 2D image slices by comparing the data to synthetically blurred slices, and average the network output from two orthogonal views to improve resolution isotropy^40^, we designed our method to operate on the full volumetric data, thereby learning the requisite 3D restoration directly. This capability is especially important in reflective applications, in which a simple spatially invariant blur cannot properly model the physics of the microscope.

We began by testing DenseDeconNet on nuclear and membrane-bound labels expressed in live *C. elegans* embryos, acquired on the diSPIM using conventional glass coverslips. We used the deconvolved dual-view data as ground truth. When using only a single view as the input to the network, DenseDeconNet provided resolution enhancement intermediate between the raw data and the deconvolved result (**Supplementary Video 17**). To some extent this is unsurprising; presumably only with both views is there enough information to recover the isotropic resolution provided by diSPIM. However, for highly dynamic structures, the network output with a single-view input sometimes provided more accurate reconstructions than the deconvolved ground truth (**Supplementary Note 3**). We suspect this result is due to the lessened effect of motion blur, which otherwise causes errors in both registration and deconvolution. Additionally, in bypassing the registration, the DenseDeconNet with single-view input provided a 5-fold reduction in total processing time compared to WB deconvolution, i.e. ∼ 1 s for application of DenseDeconNet vs. 5 s for the new registration method (**Fig. 2h**) and 1 iteration WB deconvolution (**Supplementary Table 5**).

Using both registered views for network input enabled resolution enhancement very similar to the ground truth joint deconvolution on data acquired with glass coverslips (**Supplementary Note 3**). This result also extended to the reflective datasets. When training the network using the raw specimen views as inputs and the WB result as the ground truth, DenseDeconNet produced outputs that were nearly identical to the ground truth (based on visual inspection, **Fig. 4c, f;** mean square errors (MSE) of 4.8e-4 (**Fig.4c**) and 5.0e-5 (**Fig.4f**); and structural similarity (SSIM) indices of 0.923 (**Fig.4c**) and 0.965 (**Fig.4f**)), resulting in clear images of calcium dynamics in embryonic muscle (**Supplementary Video 15**) and α-actinin dynamics at the cell boundary (**Fig. 4g, Supplementary Video 16**). Importantly, the network output offered a 50× speed improvement over WB deconvolution (1.68 s/volume, or 500× over traditional RLD) when processing the *C. elegans* data (**Fig. 4h**) and 160× (2 s/volume, or 2400× over traditional RLD) when processing the α-actinin data (**Fig. 4i**).

## Discussion

The WB filter enables RLD with fewer iterations than a traditional back projector, but the potential to introduce artifacts still exists, particularly if too many iterations are applied (**Fig. 1d**). We recommend a single iteration as a good rule of thumb, since this choice resulted in resolution-limited performance on the majority of datasets we examined (**Table S2.1** in **Supplementary Note 2**). With this caveat in mind, the algorithmic improvements we describe here should accelerate image-based biological discovery, especially for the increasingly rich and large datasets that can be obtained with modern light microscopes. For raw data that fit within the memory of a single GPU card (**Fig. 1, 2, 4**), our methods now enable multiview registration and deconvolution on a timescale on par with, and frequently less than, image acquisition. For much larger multiview light-sheet datasets (**Fig. 3**), our approach drastically shortens the post-processing time necessary for image reconstruction, instead placing the bottleneck on file reading, writing, and image stitching (**Supplementary Table 3**). Further speed improvements are possible if these operations are optimized, and our software could benefit from and synergize with state-of-the-art stitching methods^41^. Alternatively, compressing the image data or using multiple graphics cards for additional parallelization^12^ could further shorten post-processing time. We freely provide our software (**Supplementary Software**) in the hope that others may improve it, and strongly suspect that other multiview light-sheet^12^ or light-field configurations^42^ could benefit from our work.

When performing deconvolution with a spatially varying PSF, the WB method provides a substantial speedup over traditional RLD, yet we obtained an even greater acceleration with deep learning. We note several caveats, however, when using deep learning methods. First, enough high-quality training data (for our network, ∼50-100 training pairs) must be accumulated prior to application of the network, underscoring the point that deep learning augments, but does not replace, more classic deconvolution. Second, although application of the trained network takes only seconds per volume, training the network still takes days on a single graphics card. Finally, the networks are ‘brittle’; we obtained optimal results by retraining the network on each new sample. Designing more general neural networks remains an important area for further research.

## Supporting information

Supplementary Software

## Acknowledgements

We thank Owen Schwartz and the Biological Imaging Section (RTB/NIAID/NIH) for supplying the confocal microscope platform and providing technical assistance with experiments, W. Scott Young (NIMH) for providing early access to the V1b mouse line (publication submitted), Jon Daniels (ASI) for advice on integrating the multimmersion objectives into our cleared tissue diSPIM, Matt Anthony (ASI) for providing CAD drawings of our diSPIM assembly, James Shaw (Bitplane) for help with Imaris and the neurite tracing plugin, Andrew Lauziere for his feedback and discussion on the neural network portion of the work, Ryan Christensen for testing aspects of the registration and deep learning pipelines, Evan Ardiel for helping us to acquire the embryonic GCaMP muscle data with the reflective diSPIM, Nico Stuurman for advice in developing ImageJ-compatible software, and Hank Eden and George Patterson for valuable feedback on the manuscript. This research was supported by the intramural research programs of the National Institute of Biomedical Imaging and Bioengineering, the National Institute of Allergy and Infectious Disease, the National Institute of Arthritis and Musculoskeletal and Skin Diseases, the Eunice Kennedy Shriver National Institute of Child Health and Human Development, the National Institute of Mental Health, and the National Cancer Institute within the National Institutes of Health, the National Natural Science Foundation of China (No: 61525106, U1809204, 61427807), the National Key Technology Research and Development Program of China (No: 2017YFE0104000), and Shenzhen Innovation Funding (No: JCYJ20170818164343304, JCYJ20170816172431715). A.U. acknowledges support from NSF awards PHY-1607645 and PHY-1806903. P.J.L. acknowledges support of NIH R01EB107293. H.S., D.C-R. and P.J.L acknowledge the Whitman and Fellows program at MBL for providing funding and space for discussions valuable to this work. D.C-R., R. I., A. S., W.A.M. and Z. B. were supported by NIH grant No. R24-OD016474, L. D. was supported by a Diversity Supplement to R24-OD016474, and M.M. was supported by F32-NS098616. Z.B. additionally acknowledges support via NIH grants R01-GM097576 and the MSK Cancer Center Support/Core Grant (P30 CA008748). A.S. is additionally supported by grant 2019-198110 (5022) from the Chan Zuckerberg Initiative and the Silicon Valley Community Foundation. J. C. W. also acknowledges support from the Chan Zuckerberg Initiative.

## Author Contributions

Conceived project: M.G., Y.L., H.L., Y.W., H.S. Designed experiments: M.G., Y.L., Y.S., T.L., D.D.N., M.W.M., L.H.D., I.R-S., D.G., J.C., H.V., D.C-R., Y.W., H.S. Performed experiments: M.G., Y.L., Y.S., T.L., D.D.N., M.W.M., L.H.D., I.R-S., D.G., J.C., H.V., S.G., T.B.U., Y.W. Prepared samples: Y.S., T.L., D.D.N., M.W.M., L.H.D., R.I., I.R.-S., J.C., H.V., T.B.U., Y.W. Built instrumentation: T.L., H.V., Y.W. Developed and tested deep learning algorithms/software: Y.L., H.L., Y.W. Developed new registration, deconvolution algorithms/software: M.G., Y.L., P.J.L., Y.W. Recognized link between medical imaging algorithms and improved deconvolution: P.J.L. Tested new registration, deconvolution algorithms/software: M.G., W.A.M., Y.W. Developed and tested big data pipeline: M.G., Y.S., Y.W. Contributed lineaging/segmentation software and expertise: D.D.N., A.S., Z.B. Contributed samples: C.M.A., M.H., A.B.C. Wrote manuscript: M.G., Y.L., Y.S., P.J.L., Y.W., H.S. with input from all authors. All authors inspected data and contributed to the drafting of the manuscript. Supervised research: J.C.W., C.M.A., M.H., W.A.M., A.B.C., A.U., T.B.U., Z.B., D.C-R. P.J.L., H.L., Y.W., H.S. Directed research: H.S.

## Methods

### Widefield fluorescence imaging

Widefield imaging was performed on a previously described home-built system. In these experiments, we used a 60× NA=1.42 Oil Objective (Olympus) on an Olympus IX81 inverted microscope equipped with XT 640-W (Lumen Dynamics Group Inc.) as illumination source, and an automated XY stage with an additional Z piezoelectric stage (100 μm range, Applied Scientific Instrumentation, PZ-2000). The illumination was filtered with an excitation filter (ET470/40×, Chroma) and then reflected towards the sample via a dichroic mirror (T495lpxr, Chroma). The emission was collected by the same objective, and filtered with a bandpass emission filter (ET525/50m, Chroma) prior to imaging with an electron-multiplying charge-coupled device (EMCCD) (Evolve Delta, Photometrics). An exposure time of 20 ms and EM gain of 20 were used. The imaging axial step for both beads and fixed actin samples was 150 nm.

#### Fixed, phalloidin labeled actin samples

U2OS cells were cultured on glass bottomed dishes (MatTek, Cat. # P35G-1.5-14C) at 37 C and 5% CO_2_. Prior to labeling, cells were rinsed 3 times with 1× PBS, fixed with 1 mL paraformaldehyde/glutaraldehyde (4%/2%) in 1× PBS for 20 minutes at 37C, rinsed twice in 2 mL 750 mM Tris-HCL pH 7.5 and permeabilized in 0.2% Triton-X/1X PBS for 10 minutes. Next, samples were washed 3 times in staining buffer and blocked in staining buffer containing 1% BSA for 30 minutes. Blocking buffer was removed, and the samples stained with 200 µL of 1:50 Alexa Fluor Phalloidin-488 (Thermo Fisher Scientific, Cat. # A12379):0.2% Tween-20/1× PBS for 1 hour. Cells were washed in 0.2% Tween-20/1× PBS 3 times and imaged in 1× PBS.

#### Bead samples

Glass bottomed dishes (MatTek, Cat. # P35G-1.5-14C) were cleaned with 100% ethanol and coated with 0.1% poly-l-lysine (PLL; Sigma-Aldrich, Cat. # P8920) for 10 minutes. 100-nm yellow-green beads (Thermo Fisher Scientific, Cat. # F8803) were diluted ∼10^5^-fold and 20 μL were added to the coverslip. After 10 minutes, the dish was washed four times with clean water prior to imaging. Bead images were used for estimating the widefield PSFs used in **Supplementary Fig. 6.**

### Confocal imaging

Confocal imaging was performed on a Leica SP8 confocal microscope with 1.40 NA oil lens (HCX PL APO CS 63.0×1.40 OIL UV). The 488 nm argon laser power was set at 20% and the AOTF (488) was set at 5%. The sample was scanned bidirectionally with a voxel size of 48.1 nm in xy and 125.9 nm in z at 200Hz with 6× line average. The pinhole size was set to 20.1 µm (0.21 Airy units). The fluorescence signal was collected from 510 nm to 580 nm with a Leica HyD hybrid detector operating in photon counting mode (10% gain). Data were saved in 8-bit format.

#### Immunolabeled microtubule samples

U2OS cells were cultured on No. 1.5 coverslips (Fisherbrand, Cat. # 12-545-81) at 37 C and 5% CO_2_. Prior to labeling, cells were rinsed 3 times with 1× PBS, fixed with 1 mL methanol for 3 minutes at −20C, and rinsed twice in 2 mL 1× PBS. Next, samples were washed 3 times in staining buffer and blocked in staining buffer containing 1% BSA for 30 minutes. The blocking buffer was removed, and the samples stained with 200 µL of 1:100 anti-alpha Tubulin primary antibody (Thermo Fisher Scientific, 322500) for 1 hour. Cells were washed in 0.2% Tween-20/1× PBS and stained with 200 µL of 1:200 Alexa-488 conjugated Goat anti-mouse secondary antibody (Invitrogen, A11001): 0.2% Tween-20/1× PBS for 1 hour. Finally, cells were washed 3 times in 0.2% Tween-20/1× PBS and twice in distilled water before mounting in Prolong Diamond (Thermo Fisher Scientific, P36961).

### Instant SIM imaging

The instant structured illumination microscopy (iSIM) system has been previously described^13^. For all experiments, a 60× NA=1.42 oil immersion objective (Olympus PlanApo N 60× Oil) was used, resulting in an image pixel size of 55.5 nm and a lateral resolution of ∼150 nm. Fluorescence data were acquired with a pco.edge 4.2 sCMOS camera, and the exposure time was set to 40 ms per image frame. The imaging axial step for beads, immunolabeled mitochondrial samples, and transfected endoplasmic reticulum samples was set to 100 nm, 100 nm, and 500 nm, respectively.

#### Immunolabeled mitochondrial samples

U2OS cells were cultured on glass bottomed dishes (MatTek, Cat. # P35G-1.5-14C) at 37 C and 5% CO_2_. Prior to labeling, cells were rinsed 3 times with 1× PBS, fixed with 1 mL paraformaldehyde/glutaraldehyde (4%/2%) (Electron Microscopy Sciences, Cat. # 15710 and 16120) in 1× PBS for 20 minutes at 37C, rinsed twice in 2 mL 750 mM Tris-HCL pH 7.5 (Corning, Cat. # 46-030-CM), and permeabilized in 0.2% Triton-X (Sigma, Cat. # T9284)/1X PBS for 10 minutes. Next, samples were washed 3 times in staining buffer (0.2% Tween-20 (Sigma, Cat. # P9416)/1× PBS) and blocked in staining buffer containing 1% bovine serum albumin (BSA, Thermo Fisher Scientific, 37525) for 30 minutes. The blocking buffer was removed, and the samples stained with 200 µL of 1:200 anti-Tomm20 primary antibody (Abcam, Cat. # 78547): 0.2% Tween-20/1× PBS for 1 hour. Cells were washed in 0.2% Tween-20/1× PBS and stained with 200 µL of 1:200 Alexa-488 conjugated donkey anti-rabbit secondary antibody (Invitrogen, Cat. # A21206) for 1 hour. Finally, cells were washed 3 times in 0.2% Tween-20/1× PBS and imaged in the instant SIM in 1× PBS.

#### Transfected ER samples

U2OS cells were cultured in 1mL media using MatTek glass bottomed dish at 37 C and 5% CO_2_. At 80% confluency, cells were transfected with 100 µL of transfection buffer containing 2 µL of X-treme GENE, 2 µL plasmid DNA (ERmoxGFP^43^, Addgene Cat. # 68072, 420 ng/µL), and 96 µL of PBS. Cells were imaged 1 day after transfection.

#### Beads samples

Yellow-green fluorescent beads (Thermo Fisher Scientific, Cat. # F8803, 100 nm diameter) were used for experimental FWHM measurements for iSIM. Beads were diluted from the stock concentration 1:1,300 (1:100 in distilled water and 1:13 in ethanol) and spread over cleaned glass cover slips. After air-drying for 5 minutes, coverslips were washed twice in distilled water to remove unattached beads. After air-drying again, beads were mounted in oil (Cargille, Cat. # 16241) onto glass slides and sealed with nail polish.

### Fiber-coupled diSPIM imaging

We used our original fiber-coupled diSPIM system^44^ in addition to another, recently described fiber-coupled diSPIM system^45^ to acquire volumetric time lapse datasets of zebrafish embryo lateral line and nematode embryo neurodevelopment, respectively. Data were acquired in light-sheet scan mode (scanning the light sheet through the stationary sample) with the ASI diSPIM Micromanager^46,47^ (http://dispim.org/software/micro-manager) plugin instead of the LabVIEW control software used previously^44^. For zebrafish data, the XY stage was manually moved periodically in order to ensure that the growing tip of the lateral line did not exit the field of view.

#### Nematode embryos

The 718 bp promoter in plasmid DACR3078 [*fmi-1p(718bp)(EcoRV-EcoRV)::Syn21-GFP-CAAX::p10 3’UTR*] is a bashed fragment from the 3186 bp promoter upstream of the *fmi-1* start codon. To make plasmid DACR3078, EcoRV was used to digest plasmid DACR2984 [*fmi-1p(3186bp)::Syn21-GFP-CAAX::p10 3’UTR*] followed with subsequent religation. Transgenic strain DCR6371 was made by injecting plasmid DACR3078 at 50 ng/⍰l into the lineaging strain, BV514, which ubiquitously expresses the *mCherry::Histone* reporter constructs, *pie-1p::mCherry::H2B::pie-1 3’UTR* and *nhr-2p::his-24::mCherry::let-858 3*’*UTR*^16^. From a spontaneous integration of DACR3078 into BV514, *olaIs98* was isolated. The integrated strain was designated as DCR6371. The Syn21 and p10 3’UTR is a translational enhancer system used in *Drosophila* to boost translational expression^48^. We have found that this also seems to help boost expression in the worm.

Worms were cultivated at 20°C on nematode growth medium seeded with a lawn of *Escherichia coli* strain OP50 using standard methods. Embryos were laid by gravid adults and picked from the plate into M9 buffer with 0.25% Methylcellulose, and then pipetted onto a poly-l-lysine-coated coverslip and imaged in M9 buffer, as previously described^9^. Samples were imaged every 100 s for 50 timepoints with both 561 nm and 488 nm lasers. Further details are available in ref.^45^.

#### Zebrafish embryos

For Zebrafish posterior lateral line imaging, ClaudinB:lynGFP^24^ embryos at 30-32 hpf were placed in embryo media (60 mg RedSea Coral Pro Salt (Drs Foster and Smith Pet Supplies) per liter ddH2O) supplemented with 600 μM MS-222 (Sigma, E10521). For diSPIM imaging, embryos were mounted in 1% low melt agarose (Cambrex, 50080), covered with embryo media, and the agarose above the posterior lateral line primordium was manually removed using forceps prior to imaging.

### Quad-view light-sheet microscopy

We modified our previously described triple-view SPIM system^11^ to acquire 4 volumetric views. Two 40×, 0.8 NA water-immersion objectives [(OBJ A and OBJ B in **Supplementary Fig. 8**, Nikon Cat. # MRD07420] were used in an free-space coupled diSPIM configuration^9^. A 60×, 1.2 NA water-immersion objective (OBJ C in **Supplementary Fig. 8**, Olympus UPLSAPO60XWPSF) was mounted beneath the coverslip. Each objective was housed within a piezoelectric objective positioner (PZT, Physik Instrumente, PIFOC-P726), enabling independent axial control of each detection objective.

Four volumetric views were obtained with the three objectives in stage-scanning mode, i.e., samples were translated though the light sheet via an XY piezo stage (Physik Instrumente, P-545.2C7, 200 μm × 200 μm). When excitation was introduced from OBJ B, one top view (collected from OBJ A) and one bottom view (from OBJ C) were simultaneously acquired. Similarly, when illumination was introduced from OBJ A, another top view (collected from OBJ B) and bottom view (from OBJ C) were again simultaneously acquired. Views collected from OBJ A/B were acquired as usual in light-sheet microscopy (i.e. they are perpendicular to the illumination); views collected from OBJ C were acquired by scanning OBJ C vertically during each exposure. Thus, the top two sCMOS cameras corresponding to OBJ A/B were operated in hybrid rolling/global shutter mode, but the lower camera was operated in a virtual confocal slit mode to obtain partially confocal images during light-sheet illumination introduced from OBJ A/B.

#### T cells

Stably transfected EGFP-Actin E6-1 Jurkat T cells were grown in RPMI 1640 medium with L-glutamine and supplemented with 10% FBS, at 37°C in a 5% CO_2_ environment. Glass coverslips (24 mm × 50 mm × 0.17 mm, VWR, Cat. # 48393241) were coated with 0.01% Poly-L-Lysine (weight/volume) (Sigma-Aldrich, St. Louis, MO) and incubated with Anti-CD3 antibody (Hit-3a, eBiosciences, San Diego, CA) at 10 μg/ml for 2 h at 37°C the same day that cells were imaged. Before imaging, 1 ml of cells was centrifuged at 250 RCF for 5 min, resuspended in the L-15 imaging buffer supplemented with 2% FBS, and plated onto the coverslips.

### Cleared tissue imaging

We modified our original fiber-coupled diSPIM^44^ for cleared tissue imaging by incorporating elements of the commercially available Applied Scientific Instrumentation (ASI) DISPIM and the DISPIM for Cleared Tissue (CT-DISPIM). All components were designed and manufactured by ASI unless otherwise specified. The microscope body was built inside an incubator box (RAMM-Incu) on a 450 mm × 600 mm breadboard (Incu-breadboard). Samples were placed on a FTP-2000 Focusing Translation Platform to provide precise and repeatable x,y,z positioning of the sample as well as rapid stage scanning^29^ during cleared tissue imaging. CAD drawings of the setup are shown in **Supplementary Fig. 12**.

Dovetail mounts (DV-6010) were attached to the SPIM head (SPIM-DUAL-K2) lower Cube III modules and connected to angled dovetails on support arms from posts mounted to the breadboard (Camera Support Kit CAM_SUP-K4-13-5). This configuration fixes the SPIM head while the sample can be moved relative to the head using the FTP-2000, minimizing vignetting of the fluorescence emission that compromised earlier diSPIM performance on large samples.

Each camera (Hamamatsu Orca Flash 4.0) was attached to a tube lens assembly (MIM-Tube-K) which was clamped to Ø1.5” support posts (Thorlabs) from the breadboard leaving an air gap of 1-2mm between the tube lens assembly and the SPIM head. The resulting vibrational decoupling of the cameras from the SPIM head minimized image jitter caused by the camera fans. The cameras themselves were additionally supported on 45° angle brackets (Thorlabs AP45) mounted on Ø1.5” vibrationally damped posts (Thorlabs DP14A).

For cleared tissue imaging we used a pair of Special Optics 0.4 NA cleared tissue immersion objectives (ASI 54-10-12). At the refractive index of the solvent we used (dibenzyl ether), the magnification of these lenses is ∼17.9. Since the back focal planes of these objectives are at different location than the Nikon 40× 0.8 NA water immersion objectives used for live work, the excitation scanners and their associated tube lenses were mounted to adjustable spacers (C60-SPACER-ADJ ASSEMBLY) to ensure 4f spacing of the light-sheet excitation path. All cleared tissue experiments used quad notch filters (Semrock StopLine Notch Filter NF03-405/488/561/635E-25) and associated dichroic mirrors (Semrock BrightLine Laser Dichroic DiO3-R405/488/561/635-t1-25×36), which together isolated the fluorescence from the excitation light (405, 488, 561, 637 nm from Coherent OBIS sources).

Data were acquired by moving the stage in a raster pattern with aid of the ASI diSPIM Micromanager^46^ plugin (http://dispim.org/software/micro-manager, ref.^47^). The number of imaging tiles/rows as well as other acquisition parameters of interest are reported in **Supplementary Table 3**.

Due to the volume size and speed of data acquisition during cleared tissue imaging, it was necessary to use a NVMe solid state drive (Samsung 960 PRO M.2 2TB) to write data during an acquisition. Data were transferred to a local 300 TB server after acquisition for longer term storage.

#### Cleared brain slab

The mouse brain sample was prepared using the iDISCO+ procedure^27^. Briefly, the brain from an adult vasopressin receptor 1B Cre X Ai9 (B6.Cg-Gt(ROSA)26Sortm9(CAG-tdTomato)Hze^49^); Cre recombinase dependent tdTomato) mouse (gift of W. Scott Young, unpublished) was fixed by trans-cardiac perfusion with 4% paraformaldehyde. It was then cut into 2mm slabs and dehydrated through a methanol series, rehydrated, immunolabeled with an antibody that recognizes tdTomato (1:200 dilution Rabbit anti-RFP, Rockland Antibodies and Assays, Cat. # 600-401-379) and an Alexa 555 secondary antibody (Invitrogen, Cat. # A27039 used 1:100), then dehydrated with a methanol series, and dichloromethane before equilibration in dibenzyl ether (Sigma, Cat. # 108014) and imaging.

#### Cleared gut, stomach, and ovary

Mouse tissue stored in 4% paraformaldehyde was dissected and washed in 20 mL 1× PBS for 1 hour at room temperature. Desired organs were dehydrated and rehydrated in a serial dilution of methanol/water and bleached in 5% hydrogen peroxide/methanol mixture according to the iDISCO protocol^28^. After rehydration, pretreated samples were stained with 400 µL of primary antibody dilution (1:100) in a PBS buffer containing 0.5% Triton-X and 0.05% sodium azide and shaken at 37 C for 4 days. Samples were washed in 5 mL washing buffer consisting of 0.5% Triton-X/PBS and 0.05% sodium azide on a rotator for 1 day at room temperature. The next day, samples were stained with 400 µL of secondary antibody dilution (1:100) made of 0.5% Triton-X/PBS and 0.05% sodium azide in a 37 C shaker for 4 days. Samples were washed for one day before optical clearing. For some samples, 1:1000 DAPI (1mg/mL stock) stain was incorporated in the first washing step. All labels are indicated in **Supplementary Table 4**.

Immunolabeled samples were dehydrated in 5 mL of 20%/40%/60%/80%/90%/100% tetrahydrofuran/water mixture (30 minutes at room temperature for every step). Samples were washed in 5 mL of 100% tetrahydrofuran for another 30 minutes at room temperature and incubated in 5 mL of 100% dichloromethane until samples sank to the bottom of the tube. Samples were then incubated overnight at room temperature in another 5 mL of fresh 100% dichloromethane. The next day, samples were cleared in 5 mL of dibenzyl ether (Sigma, Cat. # 108014) twice at room temperature for 30 minutes each time. Cleared samples were mounted on glass slide with a minimal amount of Krazy Glue surrounding the bottom of the samples for imaging with the cleared tissue diSPIM.

#### Beads sample

No. 1.5 coverslips (VWR, 48393241) were cleaned with 100% ethanol and coated with 0.1% poly-l-lysine (PLL; Sigma-Aldrich) for 10 min. Then 100-nm yellow-green beads (Thermo Fisher Scientific; F8803) were diluted ∼10^5^-fold and 20 μL were added to the central region of the coverslip. After 10 min, the coverslip was washed four times with clean water before imaging. During imaging, the beads were immersed in dibenzyl ether (Sigma, Cat. # 108014).

### Free-space coupled diSPIM, conventional and reflective imaging

The geometry of the diSPIM (0.8/0.8 NA) used for conventional and reflective imaging has been previously described^13^. Glass coverslips (24 mm × 50 mm × 0.17 mm, VWR, Cat. # 48393241) for conventional experiments were modified for reflective experiments by sputtering a 150-nm-thick aluminum film over their entire surface and then protecting them with a 700-nm-thick layer of SiO_2_ (Thin Film coating, LLC). During conventional imaging, dual views were sequentially acquired in light-sheet scanning mode via two objectives (Nikon, Cat. # MRD07420, 40×, 0.8 NA) and imaged with 200-mm tube lenses (Applied Scientific Instrumentation, C60-TUBE_B) onto two scientific-grade, complementary, metal-oxide-semiconductor (sCMOS) cameras (PCO, Edge 5.5), resulting an image pixel size of 162.5 nm. During reflective imaging, four views (direct fluorescence and mirror images) were simultaneously collected in stage scanning mode with the same detection optics. In all acquisitions, the exposure time for each plane was 5 ms.

#### Nematode embryos

*C. elegans* were maintained on nematode growth medium seeded with *Escherichia coli* (OP50). Embryos were dissected from gravid adults, placed on poly-l-lysine-coated coverslips and imaged in M9 buffer, as previously described^9^. Strain BV24 [*ltIs44* [*pie-1*p-mCherry::PH(PLC1delta1) + *unc-119*(+)]; *zuIs178* [(*his*-72 1 kb::HIS-72::GFP); *unc-119*(+)] V] was used for imaging nuclei in conventional mode and strain AQ2953 *ljIs131*[*myo-3*p::GCaMP3-SL2-tagRFP-T] for imaging calcium flux within three-fold embryos in reflective mode.

### Lattice light-sheet microscopy, conventional and reflective imaging

The lattice light-sheet microscope (1.1/0.71 NA) for reflecting imaging was constructed as previously described^31^. The annular mask was set at 0.325 - 0.4 NA and a square lattice in the dithered mode was produced at the sample. The excitation power (488 nm) was measured at the back focal plane of the excitation objective at ∼25µW. The 25× Nikon CFI APO LWD detection objective was paired with a 500mm achromat lens for an effective magnification of 63.7×, resulting an image pixel size of 102 nm. The exposure time for each plane was 8 ms, and the stage-scanning step size for the volumetric imaging was 0.4 µm, corresponding to 209 nm along the optical axis after deskewing. When deconvolving the data with a spatially variant PSF for resolution recovery and removal of epifluorescence contamination^13^, the excitation pattern was based on the measured lattice light-sheet dimensions (propagation distance of ∼26.6 µm FHWM along the optical axis and a waist of 0.99 µm FWHM), and the detection PSF was simulated as a widefield PSF with 1.1 NA using the PSF generator ImageJ plugin (http://bigwww.epfl.ch/algorithms/psfgenerator/). The light-sheet dimensions were measured by sweeping the sheet axially through a 0.1 µm diameter fluosphere (ThermoFisher) while stepping the bead along the propagation length of the sheet. Conventional imaging experiments were conducted on 5 mm diameter × 0.15 mm glass coverslips (Warner Instruments, CS-5R). For reflective experiments, 5 mm diameter × 0.17 mm glass coverslips were sputtered as for the free-space diSPIM experiments with a 150-nm-thick film of aluminum followed by a 700-nm-thick layer of SiO_2_ (Thin Film Coating, LLC).

#### Microtubule and actin samples

For imaging microtubules, human osteosarcoma U2OS cells (ATCC HTB-96) were grown on uncoated coverslips, fixed with glutaraldehyde, washed with PBS in room temperature, and then immunolabeled with DM1A antibody conjugated with Alexa-488 (Sigma, T9026). For imaging alpha-actinin, U2OS cells stably transfected with alpha-actinin mEmerald (a gift from Michael Davidson, FSU) were plated onto coverslips 24 hours before imaging. Cells were imaged within 1 h after plating on the reflective coverslips.

### Data processing

#### Dispim deconvolution

The joint RL deconvolution scheme used in diSPIM improves the overall estimate *e* of sample density by alternately considering each view:

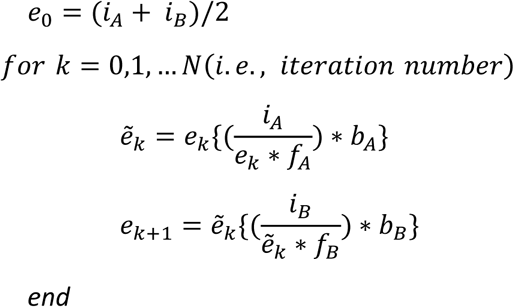

where *i*_*A*_, *f*_*A*_, *b*_*A*_ and *i*_*B*_, *f*_*B*_, *b*_*B*_ are the raw images, forward projector (PSF) and backwards projector corresponding to views A and B, respectively. Traditionally, *b* is taken to be the transpose of *f*. However, as in single-view deconvolution, we found that using unmatched back projectors (e.g. Gaussian, Butterworth, or WB filters) considerably accelerated this procedure (reducing *N*).

#### Quad-view deconvolution

In quadruple-view deconvolution, we start with the additive RLD update, finding as previously reported^11^ that this method yields better reconstructions than the alternating joint deconvolution update used for diSPIM:

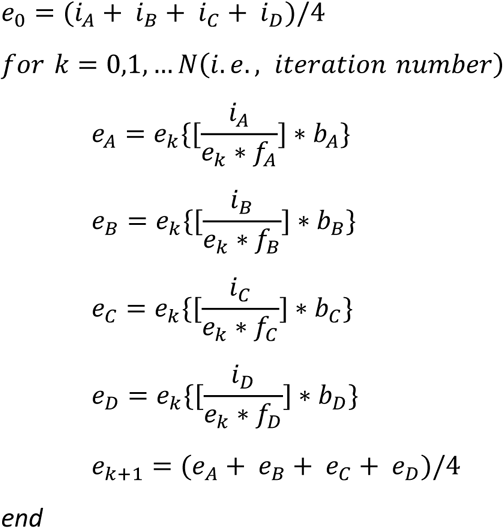

With *f, b, e, i* defined as above and the subscripts *A, B, C*, D indicating each view. Choosing each back-projector *b* to be the transpose of the forwards operator *f* yields the traditional RL update. Choosing the back projectors as follows yields the previously-described ‘virtual-view’ update in EBMD^10^ (* denotes convolution and ^ the transpose), speeding up this procedure:

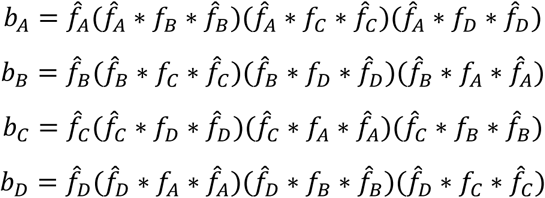

Finally, setting *b* to be the unmatched WB filter appropriate for each view provides the fastest update, as for dual-view and single-view microscopes.

#### Joint deconvolution for reflective light-sheet imaging

Raw image data from the four views in reflective diSPIM imaging (0.8/0.8 NA) or two views in reflective lattice light imaging (0.7/1.1 NA) are merged to produce a single volumetric view, after processing steps that include background subtraction, interpolation, transformation, fusion, registration, epifluorescence removal and joint deconvolution. The data processing steps for removing epifluorescence contamination and enhancing resolution for reflective diSPIM imaging are similar to those previously described^13^, except we modified the forward and backward projectors for Wiener-Butterworth deconvolution as needed. When processing the reflective lattice light imaging data, raw data are transformed so that they are viewed from the perspective of the coverslip. Data processing steps are similar to the asymmetric configuration (0.7NA/1.1NA) used previously in reflective diSPIM imaging^13^, except that the excitation profile is based on the measured dithered lattice light-sheet illumination and the previous forward and backward projectors are modified for WB deconvolution as needed.

In more detail, we form view *U*_1_ (that includes both conventional view and mirrored views) and a second, virtual view *U*_2_ by reflecting view *U*_1_ across the mirror as previously described^13^. *U*_1_ and *U*_2_ are thus blurred with complementary detection PSFs. We register the two views *U*_1_ and *U*_2_, and perform joint deconvolution on them by applying the joint Richardson-Lucy update with WB back projector for each view as follows:

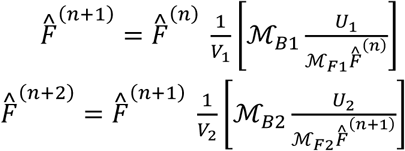

Here, 𝓜_F1_ and 𝓜_F2_ are the forward operators that map the object stack 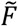 to either measured conventional view stack *U*_1_ or virtually reflected view stack *U*_2_, respectively, and 𝓜_B1_ and 𝓜_B2_ are the backward operators that map from data space back to object space. Four steps are sequentially applied in obtaining each update. First, we compute 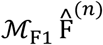 or 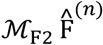 by applying the forward operator 𝓜 _F1_ or 𝓜_F2_ to the current estimate of the object 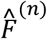 according to three cascaded operations 𝒫ℋ𝒟 at each light sheet position (or *z* slices), where matrix 𝒟 represents multiplication of the estimate 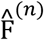 by the crossed light sheets; matrix *ℋ* represents looping over all the *z* slices and performing 2D convolution with a slice of the detection PSF at each *z*; and matrix 𝒫 applies projection over all *z* slices. Second, divide the measured data stack *U* by this quantity, and denote the resulting ratio image *R*. Third, apply the transpose operator 𝓜_B1_ or 𝓜_B2_ to *R*, which involves applying the cascaded operations 𝒟^*T*^ℋ^*T*^𝒫^*T*^ and then summing over all *z* slices. Here 𝒫^*T*^ is a backprojection matrix, which smears the vector to which it is applied back across the image grid; *ℋ*^*T*^ represents looping over *z* in the object distribution and performing 2D convolution with a slice of the transposed but unmatched detection PSF (i.e., WB back projector appropriate for the particular microscope, **Supplementary Note 3**) at each *z*; 𝒟^*T*^ is equivalent to matrix 𝒟, denoting multiplication with the illumination pattern. Last, update the current estimate 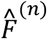 by multiplying by the correction image 𝓜_B1_ or 𝓜_B2_ and dividing by the normalization image *V*_1_ or *V*_2_ (i.e., 𝓜_B1_**1** or 𝓜_B2_**1**, where **1** denotes an image of ones).

#### CPU-based 3D affine registration

CPU-based registration was performed in the open-source Medical Imaging Processing, Analyzing and Visualization (MIPAV) programming environment (http://mipav.cit.nih.gov/). As previously described^9^, we applied an affine transform with 12 degrees of freedom (DOF) to register the source image (S, image to be registered) to the target image (T, fixed image). The DOF matrix is a 12-element transformation matrix that applies the four affine image transformation operations (translation, rotation, scaling and shearing) from S to T. We used an intensity-based method to iteratively optimize the DOF matrix by minimizing a cost function via Powell’s method (http://mathfaculty.fullerton.edu/mathews/n2003/PowellMethodMod.html). We set the search angle range from −10 degrees to 10 degrees, with a coarse angle increment of 3 degrees and a fine angle increment of 1 degree. This registration function ‘Optimized Automatic Image Registration 3D’ has already been incorporated in MIPAV as a Plugin - ‘SPIM-fusion’^44^. With this CPU-based registration environment, we registered the data imaged with diSPIM (**Fig. 2a, j**) and quad-view light-sheet microscopy (**Fig. 2e**, see below for more detail on how we registered four views), and compared the registration outcomes and computation costs with the GPU-based registration described in the following section (**Fig. 2i**). To estimate the computational costs for registering large, cleared tissue volumes with the CPU-based approach (**Fig. 3i**), we randomly chose 10 subvolumes (each 640 × 640 × 640 pixels), calculated the time for registration, averaged the times (i.e., ∼31 mins per subvolume) and then multiplied the averaged time with the total number of subvolumes (e.g., 4576 subvolumes in **Fig. 3d**) to estimate the total registration time (i.e., ∼ 100 days).

#### GPU-based 3D affine registration

We developed a new registration pipeline that accelerates the registration of multiview light-sheet data via GPU programming (**Supplementary Fig. 11**), including data acquired with the diSPIM (**Fig. 2a, j**), quad-view light-sheet microscopy (**Fig. 2e**), reflective diSPIM (**Fig. 4c**) and reflective lattice light-sheet microscopy (**Fig. 4f**). More importantly, this GPU-based registration method also enables the registration of large, cleared tissue datasets imaged with diSPIM (**Fig. 3**), which is impractical if implemented in the CPU-based registration method (e.g., ∼100 days with CPU-based registration time as estimated above vs. ∼24 hours with GPU-based registration for data in **Fig. 3d**).

Our GPU-based method uses the same method (i.e., intensity-based, iterative optimization of the transformation matrix) as in the previous CPU-based registration, but dramatically improves the registration speed and accuracy for several reasons. First, we iteratively perform affine transformations on the source image (S) which is always kept within the GPU texture memory. The main computational burden in 3D transformation is trilinear interpolation, which can be significantly lessened via the use of texture memory. Second, the correlation ratio between the intensity of the transformed source (S’) and target image (T) that is used in the cost function, can be rapidly calculated via the parallel computations enabled by the GPU. Third, when minimizing the cost function by using Powell’s method to update the 12-element transformation matrix, we don’t simultaneously optimize all 12 elements (i.e., full translation, rotation, scaling and shearing that comprise 12 DOF). Instead, the optimization is serial, successively optimizing translation; rigid body (translation and rotation, 6 DOF); translation, rotation and scaling (9 DOF); and finally the full translation, rotation, scaling and shearing operations (12 DOF). We observed that such serial optimization makes registration more accurate and robust. Finally, although the initial transformation matrix (M_0_) for beginning the optimization process is an identity matrix by default, we also provide an option to generate M_0_ by performing a 2D registration (translation and rotation) on the XY and ZY maximum intensity projections of S and T. This 2D registration is an intensity-based rigid body transformation with the same optimization routine as 3D registration, but performing registration in 2D with only translation and rotation is very rapid, only ∼1% of the time required for performing full 3D registration. This additional step also guarantees a reasonable starting initialization of M_0_ for further 3D optimization in 3D. Alternatively, a transformation matrix from a prior time point in a time lapse 4D dataset can be used as M_0_ to accelerate the registration. In some cases (e.g., **Fig. 2a)**, we observed that using a matrix from a previous time point can reduce the registration time for a new volume by as much as 65%, e.g. from ∼8.8 seconds/volume to ∼3.1 seconds/volume.

We implemented this GPU-based registration pipeline in CUDA/C++ (**Supplementary Software**) and called it in Matlab or FIJI to register the data imaged with conventional and reflective diSPIM and LLS microscopy (**Fig. 2a, j, Fig. 4c, f**). To increase registration accuracy for the quad-view data (**Fig. 2e, Supplementary Fig. 9**) acquired with the quad-view light-sheet system (**Supplementary Fig. 8**), we (1) transformed view A and view B into the coordinate system of the bottom views C/D and deconvolved each view to increase image quality; (2) registered the deconvolved view D to the deconvolved view C, thus obtaining a registration matrix mapping view D to view C; applied this registration matrix to the raw view D, thus registering it to the raw view C; (3) registered the deconvolved view B to the deconvolved view A, thus obtaining a registration matrix mapping view B to view A; applied this registration matrix to the raw view B, thus registering it to the raw view A; (4) performed joint deconvolution on the two registered, raw views A and B; (5) registered the jointly deconvolved views A/B to the deconvolved view C, thus obtaining a registration matrix mapping views A/B to view C; (6) applied both registration matrices (view B to view A, then views A/B to view C) to register all raw views to the coordinate system of the bottom views (i. e., view C/D). For deconvolving time series data (**Fig. 2e, Supplementary Video 6**), we applied this process to the first time point in each view, obtaining a set of registration matrices that were then applied to all other time points in the 4D dataset.

#### Post-processing pipeline for large, cleared tissue data imaged with diSPIM

We developed a postprocessing pipeline that can register and jointly deconvolve large datasets imaged with the diSPIM, including the cleared tissue data presented in this paper (**Supplementary Fig. 14**). Such datasets span hundreds of GB – terabytes, a size that exceeded either RAM or GPU memory on our workstation.

First, raw image data recorded by the cameras in the cleared-tissue diSPIM (multiple 16 bit TIFF files, each less than or equal to 4 GB) need to be re-organized and re-saved as TIFF stacks, each corresponding to a distinct spatial strip, color, and view. Second, strips for each color/view are combined with Imaris Stitcher. Third, these TIFF stacks are deskewed (transforming from stage-scanning mode to light-sheet scanning mode), interpolated (obtaining isotropic pixel resolution), rotated (transformed from the objective view to the perspective of the coverslip), cropped (saving memory), and resaved as TIFF files (e.g. ∼ 2 TB for the 4 colors and 2 views acquired for the dataset shown in **Fig. 3d**). Due to the large data size, and our limited memory, we could not directly register the two views via our GPU card, and performing the registration with CPU processing^9^ is impractical due to the ∼100-fold slower processing that would result (**Fig. 3e**). Our strategy for dealing with the GPU memory bottleneck is to down-sample Views A and B by a factor **β**, to View A’ and View B’, such that the total size of the views is reduced by **β^3^** (e.g., 125-fold if **β** = 5). Registering these downsampled volumes can now be achieved in GPU memory, obtaining a registration matrix ***M*_*D*_** that maps view B’ to view A’. A coarse, global 3D affine transformation matrix ***M*_*G*_** that maps view B to view A can then be derived from ***M*_*D*_:**

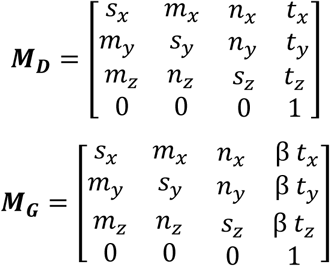

Here the three terms *t*_*x*_, *t*_*y*_, *t*_*z*_ represent translations in each dimension, while the other 9 terms *s*_*x*_, *s*_*y*_, *s*_*z*_, *m*_*x*_, *m*_*y*_, *m*_*z*_, *n*_*x*_, *n*_*y*_, *n*_*z*_ combines scaling, rotation, and shearing in 3D.

Note that ***M*_*G*_** cannot be directly applied to View B to obtain a coarsely registered view B (again due to their large size). But ***M*_*G*_** can be used to crop Views A and B into multiple subvolumes that are sufficiently small that they can be registered (e.g., ∼1000 subvolumes, each 640 × 640 × 640 pixels with an interval of 512 × 512 × 512 pixels, 80% overlapped region in each dimension). If the position of the k-th subvolume in View A is specified by the vector 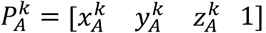, then the starting position of the k-th subvolume in View B can be obtained by:

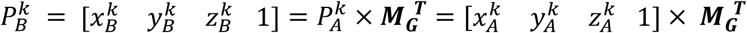

After cropping, this subvolume can be coarsely registered with the corresponding cropped subvolume in view A using a new matrix 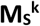, which can be derived from the cropping position matrix 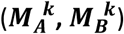 and global transformation matrix ***M*_*G*_**:

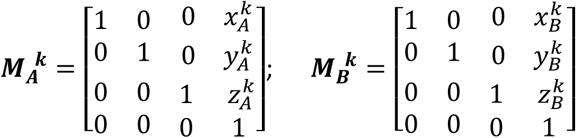

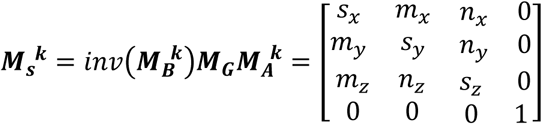

Fine registration and joint WB deconvolution are then applied to the coarsely registered, paired subvolumes of View A and View B. For each deconvolved subvolume (640 × 640 × 640 pixels), boundary regions (45 pixels from each edge, in all three dimensions) are removed to eliminate edge artifacts, and the resulting subvolumes are resaved with size 550 × 550 × 550 pixels. Finally, stitching all deconvolved and newly cropped subvolumes results in the final reconstruction (e.g., ∼1 TB for the dataset displayed in **Fig. 4d**). Note that during the stitch, linear blending is performed on the remaining overlapped regions of the adjacent subvolumes (38 pixels from each edge, in each dimension) to lessen stitching artifacts.

#### Zebrafish segmentation

For segmenting cells in the lateral line primordium (**Fig. 2o, p**), the “morphological segmentation” feature in the MorpholibJ plugin^50^ was used, with identical settings for raw and deconvolved images. Before segmentation, images were blurred in ImageJ using a Gaussian kernel with sigma = 1.5. A watershed tolerance of 15 and a connectivity of 26 was used during the segmentation. Cells in both the raw data and successfully segmented cells in the processed images were manually counted in ImageJ.

#### Full width at half maximum (FWHM) calculations

All FWHM calculations were implemented in Matlab. For statistical measures,values were averaged from 10 simulated beads (**Supplementary Fig. 2**), 10 experimental beads (**Fig. 1d, Supplementary Figs. 1, 3, 6e, f**), or 10 microtubule filaments (**Supplementary Figs. 6d and 15e**).

#### Simulation of images with different SNRs

SNR simulations were conducted in Matlab. For images shown in **Supplementary Fig. 2**, a noise-free image was obtained by blurring 10 point objects with the iSIM PSF (simulated as the product of excitation and emission PSFs). We next added Gaussian nose (simulating the background noise of the camera in the absence of fluoresence) and Poisson noise (proportional to the square root of the signal). We defined SNR as

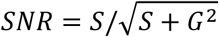

where S is the signal defined by the average of all pixels with intensity above a threshold (here set as 1% of the maximum intensity of the blurred objects in the noise-free image); G is the Gaussian noise (set as 10 counts according to the measured standard deviation of the background noise of the camera). Final images shown in **Supplementary Fig. 2c** were then generated by scaling the signal level S and adding noise according to the equation above to achieve the target SNR.

#### Bleach correction

For several time-lapse datasets (**Figs. 2a, 2e, 2j, 4c, 4f, Supplementary Videos 3, 6, 16, 17**), we performed standard bleaching correction using an ImageJ Plugin (Bleach Correction^51^; https://imagej.net/Bleach_Correction) with the “simple ratio” method.

#### Delining Data

In the mitochondrial dataset acquired with iSIM (**Fig. 1f, Supplementary Fig. 4** and **Supplementary Video 2**), we applied notch filters in Fourier space to suppress slight line artifacts in the raw data, as previously described^52^.

#### Video Compression and Rendering

The zebrafish lateral line volumes shown in **Supplementary Video 9** were median filtered with a 5 × 5 × 5 kernel in Imaris 9.2.1 (Bitplane), and manually segmented with the ‘local contrast’ function at each time point to isolate the immune cell from the skin. The isolated immune cell was then further manually segmented by an absolute intensity threshold to remove unwanted pixels, and finally false colored in red. The isolated lymphocyte was recreated as an independent channel, and false colored in red. **Supplementary Videos 10-14** were also rendered in Imaris 9.2.1 and exported as uncompressed avi files (usually multiple GB in size). These files were JPEG-compressed (down to several hundred MB) in ImageJ and then compressed again in VLC media player using H.264 compression. In some cases, the total image size was also slightly downsampled to achieve the final file size.

### Neural network for deep learning

We developed the DenseDeconNet neural network (**Fig. 4b, Fig. S3.1** in **Supplementary Note 3**) by adapting a densely connected network^39^ for 3D image data. This network consists of three dense blocks and uses multiple dense connections between convolutional layers to extract relevant features from the image volumes, learning the deblurring necessary for image reconstruction. All operations are implemented on 3D data, and thus can directly incorporate 3D information contained within the image stacks to simultaneously improve axial and lateral resolution. The total number of learned parameters in our DenseDeconNet is approximately 18 thousand. The network is optimized using the backpropagation algorithm with the adaptive moment estimation (Adam) optimizer^53^ and a starting learning rate which decays during the training procedure. More detail about this fully convolutional network is described in **Supplementary Note 3**.

In our DenseDeconNet, we designed our objective function with three terms: the mean square error (*MSE*), the structural similarity (*SSIM*) index^54^ and the minimum value of the output (*MIN*). The *MSE* term ensures that the difference between network outputs and ground truths is as small as possible. The *SSIM* term is used to preserve the global structural similarity between the network output and the ground truth. We monitor the *MIN* of the output to avoid negative values.

DenseDeconNet is implemented with the Tensorflow framework version 1.4.0 and python version 3.5.2 in the Ubuntu 16.04.4 LTS operating system. Training was performed on a workstation equipped with 32 GB of memory, an Intel(R) Core (TM) i7 – 8700K, 3.70 GHz CPU, and two Nvidia GeForce GTX 1080 Ti GPU cards with 11 GB memory each. Kernels in the convolution layers were randomly initialized with a Gaussian distribution (mean = 0, standard deviation = 0.1). For an input image 70 MB in size, fully training the network with 10000 iterations took *∼*60 h.

We tested DenseDeconNet on 3D images of membranes and nuclei in live *C. elegans* embryos acquired with diSPIM, images of GCaMP3 expression in live *C. elegans* embryos acquired with reflective diSPIM, and images of *α*-actinin in live cells acquired with reflective lattice light-sheet (LLS) microcopy. The input data are either raw single-view image volumes or dual-view image volumes. The ground truth data consist of traditional R-L joint deconvolution with 10 iterations for diSPIM data (conventional coverslips and reflective coverslips), and R-L deconvolution with the WB back projector with 1 iteration for reflective lattice light-sheet data. All data are derived from volumetric time-series (‘4D’ data); usually 80% of volumes were randomly selected for training and the remaining 20% for validation and testing. The parameters for all datasets used in deep learning are summarized in **Table S3.1** in **Supplementary Note 3**. More detail about the deep learning results are shown in **Fig. 4** and **Figs. S3.2-S3.6** in **Supplementary Note 3**.

### Supplementary Software

We attach our software as a compressed zip file. The software includes four sets of programs for implementing (1) WB deconvolution on a variety of different microscopes; (2) rapid registration of two volumetric images, e.g. for subsequent WB deconvolution; (3) registration and deconvolution of large cleared tissue datasets, imaged with the diSPIM; and (4) our convolutional neural network (‘DenseDeconNet’) for resolution recovery. We also include accessory MATLAB scripts, C++ .h/.dll files, and mex files for virtually reading big TIFF stacks, writing TIFF stacks, and performing 3D convolution in the Fourier domain. Finally, the zip file also includes a README text file that explains how to run our software on a PC with specifications similar to ours (CPU: Intel Xeon, E5-2660-v4, 28 threads; RAM: 256 GB; GPU: Nvidia Quadra M6000 graphics card, 24 GB memory).

The program that implements WB deconvolution includes MATLAB scripts for the WB single-view deconvolution of widefield fluorescence microscopy (**Supplementary Fig. 6**), confocal microscopy (**Supplementary Fig. 6**), instant SIM (**Fig. 1f, Supplementary Fig. 4, Supplementary Video 3**) and light-sheet fluorescence microscopy (**Supplementary Fig. 6**) data; WB joint deconvolution of diSPIM data acquired on glass coverslips (**Supplementary Fig. 7)**; WB additive deconvolution of quad-view light-sheet imaging data acquired on glass coverslips (**Fig. 2e, Supplementary Video 6**); WB deconvolution for data contaminated with a spatially variant PSF taken with a reflective, symmetric diSPIM (**Fig. 4c, Supplementary Video 15**); and WB deconvolution for data contaminated with a spatially varying PSF acquired with reflective lattice light-sheet microscopy (**Fig. 4f, Supplementary Video 17**).

There are two main MATLAB scripts that are used in performing a 12-degree affine registration, one that performs registration by calling the registration dll written in C++/CUDA (**Supplementary Fig. 2e, Supplementary Video 6**); the other for conducting both registration and WB deconvolution for diSPIM data by calling the hybrid registration and deconvolution dll written in C++/CUDA (**Fig. 2a, f, Supplementary Videos 5, 7**).

Registration and joint deconvolution (both traditional and WB deconvolution) for fusing diSPIM data can be also achieved via our custom FIJI plugin. Unlike the MATLAB scripts, in this plugin, users have the options to rotate and interpolate the two perpendicular views for obtaining isotropic pixels before registration. In addition, the plugin can process either single-color or dual-color data. More details can be found from the README text file that explains how to install and use the plugin in ImageJ or FIJI.

The program for registration and deconvolution of large cleared tissue imaged with diSPIM (**Fig. 3, Supplementary Videos 10-14**) includes two main MATLAB scripts, the first one is for pre-processing the raw TIFF data by converting the data from stage scanning mode to the perspective of the coverslip; the second implements the coarse registration, cropping into subvolumes, fine registration, WB joint deconvolution, and stitching back into a large dataset. The last main program includes four main Python scripts for running DenseDeconNet with Tensorflow. These scripts are designed for single-view input training, single-view input validation (**Figs. S3.2, S3.3, S3.5** in **Supplementary Note 3, Supplementary Video 17**), dual-input training, and dual-input validation (**Fig. 4c, f, Figs. S3.4, S3.5, S3.6** in **Supplementary Note 3, Supplementary Videos 15, 16**).

