## Supplementary Methods for "Accelerating iterative deconvolution and multiview fusion by orders of magnitude"

**Supplementary Video Captions**

**Supplementary Video 1, Deconvolution comparison on simulated object**. Two parallel lines in 3D space are blurred by the iSIM point spread function (1st column) and deconvolved using traditional ('Trad', 2nd column) and Wiener-Butterworth ('WB', 3rd column) back projectors. For clarity, only a transverse XY plane through the object is shown. Corresponding line profiles through the middle of the image (4th column) are also shown, updated after each iteration count. **See also Fig. 1e**.

**Supplementary Video 2, Deconvolution comparison on fixed U2OS cells, immunolabeled with Tomm20 Alexa-488**. Fixed U2OS cells were immunolabeled with Tomm 20 Alexa-488 and imaged with iSIM. Single planes (top) and higher magnification views (bottom, corresponding to the yellow rectangular region) are shown, derived from image volumes deconvolved using traditional (left) and Wiener-Butterworth (right) back projectors, with iteration number (it) indicated. **See also Fig. 1f, g**.

**Supplementary Video 3, Time-lapse images of U2OS cells expressing ERmoxGFP**. Image volumes were acquired with iSIM every 2 s, over 150 time points, and deconvolved (1 iteration) using the Wiener-Butterworth back projector. The deconvolved maximum intensity projection is shown here.

**Supplementary Video 4, Angular maximum intensity projections, comparing raw and deconvolved *C. elegans* embryos showing histone and neuronal markers**. *C. elegans* embryos expressing neuronal (green, GFP-membrane) and pan-nuclear (magenta, mCherry-histone) markers were acquired with diSPIM (top, raw views) and deconvolved using traditional ('Trad', bottom left) and Wiener-Butterworth ('WB', bottom right) back projectors. Isotropic reconstructions are obtained with both back projectors. **See also Fig. 2a-d**.

**Supplementary Video 5, Two-color time-lapse images of *C. elegans* embryos expressing histone and neuronal markers**. *C. elegans* embryos expressing neuronal (green, GFP-membrane) and pan-nuclear (magenta, mCherry-histone) markers were acquired with diSPIM every 1.5 min, over 50 time points, and deconvolved using traditional and Wiener-Butterworth back projectors. Maximum intensity projections of raw (View A, left column) and deconvolutions using traditional ('Trad', middle column) and Wiener-Butterworth ('WB', right) back projectors are shown for lateral (top) and axial (bottom) views. Time is referenced as minutes post fertilization. **See also Fig. 2a-d**.

**Supplementary Video 6, Time-lapse images of Jurkat T cells expressing GFP-actin**. Jurkat T cells expressing GFP-actin were acquired with a quadruple-view light-sheet microscope every 15 s, over 30 time points, and deconvolved using traditional and Wiener-Butterworth back projectors. Left: Perspective 3D view of Wiener-Butterworth deconvolution result . Right: Maximum intensity projections of raw (top row) and deconvolved results using traditional ('Trad', middle row) and Wiener-Butterworth ('WB', bottom row) back projectors are shown for lateral (left) and axial (right) views. **See also Fig. 2e-g**.

**Supplementary Video 7, Time-lapse images of zebrafish embryo expressing Lyn-eGFP**. Volumetric images of 32-hour zebrafish embryo expressing Lyn-eGFP under the control of the ClaudinB were acquired with diSPIM every 30 s, over 900 time points (a total acquisition period of 7.5 h), and deconvolved using the Wiener-Butterworth back projector. Maximum intensity projections of deconvolutions are shown for lateral (top) and axial (bottom) views. Labels that indicate anterior and posterior directions, the direction of the coverslip; and the skin cell layer are also indicated. Images were binned 2×2 relative to the data for display and memory purposes. **See also Fig. 2j**.

**Supplementary Video 8, Higher magnification view of subregion in Supplementary Video 7, highlighting the immune cell migration.** See also Fig. 2m, n.

**Supplementary Video 9, Automated segmentation of cell membranes in zebrafish embryo lateral line.** Slices indicated by z-distance from skin surface shows the automated segmentation result based on the raw data ('Raw', View A image (top)), deconvolved image with Wiener-Butterworth back projector ('WB', middle) and the overlay of the segmentations (bottom). **See also Fig. 2o, p.**

**Supplementary Video 10, Cleared tissue: mouse brain slice**. A 4 x 2 x 0.5 $mm^3$ slab of brain tissue derived from a V1b transgenic mouse was immunolabeled and cleared using iDISCO+, imaged in the cleared tissue diSPIM, and reconstructed by registering the two views and deconvolving them. An Alexa Fluor 555 secondary antibody against tdTomato primary antibody sparsely labels neurons and neurites across the entire volume. See also **Fig. 3a**.

**Supplementary Video 11, Cleared tissue: 4-color embryonic mouse intestine**. A 2.1 x 2.5 x 1.5 $mm^3$ intestinal volume from an E18.5 mouse was immunolabeled and cleared using iDISCO, imaged in the cleared tissue diSPIM, and reconstructed by registering the two views and deconvolving them. Orange: Alexa Fluor 647 secondary antibody against Tomm20 primary antibody; purple: Alexa Fluor 568 secondary antibody against $\alpha$-Tubulin primary antibody; yellow: Alexa Fluor 488 secondary antibody against PECAM-1 primary antibody; blue: DAPI, highlighting nuclei. See also **Fig. 3d**.

**Supplementary Video 12, Cleared tissue: 2-color adult mouse intestine**. A 2.3 x 0.7 x 0.5 $mm^3$ intestinal volume from an adult C57BL/6 mouse was immunolabeled and cleared using iDISCO, imaged in the cleared tissue diSPIM, and reconstructed by registering the two views and deconvolving them. Red: Alexa Fluor 488 secondary antibody against PECAM-1 primary antibody; Cyan: autofluorescence of tissue.

**Supplementary Video 13, Cleared tissue: 2-color embryonic mouse stomach**. A 2.4 x 2.7 x 1.0 $mm^3$ volume of stomach from an E18.5 mouse was immunolabeled and cleared using iDISCO, imaged in the cleared tissue diSPIM, and reconstructed by registering the two views and deconvolving them. Red: Alexa Fluor 647 secondary antibody against PECAM-1 primary antibody; Green: Alexa Fluor 568 secondary antibody against $\alpha$-Tubulin primary antibody.

**Supplementary Video 14, Cleared tissue: 2-color adult mouse ovary**. A 2.6 x 1.9 x 0.5 $mm^3$ volume ovary from an adult C57BL/6 mouse was immunolabeled and cleared using iDISCO, imaged in the cleared tissue diSPIM, and reconstructed by registering the two views and deconvolving them. Green: Alexa Fluor 488 secondary antibody against CD11c primary antibody; Red: CF-568 secondary antibody against CD11b primary antibody.

**Supplementary Video 15, Three-fold *C. elegans* embryos expressing GCaMP3 from a *myo-3* promoter** were imaged in the reflective diSPIM on mirrored coverslips (155 volumes, each acquired every 350 ms). Maximum intensity projections of raw data (left column), traditional (Trad) joint deconvolution (second column), Wiener-Butterworth deconvolution (third column), and deep learning reconstruction (right column, 100 time points are used for training, and the other 55 time points are used for validation) are shown for lateral (top) and axial (bottom) views. See also **Fig. 4c**.

**Supplementary Video 16, U2OS cells expressing mEmerald-$\alpha$-Actinin** were imaged in the reflective LLS microscope on mirrored coverslips (100 volumes, each acquired every 2.5 s). Maximum intensity projections of raw data (left column), Wiener-Butterworth deconvolution (middle column), and deep learning reconstruction (right column, 80 time points are used for training, and the other 20 time points are used for validation) are shown for lateral (top) and axial (bottom) views. **See also Fig. 4f**.

**Supplementary Video 17, *C. elegans* embryos with GFP-histone label** were imaged with a free-space coupled diSPIM on a glass coverslip. Maximum intensity projections of raw view A (first column), raw view B (second column), traditional (Trad) joint deconvolution of view A and view B (third column), deep learning reconstruction (fourth column) based on single-view input (i.e., view A) and deep learning reconstruction (rightmost column) based on dual-inputs (i.e., both view A and view B) are shown for lateral (top) and axial (bottom) views. In the deep learning reconstructions, 180 time points are used for training, and the other 110 time points are used for validation. hpf: hours post fertilization. See also **Fig. S5** in **Supplementary Note 2**.

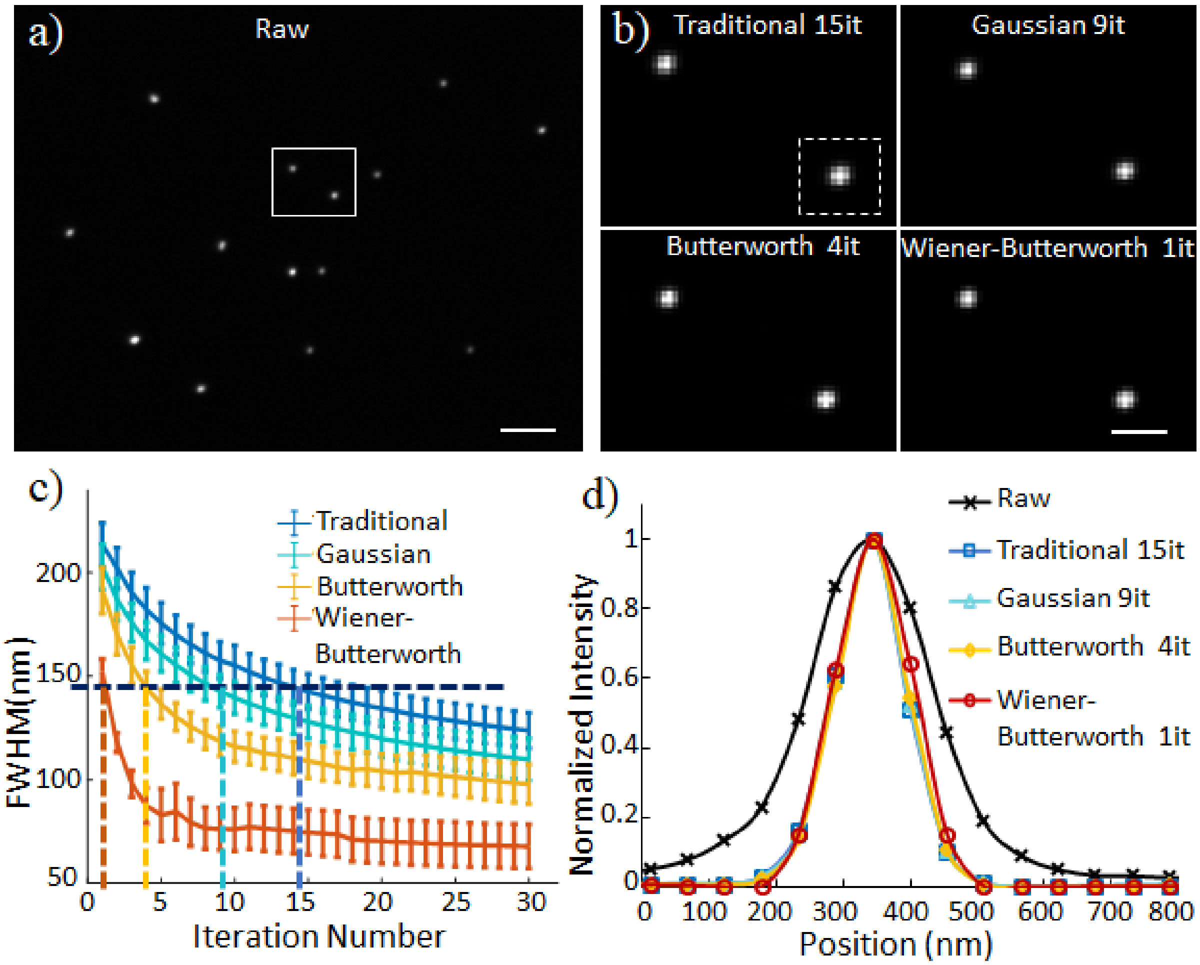


**Supplementary Fig. 1, Comparing different back projectors when deconvolving 100 nm beads captured with the iSIM. a)** 100 nm yellow green beads were imaged on the iSIM. Raw data are shown (single planes from imaging volumes). **b)** Deconvolved, higher magnification views of the white rectangle in **a)** with indicated back projector. **c)** Full width at half maximum (FWHM) values derived from 10 beads as shown in **Fig. 1**, standard deviations as well as mean values are shown. Note the considerably more rapid descent to the resolution-limited result of the Wiener-Butterworth back projector compared to other back projectors. The resolution limit of iSIM is indicated with the horizontal dotted line. **d**) Line profiles through the example bead highlighted by the dotted white rectangle in **b**), showing that all back projectors yield an equivalent estimate of the bead. Scale bars: 2 μm in **a)** and 500 nm in **b)**.

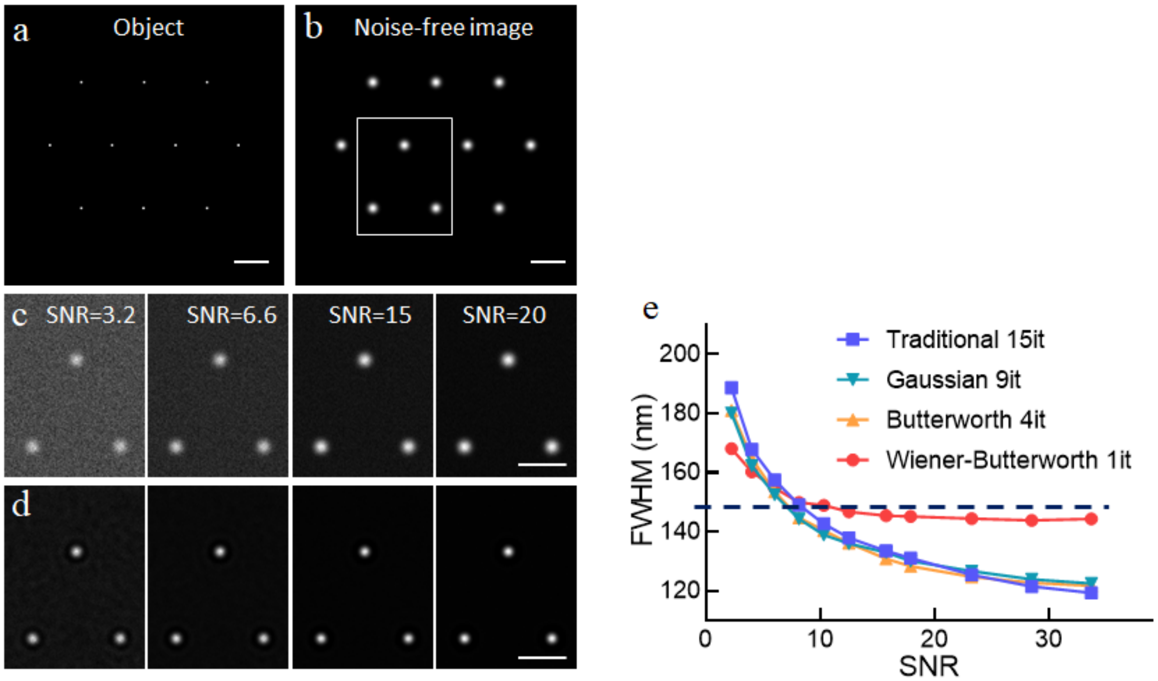


**Supplementary Fig. 2, Simulated resolution vs. input data SNR curves for different back projectors. a)** Ten synthetic point sources form the object. **b)** Blurring the object with the iSIM PSF generates a noise-free image. **c)** Gaussian and Poisson noise are added to the noise-free image to generate input images of variable SNR. See **Methods** for further detail on this procedure. **d)** Images in **c**) after 1 iteration deconvolution with Wiener-Butterworth filter. **e)** Average bead FWHM values vs. input data SNR are plotted for different back projectors. The Wiener-Butterworth filter provides more accurate (i.e. closer to the diffraction-limit) resolution enhancement than do other choices over a wide range of SNRs.

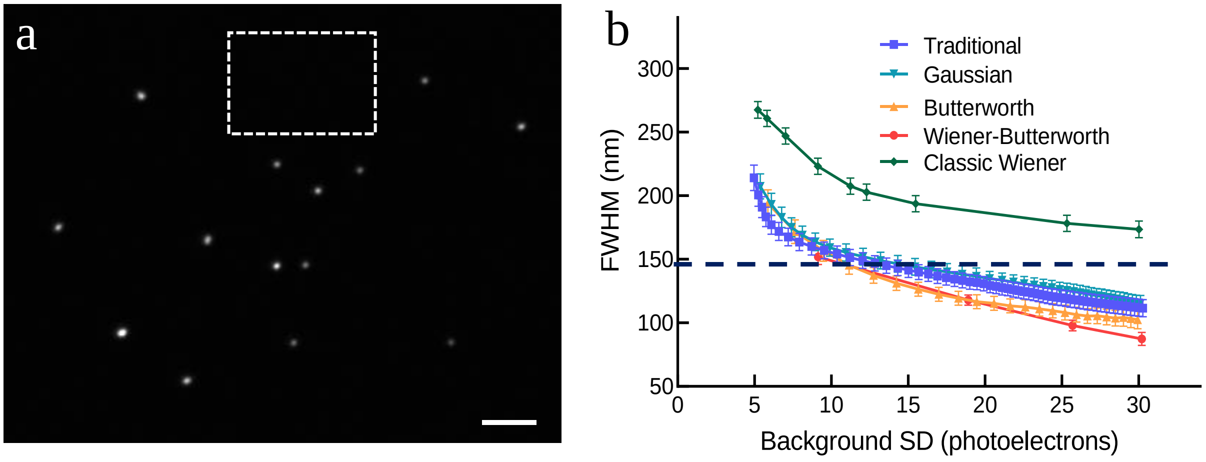


**Supplementary Figure 3, Resolution vs. background noise amplification for different back projectors. a)** Raw bead data in **Supplementary Figure 1** were deconvolved with different back projectors **b)** and the FHWM of 10 beads plotted as a function of the standard deviation of the background (corresponding to the dotted rectangular region in **a**) to assess noise-resolution tradeoffs achievable by the various deconvolution methods. For the RLD methods, different noise-resolution combinations were generated by increasing the number of iterations, which both amplifies noise and improves resolution. For the classic Wiener filter, different noise-resolution combinations were generated by changing the filter parameter. Curves closer to the lower left origin of the plot are better as they indicate the ability to achieve a specified resolution with lower noise amplification. Deconvolution with the Wiener-Butterworth (WB) back projector offers consistently comparable or lower noise amplification at any given resolution than other approaches, i.e., despite its fast descent to the resolution limit, WB noise amplification is no worse than with other methods. The horizontal dotted line indicates the resolution limit. Scale bar in **a**) 2 µm.

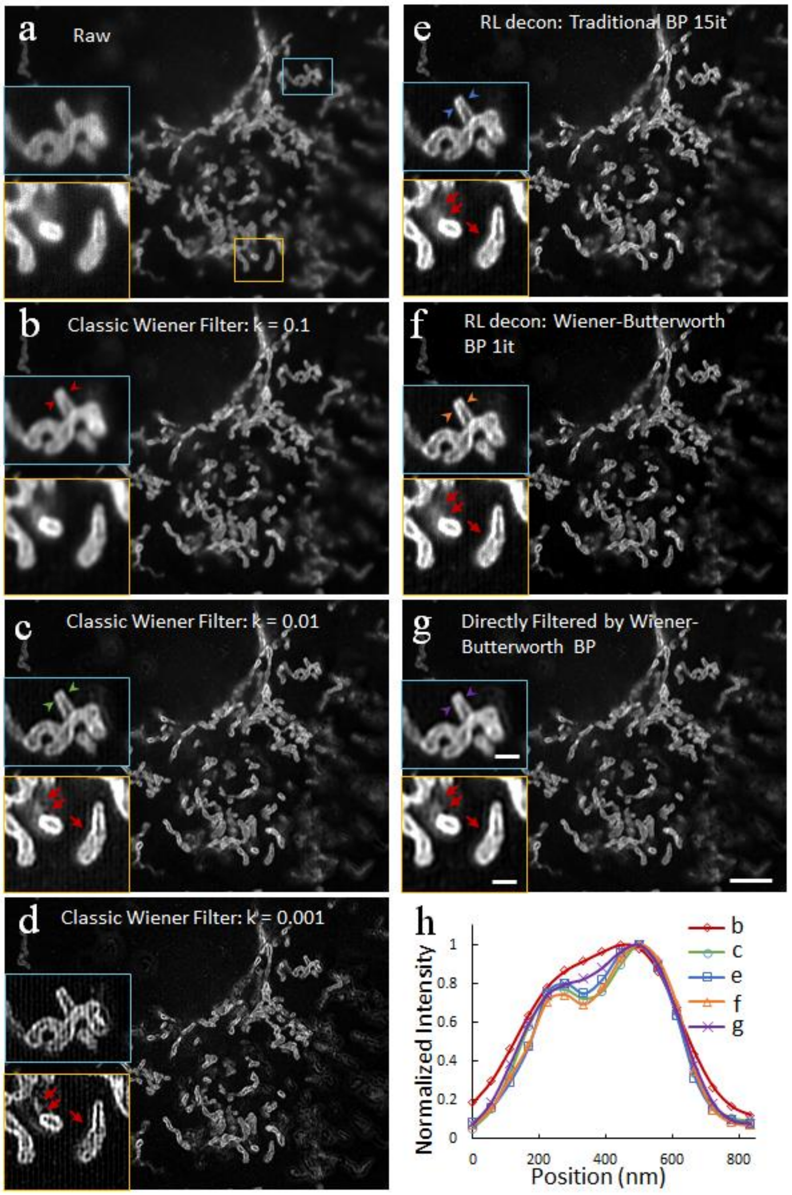


**Supplementary Fig. 4, Comparing the Wiener Filter to Richardson Lucy Deconvolution. a)** Fixed U2OS cells immunolabeled against Tomm20 and imaged with iSIM. Dataset is the same as in **Fig. 1f**. **b-d**) Data deconvolved using Wiener filter with k value as indicated. **e, f**) Richardson Lucy Deconvolution with conventional (**e**) and Wiener-Butterworth (**f**) back projectors with indicated iteration (it) numbers. **g)** Wiener deconvolution using the Wiener-Butterworth back projector instead of the typical kernel. Blue and orange rectangular insets in **a-g**) show higher magnification views. Line profiles across the mitochondrial structure marked with the arrowheads in **b, c, e-g** are shown in **h**). At k = 0.1 (**b**), the Wiener filter fails to resolve the boundaries of the mitochondria. Smaller values of k better resolve the mitochondria (**h**) but lead to progressively greater ringing artifacts (red arrows in **c-d**). Such artifacts are also present when using the Wiener-Butterworth filter in Wiener deconvolution, which also fails to fully resolve the mitochondrial boundaries. Richardson Lucy Deconvolution (**e, f**) resolve the mitochondria without introducing ringing. Scale bars: 5 μm in lower magnification view and 1 μm in inset.

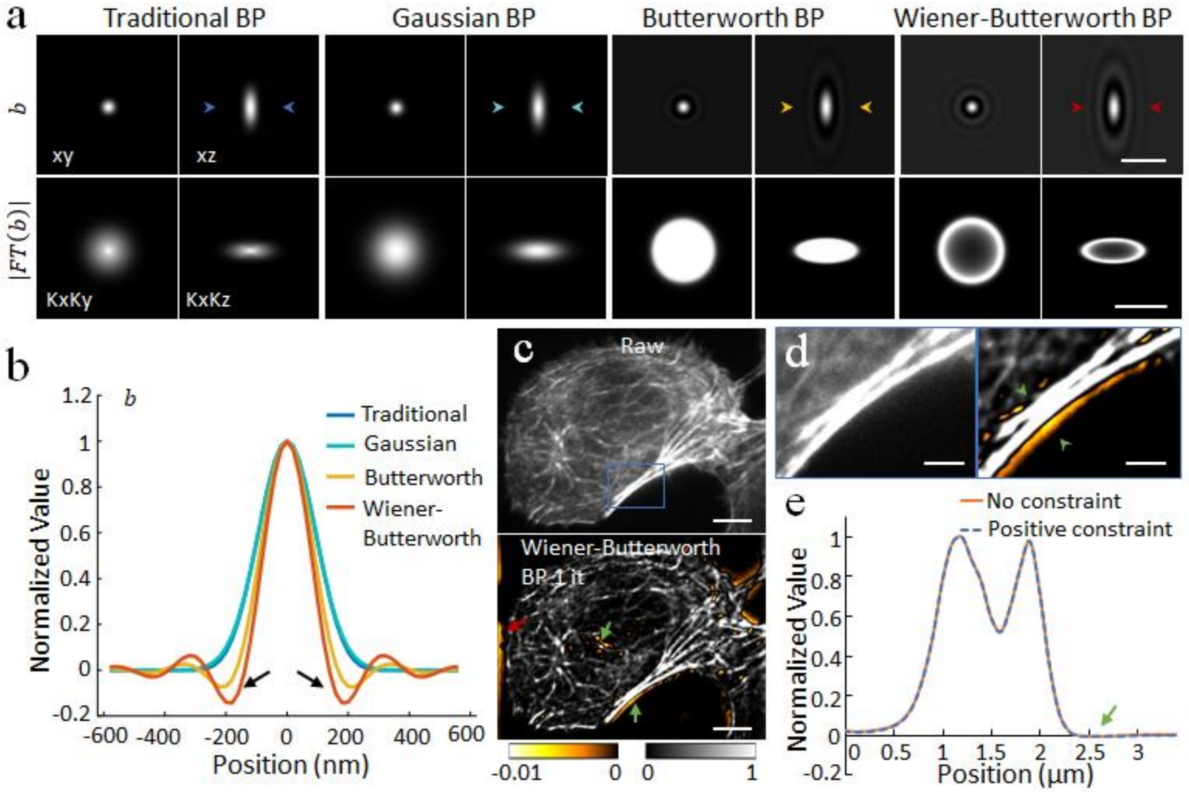


**Supplementary Fig. 5, Negative value analysis. a)** Different back projectors in real space (top row) and Fourier space (bottom row). See also **Fig. 1b. b)** Line profiles through regions demarcated by arrowheads in **a**)**.** Butterworth and Wiener-Butterworth back projectors exhibit negative values, highlighted by black arrows. **c)** Alexa Fluor 488 Phalloidin-labeled actin in fixed U2OS cell, as in **Supplementary Fig. 3**. Raw data (top) and Richardson Lucy deconvolution with Wiener-Butterworth back projector prior to negative value removal, 1 iteration (bottom) are shown. Negative values are highlighted with arrows and with yellow colormap. **d)** Higher magnification of blue rectangular region in **c**), showing raw data (left) and deconvolved result prior to negative value removal (right). Note that negative values mostly arise in background pixels, or in very low intensity pixels adjacent to regions of higher intensity. **e**) Line profile through region demarcated by arrowheads in **d**, comparing the effect of setting negative valued pixels to zero (positivity constraint, blue dotted line) to the deconvolved data without constraint (orange solid line). Profiles overlay except for very slight differences near zero on right-hand portion of curve (arrow). Scale bars: 1 μm (top) and 1/100 nm$^{-1}$ (bottom) in **a)**, 5 μm in **c)**, and 2 μm in **d**).

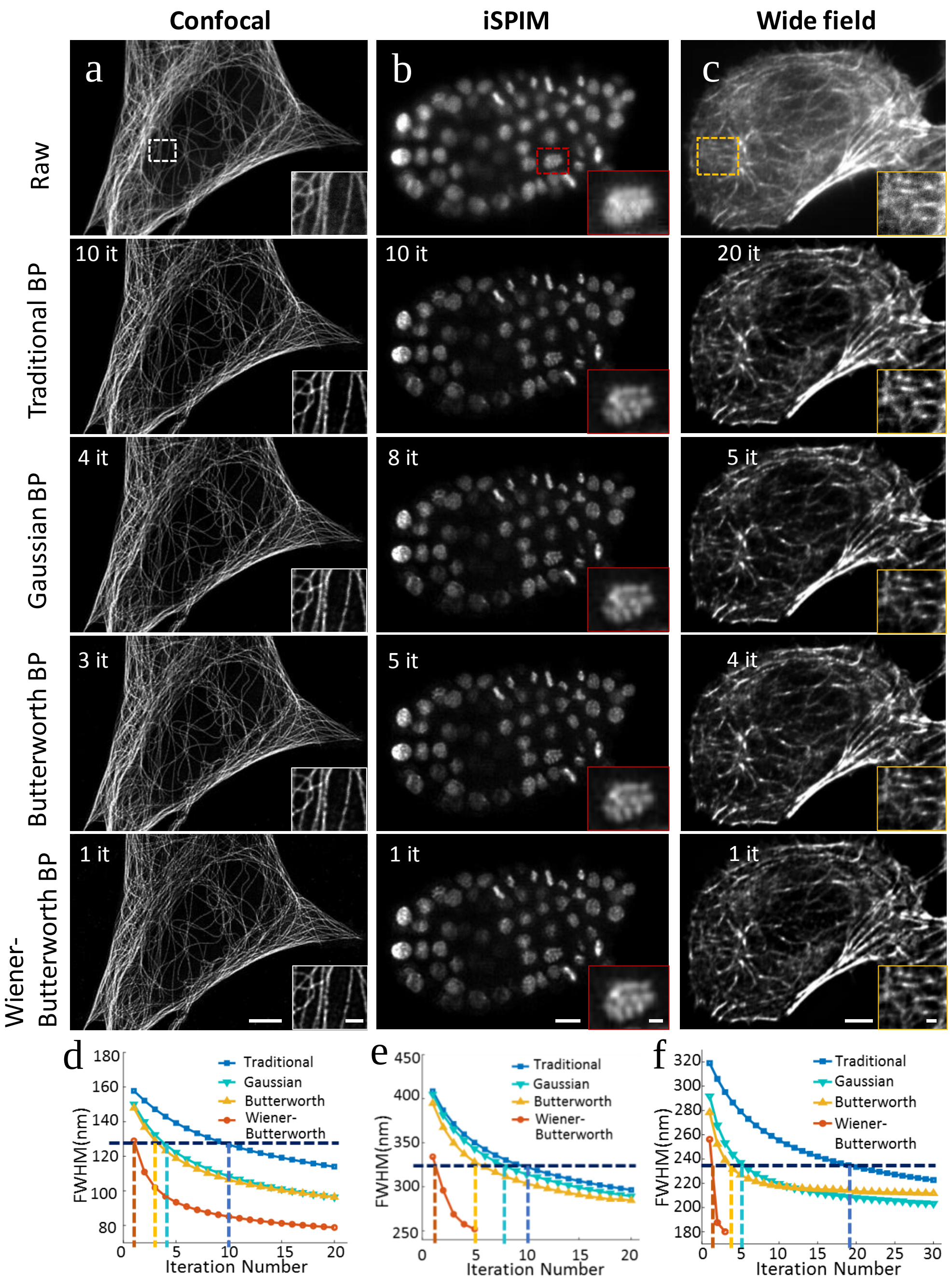


**Supplementary Fig. 6, Unmatched back projectors speed iterative deconvolution convergence in confocal, light-sheet, and widefield microscopy.** Confocal (**a**, immunolabeled microtubules), inverted selective plane illumination (iSPIM, **b**, GFP-labeled histones) and widefield (**c**, phalloidin-labeled actin) microscopy images are shown. Note higher magnification insets at lower right corner of each image. Single planes from imaging stacks are shown, with iteration number and back projector as indicated. The effect of different back projectors is illustrated, with experimental resolution vs. iteration number curves derived by plotting the mean FWHM of 10 microtubule filaments (**d**) or beads (**e** and **f**) vs. iteration number as shown in graphs at bottom. Iteration numbers (vertical dotted lines) corresponding to the diffraction limit (horizontal dotted lines) were used in generating the images, as in **Fig. 1d, f**. Scale bar: 5 μm in main figures and 5 μm in subsets.

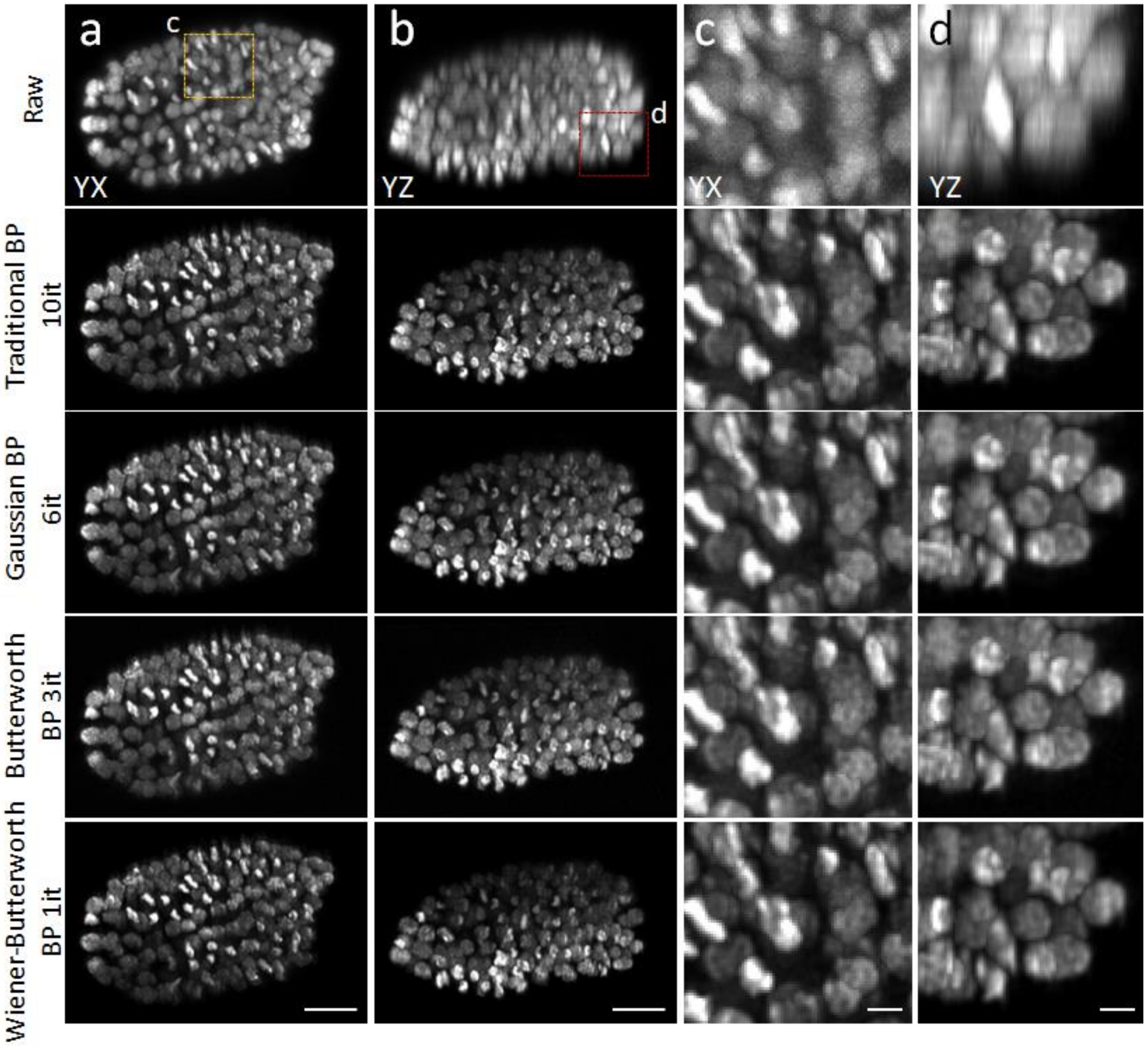


**Supplementary Fig. 7, Unmatched back projectors reduce the number of iterations required when deconvolving dual-view light-sheet microscopy (diSPIM) data.** Nematode embryos expressing GFP-labeled histones were imaged in diSPIM. Lateral (**a**) and axial (**b**) views are shown for raw and deconvolved data, with iteration ('it') numbers and back projectors as indicated. Higher magnification views **c, d**) correspond to yellow and red rectangular regions in **a, b**). Scale bars: 10 μm in **a, b**; 2 μm in **c, d**.

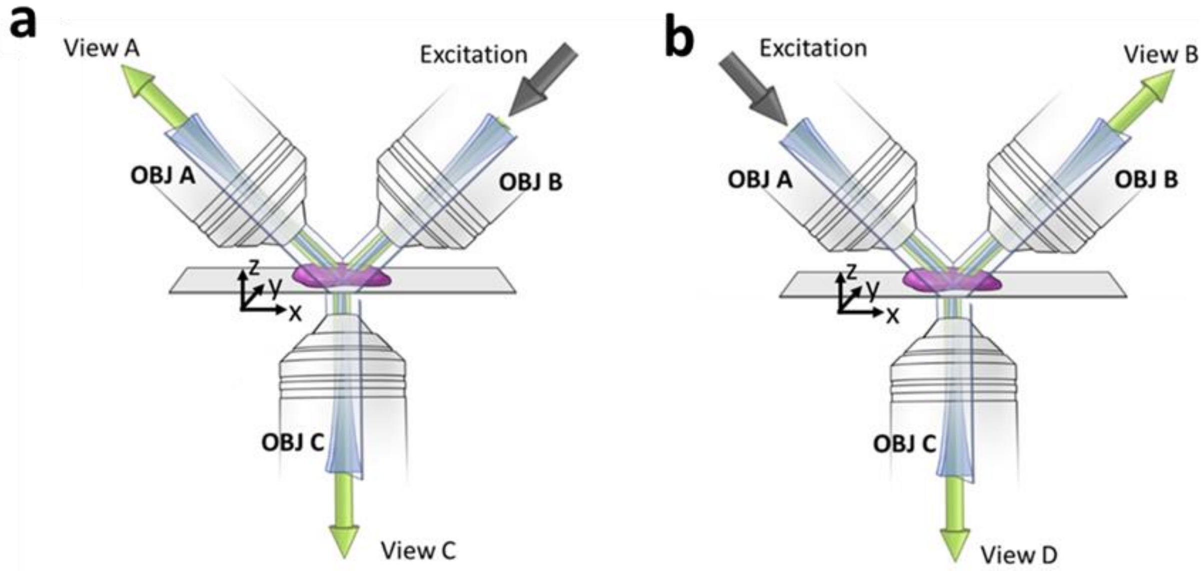


**Supplementary Fig. 8, Schematic of the quad-view SPIM system for acquiring up to 4 volumetric views.** Dual views (View A and View B) are sequentially acquired by diSPIM using the top two objectives (OBJ A and OBJ B). Four views (View A, B, C and D) are acquired with the additional third objective (OBJ C). **(a)** Views A/C are simultaneously acquired when the excitation is introduced from OBJ B; **(b)** Views B/D are simultaneously acquired when the excitation is introduced from OBJ A. Here, x, y, z coordinates are defined from the perspective of the coverslip.

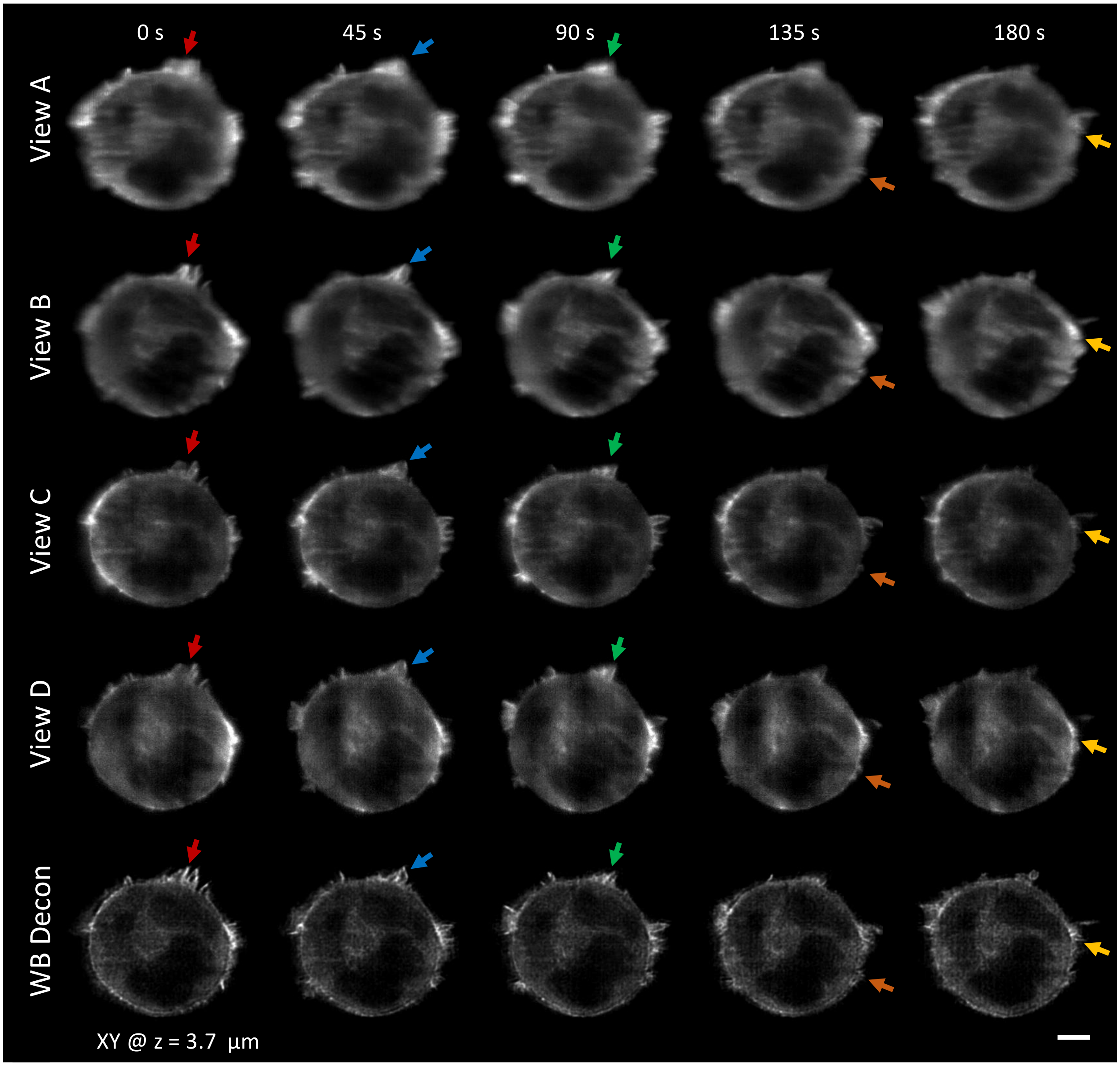


**Supplementary Fig. 9, Quad-view Wiener-Butterworth (WB) deconvolution resolves more detail than raw views.** Colored arrow pairs highlight fine, GFP-labeled actin structures expressed in Jurkat T cell for comparison. View A and View B are collected by the upper 0.8 NA collection lenses; View C and View D are collected by the lower 1.2 NA collection lens. Scale bar: 5 µm. See also **Fig. 2f**.

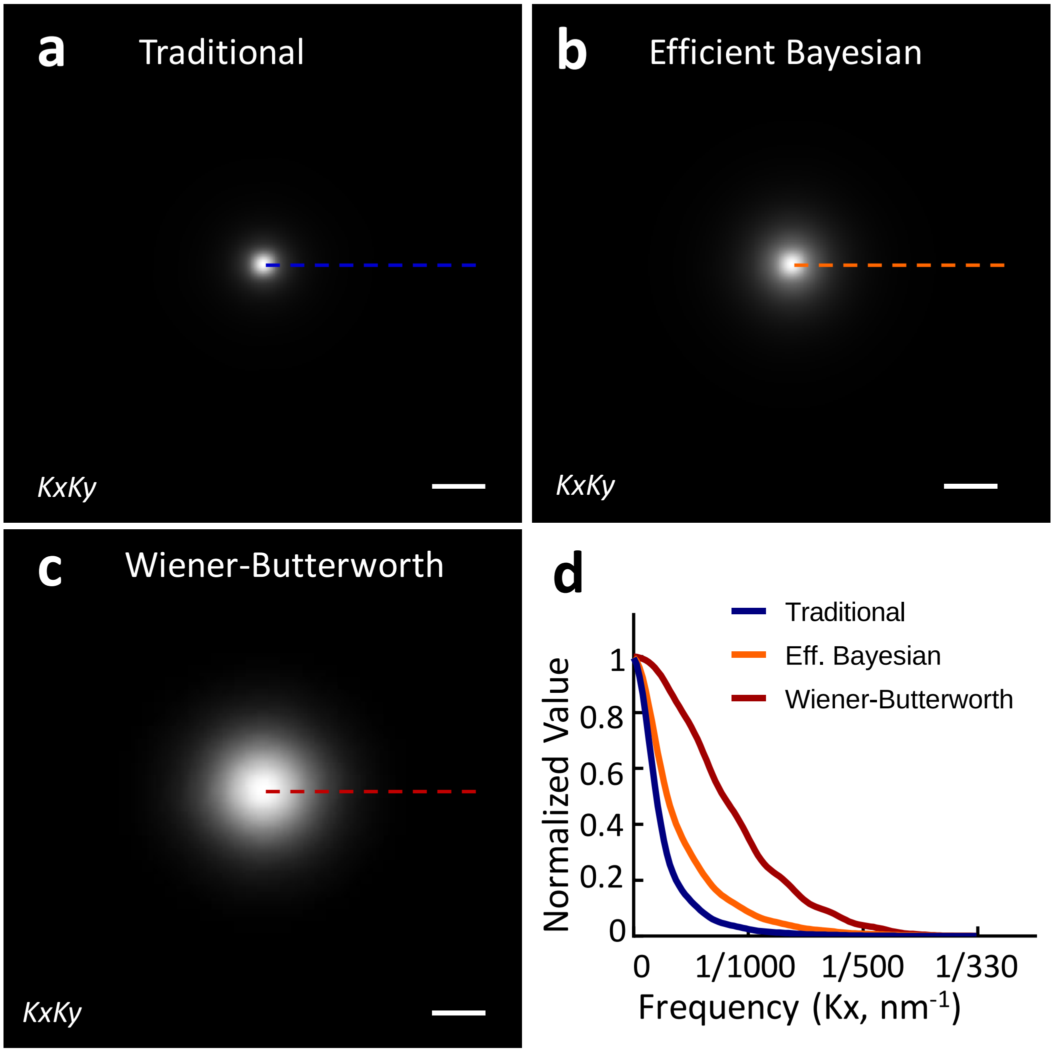


**Supplementary Fig. 10, |FT(*f*) x FT(*b*)| comparison for traditional, Efficient Bayesian and Wiener-Butterworth back projectors in Quadruple-view light-sheet microscopy.** |FT(*f*) x FT(*b*)|for the bottom objective, with traditional **(a)**, Efficient Bayesian **(b)** and Wiener-Butterworth **(c)** back projectors. **(d)** line profiles through the images in **a**, **b** and **c**. Scale bars: 1/1000 $nm^{-1}$.

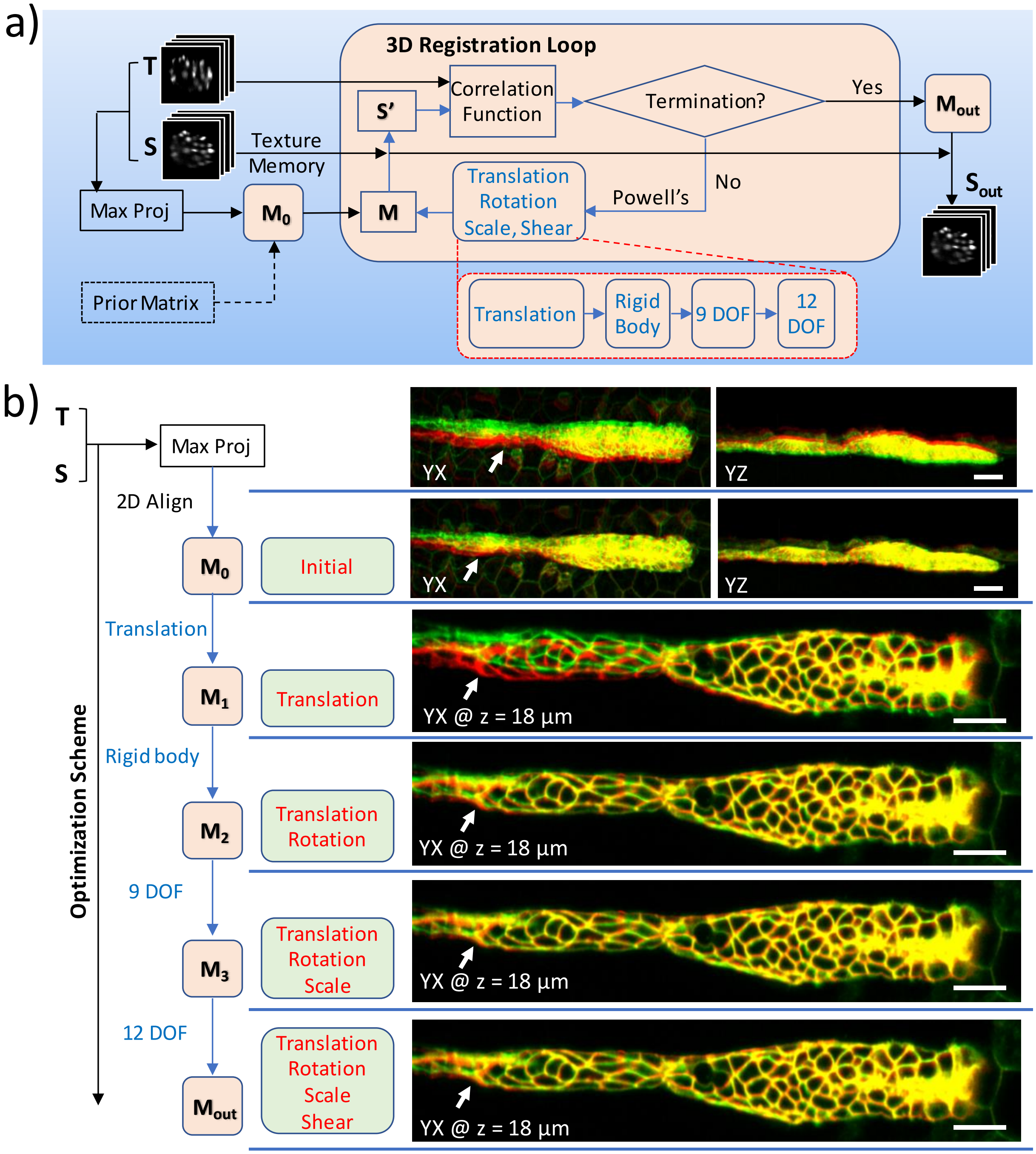


**Supplementary Fig. 11, Schematic of GPU-based 3D registration**. **a)** The inputs to the registration are usually two 3D images, referred as the source (S, image to be registered) and target image (T, fixed image). The XY and ZY maximum intensity projections of the input 3D images are used for preliminary alignment (only adjusting translation and rotation) and to generate an initial transformation matrix ($M_0$). Alternatively, a transformation matrix from a prior time point is used as $M_0$. A 3D registration loop iteratively performs affine transfomations on S (which is kept in GPU texture memory for fast trilinear interpolation) based on the transformation matrix M. The correlation ratio between the transformed source (S') and T is used as the cost function. This cost function is minimized with Powell's method, updating the transformation matrix by optimizing its four affine tranformation components: translation, rotation, scale and shear. The optimization is performed serially, optimizing translation; translation and rotation (i.e. rigid body registration with 6 degrees of freedom);translation, rotation and scale (9 degrees of freedom) ; and finally translation, rotation, scale and shear (12 degrees of freedom). **b)** Example images of zebrafish embryo expressing Lyn-eGFP (see also **Fig. 2j**) show the iteration optimization scheme and corresponding improvements in registration quality (white arrows). Target images are shown in red, source images in green, and the overlay in yellow. Images in top two rows are maximum intensity projections of lateral (left) and axial (right) views, while images in other rows are single planes from 3D stacks. See **Methods** for further details on this process. Scale bars: 20 μm for all images.

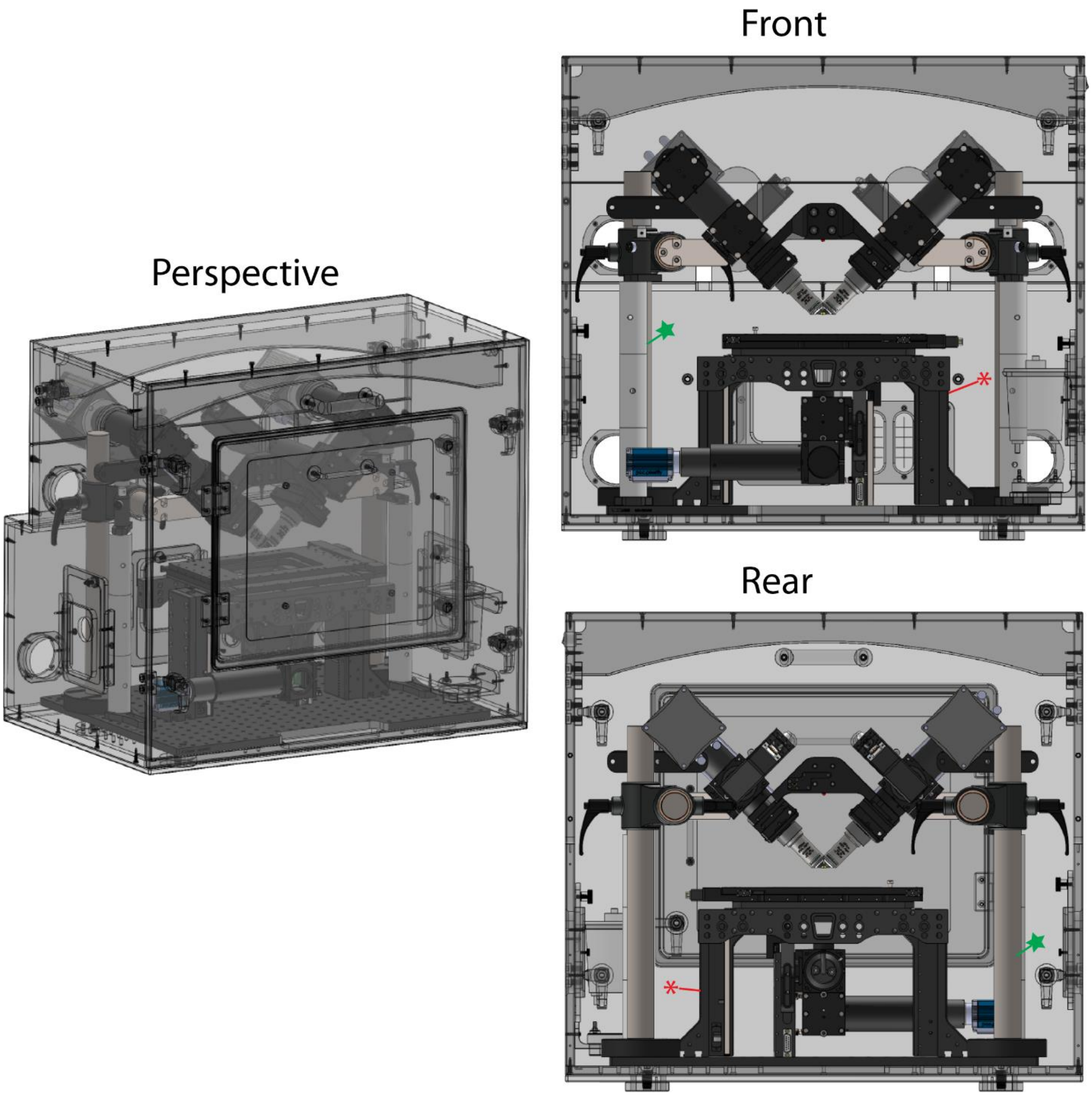


**Supplementary Fig. 12, Dispim for cleared tissue imaging**. Perspective, front, and rear drawings are shown. A substantive change from our previous diSPIM includes a movable 3D sample stage (FTP-2000, indicated with red asterisk), that allows the illumination and detection systems to remain fixed in space, improving imaging of large specimens. Additional changes include post systems (green stars) that support the fixed diSPIM head. The objectives can be easily switched between 40x, 0.8 NA water immersion lenses (shown here) for imaging live, aqueous samples and 17.9x, 0.4 NA multi-immersion lenses for cleared tissue imaging. See **Methods** for further details.

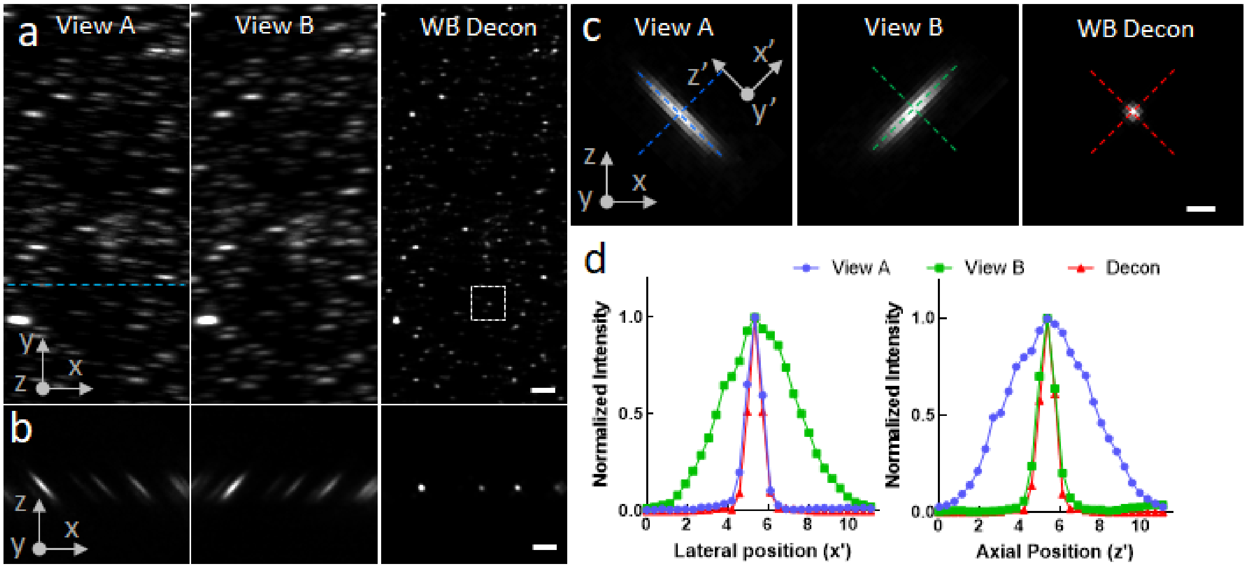


**Supplementary Fig. 13, Wiener-Butterworth deconvolution improves spatial resolution of single-view cleared tissue diSPIM.** 100 nm yellow-green fluorescent beads were deposited on a coverslip, immersed in dibenzyl ether, and imaged in the cleared-tissue diSPIM. **a)** xy maximum intensity projections corresponding to raw single views and Wiener-Butterworth (1 iteration) deconvolution of registered views. Note that data have been rotated so that x, y are parallel to the plane of the coverslip and z is normal to the coverslip. **b)** Axial plane corresponding to dotted blue line in **a)**. **c)** Higher magnification images of single beads. Primed coordinates are from perspective of light-sheet collection objective. **d)** Lateral and axial profiles as indicated in **c)**. Scale bars: **a), b)** 5 μm, **c)** 2 μm.

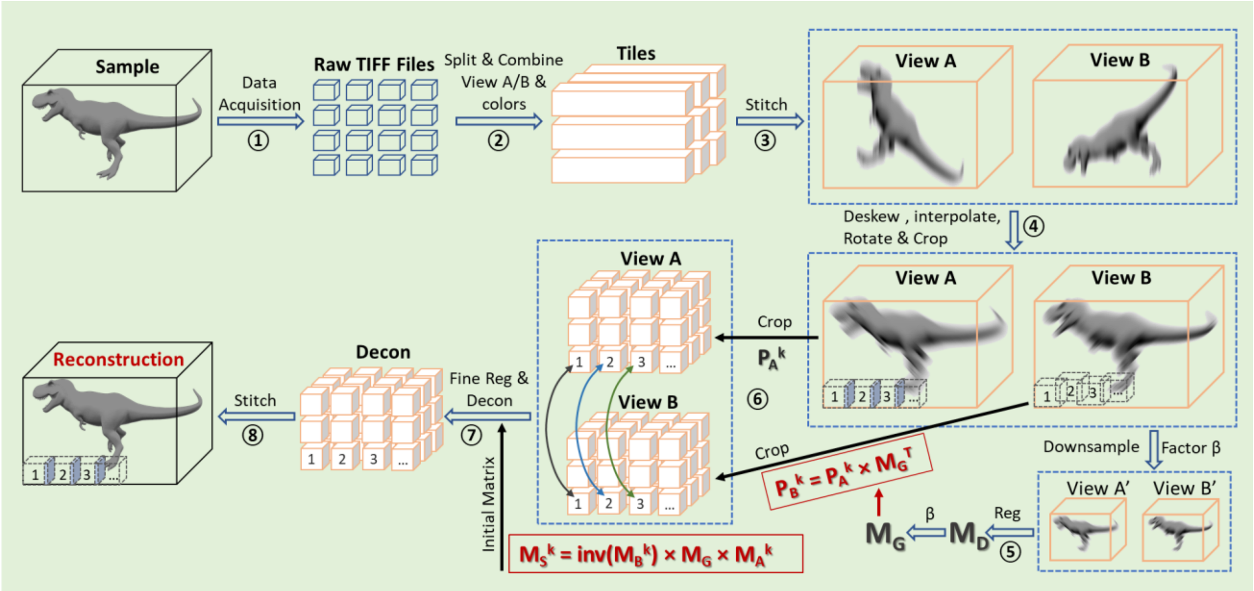


**Supplementary Fig. 14, Post-processing pipeline for large, cleared tissue data imaged with diSPIM**. Raw data acquired by the cleared-tissue diSPIM are saved as multiple 16-bit TIFF files (each less than or equal to 4 GB, step 1). These files contain mixed information, i.e. colors, views, and spatial locations are saved as they are recorded in the acquisition sequence. The XY slices are re-organized and re-saved as TIFF stacks (each usually a few hundred GB), each corresponding to a distinct spatial strip/color/view (step 2). Strips for each color/view are then combined (with Imaris Stitcher) to reassemble blurred, images of the sample (View A and View B, step 3), rotated 45 degrees relative to the coverslip. These TIFF stacks at each color and each view are deskewed (transforming from stage-scanning mode to light-sheet scanning mode), interpolated (obtaining isotropic pixel resolution), rotated (transformed from the objective view to the perspective of coverslip), cropped (saving memory), and resaved as TIFF files (e.g. ~ 2 TB for 4 colors/2 views for the dataset shown in **Fig. 3d,** step 4). By down-sampling View A and View B (typically by a factor of 5) to View A' and View B' and registering them, a coarse, global transformation matrix $\mathbf{M_G}$ that maps view B to view A can be calculated based on the registration matrix $\mathbf{M_D}$ that maps view B' to view A' (step 5). The coarsely registered View A and View B are then split into multiple subvolumes (e.g., ~1000 subvolumes, each 640 x 640 x 640 pixels, step 6). Coordinates that define the cropping locations in View B (i.e., $\mathbf{P_B}^k$) can be roughly estimated from the position in the cropped View A (i.e., $\mathbf{P_A}^k$) and the transpose of the coarse, global transformation matrix $\mathbf{M_G}$ (i.e., $\mathbf{M_G}^T$). The index k denotes the index of the subvolume and runs from 1 to the total number of subvolumes. Each of the cropped tiles in View B can be coarsely registered to the corresponding cropped tile in view A with a new matrix $\mathbf{M_S}^k$, derived from the cropping positions ($\mathbf{M_A}^k$, $\mathbf{M_B}^k$) and global transformation matrix $\mathbf{M_G}$. Fine registration and joint Wiener-Butterworth deconvolution are then performed on the paired subvolumes of View A and View B (step 7). Finally, stitching all deconvolved subvolumes results in the final reconstruction (step 8). See **Methods** for further information.

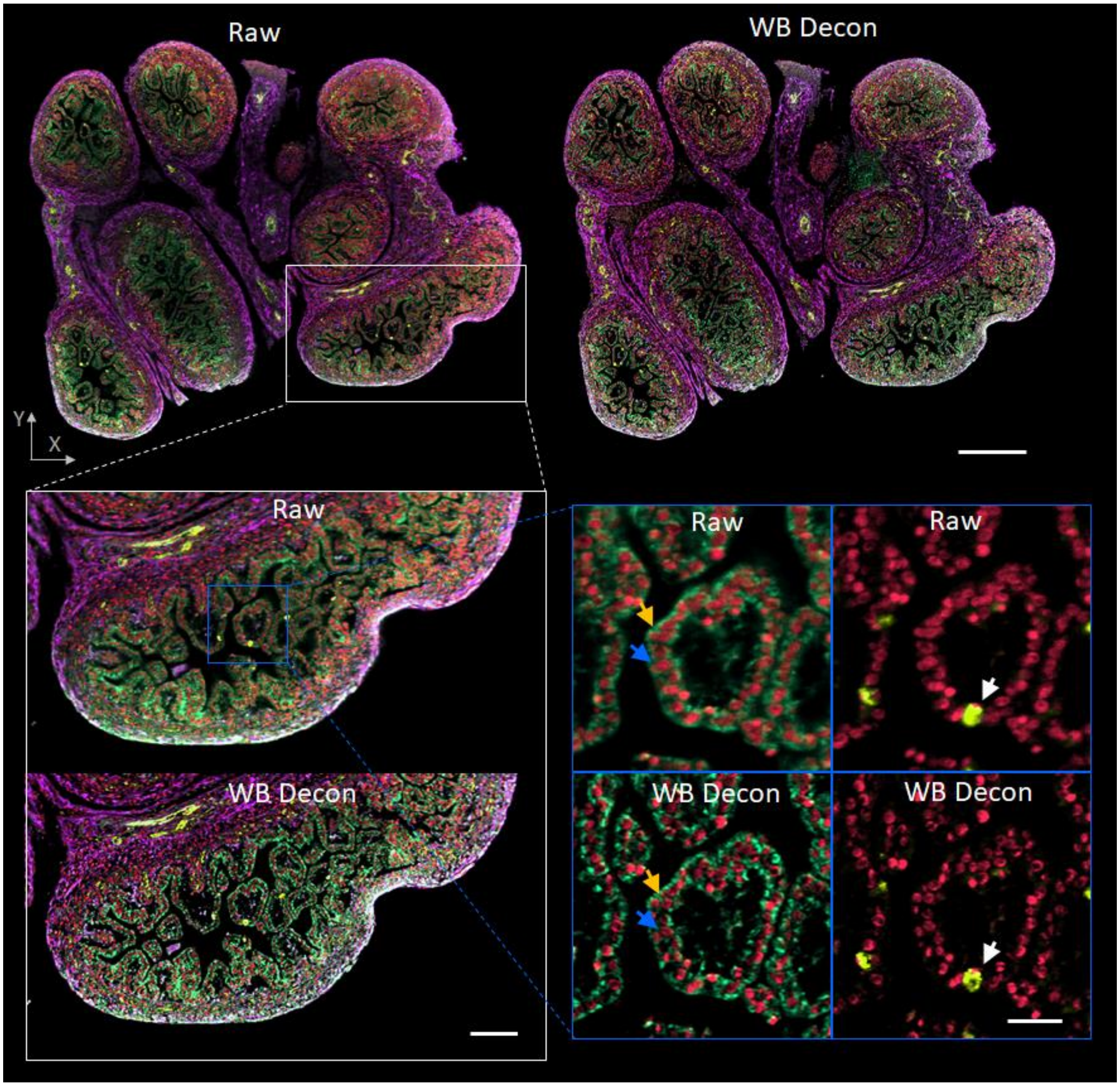


**Supplementary Fig. 15, Dual-view Wiener-Butterworth (WB) deconvolution reveals more detail than single-view imaging.** The single plane shown in **Fig. 3d** is reproduced, comparing raw single-view data versus WB deconvolution, at progressively higher levels of magnification. The increased detail evident in the deconvolved results are especially evident at the highest level of magnification, where nuclei (blue, yellow arrows) and the hollow interior of blood vessels (white arrows) are revealed in the WB deconvolution but not in raw data. Scale bars are at progressive levels of magnification: 300 μm, 100 μm, and 30 μm.

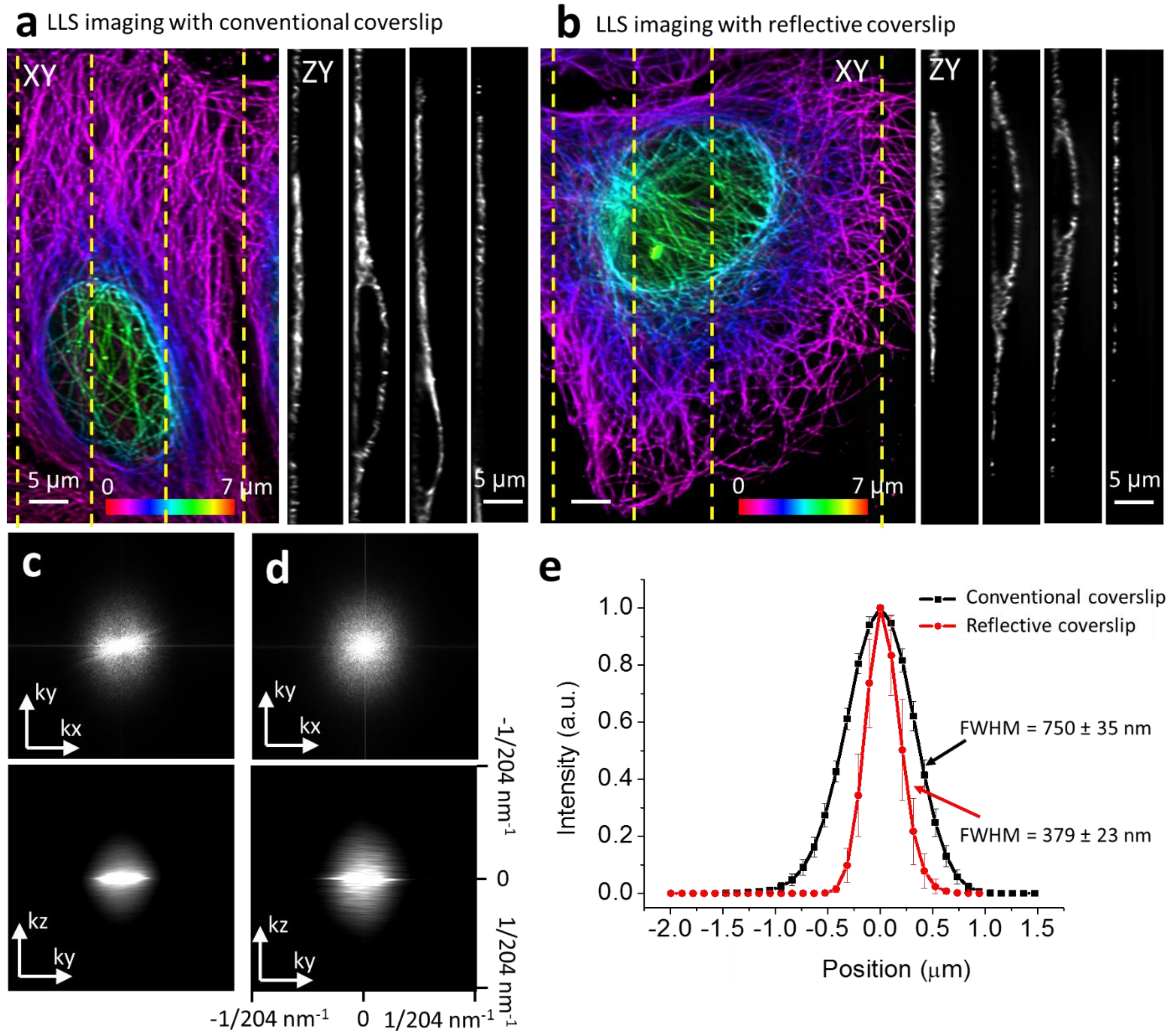


**Supplementary Figure 16, Reflective lattice light-sheet (LLS) imaging improves axial resolution.** Lateral maximum intensity projections (axial depth from coverslip indicated with color bar) derived from 3D LLS microscopy of Alexa Fluor 488 immunolabeled microtubules in fixed U2OS cells on **(a)** conventional glass and **(b)** reflective coverslip. Reconstructions were performed using traditional deconvolution with a spatially varying PSF. Axial slices (right columns) correspond to the yellow dotted lines in each panel. **(c-d)** Lateral and axial optical transfer functions (OTFs) of the images shown in a) and b), respectively. Note that the displayed OTFs were computed by averaging the 2D OTFs over all slices in the stack. Coordinates are defined from the perspective of coverslip (i.e. z is normal to the coverslip surface, xy are parallel to the coverslip surface). **(e)** The axial profiles of 10 microtubule filaments. Means and standard deviations (bars) in the plots are shown.

**Supplementary Table 1, MSE and SSIM evaluation for deconvolution results based on Gaussian, Butterworth and Wiener-Butterworth back projectors.**

| Samples | | U2OS cells Mitochondria | U2OS cells Microtubule | *C. elegans* embryo nuclei | U2OS cells actin | *C. elegans* embryo nuclei |
|---|---|---|---|---|---|---|
| **Figures/Videos** | | Fig. 1 f | Sup. Fig. 6a | Sup. Fig. 6b | Sup. Fig. 6c | Sup. Fig. 7 |
| **Microscope** | | iSIM | Confocal | iSPIM | Widefield | diSPIM |
| **MSE** | Gaussian | 5.71e-6 | 4.80e-5 | 4.67e-6 | 2.23e-4 | 2.40e-6 |
| | Butterworth | 3.21e-5 | 8.73e-5 | 1.30e-5 | 2.38e-4 | 1.61e-5 |
| | Wiener-Butterworth | 1.11e-4 | 7.81e-4 | 3.99e-5 | 7.68e-5 | 1.04e-4 |
| **SSIM** | Gaussian | 0.9978 | 0.9979 | 0.998 | 0.8418 | 0.999 |
| | Butterworth | 0.9883 | 0.9964 | 0.996 | 0.8342 | 0.9932 |
| | Wiener-Butterworth | 0.9629 | 0.9594 | 0.9889 | 0.9447 | 0.9612 |

MSE: mean square error; SSIM: structural similarity index; See **Supplementary Note 3** for detailed definitions of these quantities. All calculations are based on using the deconvolution results from the traditional RL back projector as ground truth. Note the close correspondence between ground truth and deconvolution using other back projectors, indicated by the small values of MSE and high values of SSIM.

**Supplementary Table 2. Data acquisition and processing details for data shown in Figs. 1, 2, and associated Supplementary Figures**

| Samples | | Fixed U2OS mitochondria | Live U2OS ER | Fixed U2OS microtubule | Fixed U2OS actin | *C. elegans* embryo nuclei | | *C. elegans* embryo neuron/nuclei | Jurkat T cell actin | Zebrafish embryo |
|---|---|---|---|---|---|---|---|---|---|---|
| **Figures/Videos** | | Fig. 1 e,f Sup. Fig. 4 | Sup. Video 3 | Sup. Fig. 6a | Sup. Fig. 5c, 6c | Sup. Fig 6b | Sup. Fig 7 | Fig. 2 a, b, c, d Sup. Video 4, 5 | Fig. 2 e, f, g Sup. Video 6 | Fig. 2 j-p Sup. Video 7-9 |
| **Fluorescence Label** | | Alexa-488 | ERmoxGFP | Microtubule-561 | Alexa-488 | GFP-histone | | GFP-membrane mCherry-histone | GFP-Actin | Lyn-eGFP |
| **Microscope** | | iSIM | iSIM | Confocal | Widefield | diSPIM (iSPIM) | | diSPIM | quad-view light-sheet | diSPIM |
| **View number** | | 1 | 1 | 1 | 1 | 1 | 2 | 2 | 4 | 2 |
| **Color number** | | 1 | 1 | 1 | 1 | 1 | 1 | 2 | 1 | 1 |
| **Acquisition** | Excitation | 488 | 488 | 561 | XT 640-W | 488 | | 488, 561 | 488 | 488 |
| | Step size x Slices per view per color | 0.1 µm x 95 slices | 0.5 µm x 6 slices | 0.5 µm x 6 slices | 0.15 µm x 77 slices | 1 µm x 40 slices | | 1 µm x 50 slices | 1 µm x 60 slices | 1 µm x 80 slices |
| | Acquisition time / tp | 5 s | 0.3 s | 332 s | 1.8 s | 0.5 s | | 1 s | 3.2 s | 3.5 s |
| | Time interval | -- | 2 s | -- | -- | 60 s | | 100 s | 15 s | 30 s |
| | Total time points | 1 | 150 | 1 | 1 | 780 | | 50 | 30 | 902 |
| | Total acquisition time | 5 s | 300 s | 332 s | 1.8 s | 780 min | | 83 min | 450 s | 7.5 h |
| **Image size (each volume after interpolation)** | | 1920 x 1550 x 95 | 1920 x 1550 x 6 | 1024 x 1024 x 22 | 512 x 512 x 77 | 240 x 360 x 246 | | 280 x 380 x 308 | 360 x 338 x 181 | 640 x 2048 x 496 |
| **Total data size** | | 270 M voxels, 539 MB, 16 bit | 2.5 G voxels, 5.1 GB, 16 bit | 22 M voxels, 44 MB, 16 bit | 19 M voxels, 39 MB, 16 bit | 20 M voxels, 41 MB, 16 bit | 16.1 G voxels, 31.2 GB, 16 bit | 6.1 G voxels, 12.2 GB, 16 bit | 2.5 G voxels, 5 GB, 16 bit | 1.05 T voxels, 2.1 TB, 16 bit |
| **Data processing** | Registration time / tp | -- | -- | -- | -- | -- | 1.9 s | 6.26 s | 13.4 s | 16 s |
| | Deconvolution time / tp | 2.9 s | 0.21 s | 0.29 s | 0.27 s | 0.29 s | 0.11 s | 0.16 s | 0.6 s | 8.4 s |
| | File IO / tp | 10.6 s | 1.2 s | 1.2 s | 1.0 s | 1.0 s | 0.52 s | 1.38 s | 0.82 s | 6.4 s |
| | Total time / tp | 13.5 s | 1.4 s | 1.5 s | 1.3 s | 1.3 s | 2.5 s | 7.8 s | 14.8 s | 30. 8 s |
| | Total processing time | 13.5 s | 211 s | 1.5 s | 1.3 s | 1.3 s | 33 min | 390 s | 445 s | 7.7 h |

Both registration and deconvolution refer to using new methods (faster registration, Wiener-Butterworth deconvolution) with GPU implementation on an NVIDIA Quadra M6000 card. tp: time point.

**Supplementary Table 3, Sample preparation, data acquisition and processing details for all cleared tissue datasets**

| Samples | | Adult mouse brain | Embryonic mouse intestine | Adult mouse intestine | Embryonic mouse stomach | Adult mouse ovary |
|---|---|---|---|---|---|---|
| Figures/Videos in paper | | Fig. 3 a, b, c<br>Sup. Video 10 | Fig. 3 d<br>Sup. Fig 14, Sup. Video 11 | Sup. Video 12 | Sup. Video 13 | Sup. Video 14 |
| Size | Physical Size | 4 x 2 x 0.5 $mm^3$ | 2.1 x 2.5 x 1.5 $mm^3$ | 2.3 x 0.7 x 0.5 $mm^3$ | 2.4 x 2.7 x 1.0 $mm^3$ | 2.6 x 1.9 x 0.5 $mm^3$ |
| | Number of colors | 1 | 4 | 2 | 2 | 2 |
| | Digital Size, voxels (X x Y x Z x color) | 10280 x 5160 x 1400 x 1 | 5586 x 6500 x 3930 x 4 | 6221 x 2008 x 1516 x 2 | 6370 x 7226 x 2850 x 2 | 6820 x 6100 x 1460 x 2 |
| Sample Preparation | Primary antibody | Rabbit anti RFP | Mouse anti alpha-tubulin<br>Goat anti CD31 | Rabbit anti Tomm20<br>Goat anti CD31 | Mouse anti alpha-tubulin<br>Goat anti CD31 | Mouse anti CD11c (Integrin aX)<br>Rat anti CD11b |
| | Secondary antibody | Goat anti-Rabbit IgG(H+L) Alexa-555 | Donkey anti Goat Alexa-488<br>Donkey anti Mouse Alexa-568<br>Donkey anti Rabbit Alexa-647<br>DAPI | Donkey anti Goat Alexa-488 | Donkey anti Mouse Alexa-568<br>Donkey anti Goat Alexa-647 | Goat anti Mouse IgG1 Alexa-488<br>Donkey anti Rat CF-568 |
| | Chemical Reagent | TritonX, Dichloromethane (DCM), Dibenzyl Ether (DBE), Tetrahydrofuran (THF), Methanol, Hydrogen Peroxide, Krazy Glue, Glass slide | | | | |
| Acquisition | Views | 2 | 2 | 2 | 2 | 2 |
| | Excitation Lasers | 488 | 405, 488, 561,637 | 405, 488 | 561, 637 | 488, 561 |
| | Camera ROI (slice size) | 2048 x 2048 | 2048 x 2048 | 2048 x 2048 | 2048 x 2048 | 2048 x 2048 |
| | Tile number (y x z) | 3 x 1 | 3 x 3 | 1 x 1 | 4 x 2 | 3 x 1 |
| | step size x Slices (view/color/tile) | 2 µm x 1600 slices | 2 µm x 1300 slices | 1 µm x 2200 slices | 2 µm x 1200 slices | 2 µm x 1300 slices |
| | exposure time / slice | 5 ms | 10 ms | 5 ms | 10 ms | 10 ms |
| | acquisition time / tile | 51 s | 239 s | 220 s | 104 s | 109 s |
| | Total acquisition Time (Step 1) | 154 s | 36 min | 220 s | 832 s | 327 s |

| | | | | | | | |
|---|---|---|---|---|---|---|---|
| **Post processing and time cost** | Step 2 | File I/O, splitting | 0.5 h | 2.5 h | 0.3 h | 1.2 h | 0.8 h |
| | Step 3 | File converting | 1.6 h | 4.8 h | 0.8 h | 2.4 h | 2 h |
| | | Stitching | 0.2 h | 0.8 h | 0.4 h | 0.4 h | 0.4 h |
| | | File I/O and converting | 2.5 h | 30 h | 1.3 h | 14 h | 8.1 h |
| | Step 4 | File I/O, deskew, interpolation, rotation | 4 h | 24 h | 2.2 h | 11 h | 6.6 h |
| | Step 5 | Coarse reg | 0.1 h | 0.2 h | 0.1 h | 0.2 h | 0.2 h |
| | Step 6 | Cropping subvolumes | 3.5 h | 24 h | 1.4 h | 12 h | 5.4 h |
| | Step 7 | Fine Reg / Decon | 3.4 h | 24 h | 1.4 h | 12 h | 5.3 h |
| | Step 8 | Stitching | 5 h | 30 h | 2.8 h | 12 h | 10 h |
| | Total time | | 21 h | 140 h | 10 h | 65 h | 39 h |

Overlap was in the 10-20% range in y and z for all tiling experiments, where y is defined as the lateral direction that points towards the front of the microscope and z is defined perpendicular to the coverslip. The total acquisition time is computed by multiplying the acquisition time cost for each tile by the number of tiles. This omits the time cost for moving between tiles. Since moving between tiles was done manually, we estimate an additional time cost of ~30 s per tile. This timing is negligible compared to the total acquisition time we report, and we note that it could be significantly reduced by using the microscope control software (μManager) to perform multi-tile acquisition, a capability that exists within the software.

**Supplementary Table 4, Antibodies and reagents for clear tissue preparation**

| **Primary antibody** | | | |
|---|---|---|---|
| **Name** | **Vendor** | **Catalog** | **Related Samples** |
| Rabbit anti RFP | Rockland | 600-401-379 | adult mouse brain |
| Mouse anti alpha-tubulin | Thermo Fisher Scientific | 322500 | embryonic mouse stomach, embryonic mouse intestine |
| Rabbit anti Tomm20 | Abcam | ab78547 | adult mouse intestine |
| Goat anti CD31 | R&D Systems | AF3628 | embryonic mouse stomach, embryonic mouse intestine, adult mouse intestine |
| Mouse anti CD11c (Integrin αX) | Santa Cruz | sc-398708 | adult mouse ovary |
| Rat anti CD11b | R&D Systems | MAB1124 | adult mouse ovary |
| **Secondary antibody/counter stain** | | | |
| **Name** | **Vendor** | **Catalog** | **Related Samples** |
| Goat anti Rabbit IgG (H+L) Alexa-555 | Invitrogen | A27039 | adult mouse brain |
| Donkey anti Goat IgG (H+L) AffiniPure $F(ab')_2$ Fragment Alexa-488 | Jackson ImmunoResearch | 705-546-147 | embryonic mouse intestine, adult mouse intestine |
| Goat anti Mouse IgG1 Alexa-488 | Thermo Fisher Scientific | A21121 | adult mouse ovary |
| Donkey anti Rat IgG (H+L) CF-568 | Sigma | SAB-4600077 | adult mouse ovary |
| Donkey anti Mouse IgG (H+L) Alexa-568 | Thermo Fisher Scientific | A10037 | embryonic mouse stomach, embryonic mouse intestine |
| Donkey anti Rabbit IgG (H+L) AffiniPure $F(ab')_2$ Alexa-647 | Jackson ImmunoResearch | 711-606-152 | embryonic mouse intestine |
| Donkey anti Goat IgG(H+L) Alexa-Plus-647 | Thermo Fisher Scientific | A32849 | embryonic mouse stomach |
| DAPI | Thermo Fisher Scientific | D1306 | embryonic mouse intestine |
| **Chemical Reagent** | | | |
| **Name** | **Vendor** | **Catalog** | **Purpose** |
| TritonX | Sigma | T9284 | stain/wash buffer |
| Dichloromethane (DCM) | Sigma | 270997 | Delipidation |
| Dibenzyl Ether (DBE) | Sigma | 108014 | Clearing |
| Tetrahydrofuran (THF) | Sigma | 186562 | Dehydration |
| Methanol | Sigma | 179957 | Pre-treatment |
| Hydrogen Peroxide | Sigma | H1009 | Bleaching |
| Krazy Glue | krazyglue.com (Elmer's Products) | KG385 | Sample mounting |
| Glass slide | Globe Scientific Inc. | 1380-20 | Sample mounting |

**Supplementary Table 5, Data acquisition and processing details for data imaged with reflective diSPIM and reflective LLS presented in Fig. 4**

| Samples | | *C. elegans* embryos expressing GCaMP3 | U2OS cells expressing mEmerald-α-Actinin |
|---|---|---|---|
| **Figures/Videos** | | Fig. 4 c<br>Sup. Video 14 | Fig. 4f<br>Sup. Video 16 |
| **Microscope** | | Reflective diSPIM | Reflective LLS |
| **View number** | | 2 normal views<br>2 mirror view | 1 normal view<br>1 mirror view |
| **Color number** | | 1 | 1 |
| **Acquisition** | Slices per view | 60 | 300 |
| | Exposure per slice | 5 ms | 8 ms |
| | Axial step size | 1 μm | 0.4 μm |
| | acquisition time (each time point, all views) | 0.3 s | 2.4 s |
| | time interval | 0.35 s | 2.5 s |
| | total time points | 155 | 100 |
| | total acquisition time | ~54 s | ~250 s |
| **Image size (after interpolation)** | | 360 x 310 x 360 x 2 | 425 x 540 x 256 x 2 |
| **Total data size** | | 12.5 G voxels,<br>25 GB, 16 bit | 12 G voxels,<br>24 GB, 16 bit |
| **Data processing (registration /deconvolution)** | Registration time (each time point) | 10.6 min @ CPU<br>6.8 s @ GPU | 16 min @ CPU<br>12 s @ GPU |
| | Deconvolution time (each time point) | 14 min @ Trad Decon<br>1.4 min @ WB Decon | 2 h @ Trad Decon<br>8 min @ WB Decon |
| | total deconvolution time (all time points) | 36 h @ Trad Decon<br>3.6 h @ WB Decon | 8.3 day @ Trad Decon<br>13.3 h @ WB Decon |
| | Deep learning training volumes | 100 | 80 |
| **Data processing (deep learning)** | Deep learning training time | 10.8 h | 8.2 h |
| | Deep learning validation time | 1.68 s | 2 s |
| | Total processing time for (all time points) | 260 s | 200 s |

### Supplementary Note 1, Using an unmatched projector/back projector

The modified Richardson-Lucy algorithm

$$e_{k+1} = e_k \left\{\left[\frac{i}{e_k * f}\right] * b\right\} \quad (1)$$

seeks to solve a linear system of equations

$$\boldsymbol{i} = F\boldsymbol{e}, \quad (2)$$

where the vector $\boldsymbol{i}$ represents the measured image data, the vector $\boldsymbol{e}$ represents the desired deconvolved estimate image, and the matrix $F$ represents convolution with a point spread function $f$. If the 1D image vectors are formed by lexicographically concatenating the rows of the images into single column vectors, then the matrix $F$ is very structured: it is block circulant with circulant blocks (BCCB).

Equation (1) can be recast in this matrix-vector notation using

$$e_{k+1} = e_k \left\{B\left[\frac{i}{Fe_k}\right]\right\}, \quad (3)$$

where the division operation in square brackets is performed elementwise and the backprojection matrix $B$ implements convolution with a backprojection kernel $b$. The matrix $B$ is also BCCB. Note that this form of the R-L equation without normalizing prefactors assumes the PSF $f$ and backprojection kernel $b$ are both normalized to sum to one, which preserves counts across iterations. The standard R-L equation would set $B = F^T$, where $T$ denotes transpose, which corresponds to convolution with a flipped version of the PSF $f$ (or equivalently to convolution with the PSF $f$ itself if $f$ is symmetric).

In the context of image reconstruction in medical imaging, where the R-L algorithm is widely used under the name maximum likelihood emission maximization (MLEM), Zeng and Gullberg showed that the R-L algorithm can be accelerated through careful choice of the backprojection matrix $B$[1]. Specifically, the R-L algorithm was shown to move more rapidly toward desirable reconstructed images when the eigenvalue spectrum of the matrix product $BF$ is flatter (i.e., the eigenvalues cluster closely together). Zeng and Gullberg discuss additional constraints on the back projector $B$ that guarantee convergence to the same solution as when using the traditional matched back projector $B = F^T$. However, these constraints are mainly important for the additional class of Landweber iterations they study. R-L algorithms are rarely run to convergence since that yields unacceptably noisy solutions. Moreover, as in Zeng and Gullberg, we truncate any negatives that arise in the modified R-L iteration at each iteration, a step that invalidates any analysis based on these convergence constraints.

Thus, the key result from Zeng and Gullberg that we invoke (and validate) in this paper is that R-L can be accelerated by choosing the backprojection matrix $B$ such that the eigenvalue spectrum of the matrix product $BF$ is as flat as possible.

It is straightforward to determine the eigenvalue spectrum of $BF$. Both $B$ and $F$ are BCCB and circulant matrices can be diagonalized by the discrete Fourier transform (DFT) matrix:

$$B = QD_BQ^\dagger, \quad (4)$$

where $Q$ is a matrix representing application of the DFT and $Q^\dagger$ is its conjugate transpose[2]. $D_B$ is a diagonal matrix whose diagonal elements are given by the Fourier transform of the backprojection kernel $b$. These diagonal elements are also the eigenvalues of $B$. Thus

$$D_B = \mathrm{Diag}[\mathrm{DFT}(b)], \quad (5)$$

where the elements of $\mathrm{DFT(b)}$ are appropriately organized into a 1D vector. This is a formal way of expressing the well-known fact that convolution in the spatial domain is equivalent to multiplication in the Fourier domain.

Likewise, for the forward projection matrix $F$ we have

$$F = QD_F Q^\dagger, \tag{6}$$

where

$$D_F = \mathrm{Diag}[\mathrm{DFT}(f)]. \tag{7}$$

Note that the DFT of the system PSF is the system optical transfer function (OTF). I.e,

$$OTF_f \equiv \mathrm{DFT}(f). \tag{8}$$

The product $BF$ is thus given by

$$BF = QD_B Q^\dagger Q D_F Q^\dagger. \tag{9}$$

Invoking the fact that the DFT matrix is unitary, i.e., $QQ^\dagger = Q^\dagger Q = I$, we have

$$BF = QD_B D_F Q^\dagger. \tag{10}$$

We see that $BF$ is also a circulant matrix since it is diagonalized by the DFT and that its eigenvalue spectrum is given by the product of the diagonal matrices $D_B D_F$. Thus, the eigenvalue spectrum of $BF$ is given by the product $\mathrm{DFT}(b)\mathrm{DFT}(f)$, the elementwise product of the DFTs of the kernels used in the forward and back projector steps of the algorithm.

Let's consider a few cases:

1. In the usual case with a symmetric PSF and matched back projector, we have $B = F^T = F$ and so the eigenvalues of $BF$ will be given by $[\mathrm{DFT}(f)]^2 = OTF_f^2$, the squared OTF of the system. This is not especially flat.
2. In previous efforts to accelerate R-L based deconvolution[3], the kernel $b$ has been constructed in such a way that it is narrower than the kernel $f$. For Gaussian-like kernels, this makes $\mathrm{DFT}(b)$ broader and higher than $\mathrm{DFT}(f)$ and thus $\mathrm{DFT}(b)\mathrm{DFT}(f) > [\mathrm{DFT}(f)]^2$. This gives a flatter eigenvalue spectrum than in case 1, consistent with the finding of faster production of the resolution-limited result.
3. The flattest possible eigenvalue spectrum for $BF$ would be obtained if $\mathrm{DFT}(b) \sim \frac{1}{\mathrm{DFT}(f)} = \frac{1}{\mathrm{OTF_f}}$. This implies we should construct the backprojection kernel $b$ such that the inverse of its DFT approximates the system OTF. Obviously $OTF_f$ goes to zero or falls below the noise floor at some point for most PSFs $f$, but this suggests using a kernel corresponding to an inverse OTF filter with appropriate apodization. This is the basis for the Wiener-Butterworth kernel studied in this paper.

1 Zeng, G. L. & Gullberg, G. T. Unmatched Projector/Backprojector Pairs in an Iterative Reconstruction Algorithm. *IEEE Trans Med Imaging* **19**, 548-555 (2000).

2 Hunt, B. R. The application of constrained least squares estimation to image restoration by digital computers. *IEEE Trans. Comp.* **22**, 805-812 (1973).

3 Preibisch, S. *et al.* Efficient Bayesian-based multiview deconvolution. *Nat Methods* **11**, 645-648 (2014).

### Supplementary Note 2, Generating unmatched back projectors

**Supplementary Note 1** discusses how acceleration of the R-L algorithm can be achieved by using an unmatched back projector that provides a flatter product of the Fourier transforms of forward and backwards projectors, $|DFT(b)DFT(f)|$. This section describes how we create such unmatched back projectors, including the Gaussian back projector, Butterworth back projector and Wiener-Butterworth back projector. We also provide the parameters that we use in implementing these filters in our manuscript. Consider a fixed coordinate system $(x, y, z)$ in 3-dimensional (3D) space and the corresponding spatial frequency coordinate system $(k_x, k_y, k_z)$ in the 3D Fourier domain. For simplicity, we will also assume the long axis of the forward projector aligns along the axial direction, i.e. along the $z$-axis.

Given a forward projector $f(x, y, z)$, the traditional R-L algorithm defines the back projector as:

$$b(x, y, z) = \hat{f}(x, y, z), \tag{11}$$

where $\hat{f}$ is the transpose of $f$.

If we transition to discrete notation, the transpose can be expressed as

$$b(i, j, k) = \hat{f}(i, j, k) = f(M - i + 1, N - j + 1, L - k + 1), 1 \leq i, j, k \leq M, N, L \tag{12}$$

where the $M, N$ and $L$ denote the dimensions of the 3D image and $i, j$ and $k$ are the voxel indices.

In the Fourier domain, the discrete Fourier transform of the back projector is the conjugate of the discrete Fourier transform of the forward projector:

$$B_{Trad}(k_x, k_y, k_z) = DFT(b) = conj[DFT(f)]. \tag{13}$$

#### 1. Gaussian back projector

The Gaussian back projector uses a 3D Gaussian function with full width at half maximum (FWHM) matching the FWHM of the forward projector (i.e. the microscope PSF). The 3D normal Gaussian distribution is defined as

$$h(x, y, z) = a \exp\left(-\frac{x^2}{2\sigma_x^2} - \frac{y^2}{2\sigma_y^2} - \frac{z^2}{2\sigma_z^2}\right), \tag{14}$$

with $a = \frac{1}{(2\pi)^{3/2}\sigma_x\sigma_y\sigma_z}$, and $\sigma_x, \sigma_y, \sigma_z$ are the standard deviations in 3 dimensions, respectively.

The coefficient $a$ can be omitted since the back projector is normalized before deconvolution, and the relationship between the standard deviation and FWHM for a Gaussian distribution is:

$$\sigma_x = FWHM_x/(2\sqrt{2\ln 2}) \tag{15}$$

$$\sigma_y = FWHM_y/(2\sqrt{2\ln 2}) \tag{16}$$

$$\sigma_z = FWHM_z/(2\sqrt{2\ln 2})\,. \tag{17}$$

Then the Gaussian back projector is derived as:

$$b_{Gauss}(x,y,z) = \exp\left(-\frac{4\ln 2 \cdot x^2}{FWHM_x^2} - \frac{4\ln 2 \cdot y^2}{FWHM_y^2} - \frac{4\ln 2 \cdot z^2}{FWHM_z^2}\right), \tag{18}$$

where $FWHM_x$, $FWHM_y$, $FWHM_z$ are the FWHMs of the Gaussian back projector in x, y, and z, respectively, and are equivalent to the FWHMs of $f$ and $\hat{f}$.

Using the same sized FWHMs as $f$ in the spatial domain implies that the Gaussian back projector possesses a greater spatial frequency extent i than the traditional back projector in the Fourier domain, and as such results in a slightly flatter spectral product $|DFT(b)DFT(f)|$ as shown in **Figure S2.1**. This explains why we observe an acceleration with a Gaussian back projector compared to the traditional R-L back projector.

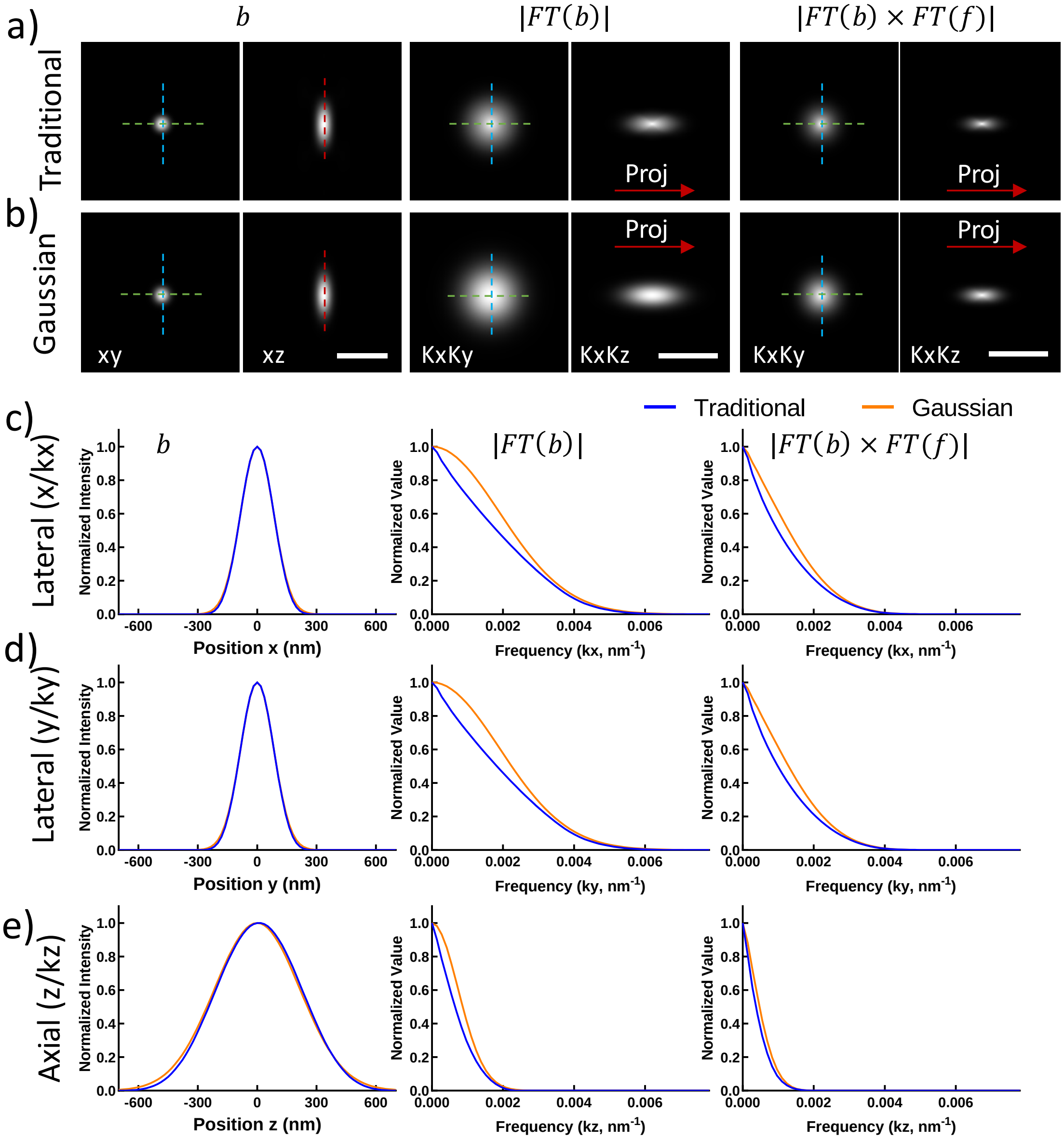

**Figure S2.1, Comparison of traditional and Gaussian back projectors with equivalent FWHM a), b)** Lateral and axial slices through the traditional (**a**) and Gaussian (**b**) back projectors for iSIM, shown in real space (left, b), Fourier space (middle, |FT(b)|) and the spectral product of forward and back projectors (right, |FT(b) FT(f)|). **c)** Lateral (x, kx) line profiles through the center of the slices showing in **a**, **b** (green dashed lines), comparing back projectors in real space (left), Fourier space (middle), and the spectral product (right). **d)** Lateral (y, ky) line profiles through the center of the slices shown in **a**, **b** (cyan dashed lines), comparing back projectors in real space (left), Fourier space (middle), and the spectral product in Fourier space (right). **e)** Axial (z or kz) profiles of the slices shown in **a**, **b**, comparing back projectors in real space (left), Fourier space (middle), and the spectral product (right). Note the left profiles correspond to line profiles through the center of the corresponding images in **a, b** (red dashed lines), while the middle and right profiles are maximum intensity projections in the kx direction as indicated by the red arrows ('Proj') in **a**, **b**, to capture the full extent of the spatial frequencies in the axial direction. Scale bars: **a**, **b**) 1 μm in left column, 1/100 $nm^{-1}$ in middle, right columns.

**2. Butterworth back projector**

To make the spectral product $|DFT(b)DFT(f)|$ flatter, one possibility is to make the discrete Fourier transform of the back projector $DFT(b)$ broader (which is equivalent to making the back projector narrower in the spatial domain). An extreme example is the Dirac delta function, i.e. the narrowest positive distribution in the spatial domain, which produces a uniform distribution in the Fourier domain. However, all microscopes function as low-pass filters, allowing only frequencies lower than the resolution limit to be transferred to the image. If the delta function is used as a back projector, higher spatial frequencies would be introduced into the image, thus boosting the noise during the R-L iteration as shown in **Figure S2.2**. To suppress this noise, we employ a Butterworth low-pass filter to ensure that the frequency contribution of the back projector remains within the resolution limit of the microscope. The one- dimensional Butterworth filter is defined as:

$$H(k) = \frac{1}{\sqrt{1+\varepsilon^2 k^{2n}}} \tag{19}$$

where $k$ is the spatial frequency in the Fourier domain, $n$ is the order of the filter and $\varepsilon$ is the maximum passband gain.

To obtain a 3D Butterworth filter, we decompose $k$ into 3 components, accounting for the microscope's resolution limit in each dimension:

$$k = \sqrt{\left(\frac{k_x}{k_{cx}}\right)^2 + \left(\frac{k_y}{k_{cy}}\right)^2 + \left(\frac{k_z}{k_{cz}}\right)^2}, \tag{20}$$

where the $k_{cx}, k_{cy}, k_{cz}$ are the cutoff frequencies corresponding the the resolution limit in the xyz dimensions, respectively. We then define the the Butterworth back projector in the Fourier domain, $FT(b)$,as:

$$B_{Butterworth}(k_x, k_y, k_z) = \frac{1}{\sqrt{1+\varepsilon^2\left[\left(\frac{k_x}{k_{cx}}\right)^2 + \left(\frac{k_y}{k_{cy}}\right)^2 + \left(\frac{k_y}{k_{cz}}\right)^2\right]^n}} \tag{21}$$

In the spatial domain, the Butterworth back projector can be obtained by an inverse Fourier transformation:

$$b_{Butterworth}(x,y,z) = FT^{-1}[B_{Butterworth}(k_x,k_y,k_z)]. \qquad (22)$$

The spectral intensity at the cutoff frequency is

$$\beta = B_{Butterworth}(k_{cx},0,0) = B_{Butterworth}(0,k_{cy},0) = B_{Butterworth}(0,0,k_{cz}) = \frac{1}{\sqrt{1+\varepsilon^2}}. \qquad (23)$$

$\beta$ is also referred to as the cutoff gain, i.e. it represents the spectral amplitude that is passed at the microscope's resolution limit.

The Butterworth back projector $B_{Butterworth}(k_x,k_y,k_z)$ (or $b_{Butterworth}(x,y,z)$ in the spatial domain) can thus be characterized by the cutoff gain $\beta$ and the filter order . $\beta$ is often set as a small value that suppresses high frequencies beyond the resolution limit, thus reducing the noise in the deconvolution (**Figure S2.2**). The filter order $n$ is related to the transition slope at the cutoff. Ideally the transition slope would be as steep as a "brick wall"; a higher filter order brings the filter closer to an ideal "brick wall". In practice however, this ideal transition slope is undesirable as it produces excessive ringing in the spatial domain.

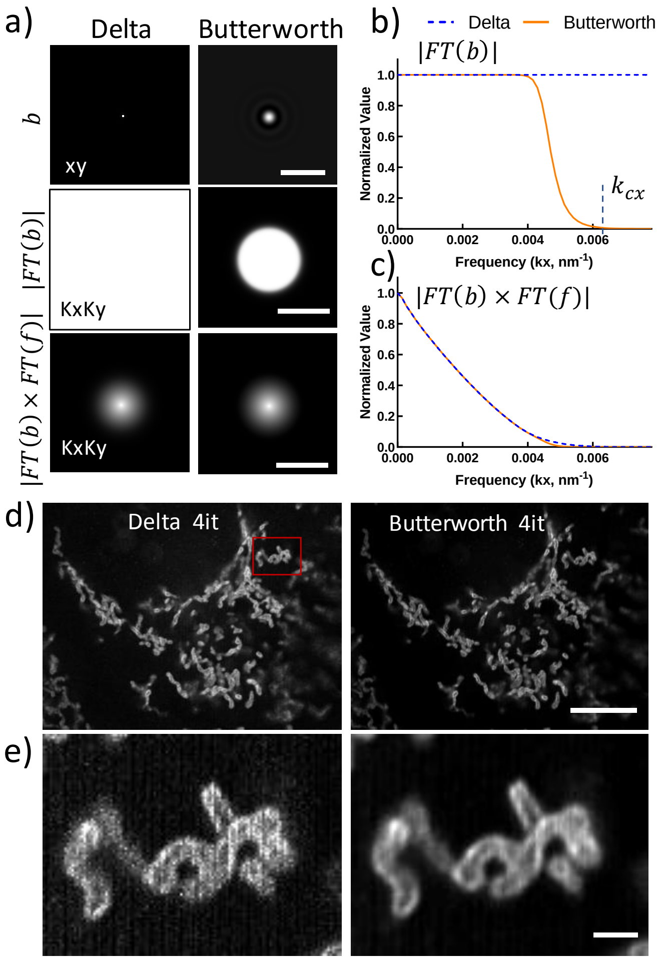

**Figure S2.2, Delta and Butterworth back projectors and corresponding R-L deconvolution. a)** Lateral slices through the Dirac delta (left) and Butterworth (right) back projectors for iSIM, shown in real space (top, b), Fourier space (middle, |FT(b)|) and the spectral product of forward and back projectors (bottom, |FT(b) FT(f)|). **b)** Lateral (kx) line profiles through the center of the image in **a,** middle row, comparing back projectors in Fourier space. The cutoff frequency $k_{cx}$ is indicated by a dotted line. **c)** Lateral (kx) line profiles through the center of the image in **a** bottom row, comparing the spectral product using the two different back projectors. **d)** R-L deconvolution of images of U2OS cells immunolabeled by Tomm20 Alexa-Fluor 488 and imaged with iSIM. Single planes from imaging stacks are shown, with iteration number (it) and back projector as indicated. **e)** Higher magnification views, corresponding to the red rectangular region in **d**, highlighting the additional noise that results when using the delta function back projector. Note this noise is suppressed when using the Butterworth back projector. Here we used $\beta = 0.001$ and $n = 8$ for the Butterworth back projector. Scale bars: **a**) 1 µm in top row, 1/100 $nm^{-1}$ in middle and bottom rows; **d**) 10 µm; **e**) 1 µm.

**3. Wiener-Butterworth back projector**

Finally, the choice that maximally flattens the spectral product $|DFT(b)DFT(f)|$ , sets the discrete Fourier transform of the back projector equal to the inverse of the discrete Fourier transform of the forward projector, i.e. $DFT(b) \approx 1/\,DFT(f)$. Obviously $DFT(f)$ goes to zero at some point, but one could consider

$$B_{Wiener}(k_x, k_y, k_z) = \frac{Conj[DFT(f)]}{[DFT(f)]^2 + \alpha}, \tag{24}$$

where $\alpha$ is a small value that prevents the denominator from becoming zero. This is like a classic Wiener filter that seeks to invert the forward projector $DFT(f)$ out to some point and then rolls off gently. However, when using such a filter, the results are highly sensitive to the choice of $\alpha$. The Wiener filter either fails to recover the best resolution possible with the microscope (if $\alpha$ is too large) or amplifies the noise and introduces more ringing artifacts (if $\alpha$ is too small, see also **Supplementary Fig. 4**). We would ideally circumvent these issues by combining the Wiener and Butterworth filters, obtaining a back projector that enables a flat spectral product but also suppresses spatial frequencies beyond the resolution limit. This is the Wiener-Butterworth back projector we use in our paper:

$$\begin{aligned} B_{WB}(k_x, k_y, k_z) &= B_{Wiener}(k_x, k_y, k_z) \times B_{Butterworth}(k_x, k_y, k_z) \\ &= \frac{Conj[DFT(f)]}{[DFT(f)]^2 + \alpha} \times \frac{1}{\sqrt{1 + \varepsilon^2 \left[ \left(\frac{k_x}{k_{cx}}\right)^2 + \left(\frac{k_y}{k_{cy}}\right)^2 + \left(\frac{k_z}{k_{cz}}\right)^2 \right]^n}}. \end{aligned} \tag{25}$$

The spatial analog of this filter can be derived by taking an inverse Fourier transform:

$$b(x, y, z)_{WB} = FT^{-1}[B_{WB}(k_x, k_y, k_z)]. \tag{26}$$

Again, we define the cutoff gain as

$$\beta = B_{WB}(k_{cx}, 0, 0) = B_{Wiener}(k_{cx}, 0, 0) \times \frac{1}{\sqrt{1 + \varepsilon^2}}, \tag{27}$$

so that as before $\beta$ represents the spectral amplitude that is passed at the microscope's resolution limit.

Given a forward projector $f$, we now need to determine three parameters to fully specify the Wiener-Butterworth back projector: $\alpha, \beta$ and $n$. $\alpha$ is a small value used to ensure that inverting the forward projector does not result in division by zero. As with the Butterworth back projector, the cutoff gain $\beta$ is set at a small value that suppresses spatial frequencies beyond the resolution limit, and the filter order $n$ is used to set the transition slope at the cutoff frequency. We'll discuss these parameters in more detail in the next section.

**4. Parameter configuration**

1) Parameter $\alpha$

As shown in equation (25), $\alpha$ is used to invert $DFT(f)$ in the first, Wiener filter term. We set $\alpha$ at a value that slightly boosts the spatial frequencies past the cutoff frequency, relying on the Butterworth filter term to suppress noise. In our experience, good results are obtained if $\alpha$ is set in the range $0.001 \sim 0.05$ (**Table S2.1**).

2) Parameter $\beta$

$\beta$ defines the spectral amplitude at the cutoff frequency or resolution limit, and in our experience good results are obtained when this is set at a small value in the range from 0.001 to 0.05. Alternatively one can also set $\beta$ based on the theoretical cutoff of the traditional R-L back projector $B(k_x, k_y, k_z)_{Traditional}$, or $DFT(\hat{f})$:

$$\beta = (B_{Trad}(k_{cx}, 0, 0) + B_{Trad}(0, k_{cy}, 0) + B_{Trad}(0, 0, k_{cz}))/3. \quad (28)$$

In this case, $\beta$ is an average of the 3 cutoff gains corresponding to each dimension of the traditional back projector. All $\beta$ values used for datasets in our paper are listed in **Table S2.1**.

3) Parameter $n$

The filter order $n$ is related to the flatness of the frequency distribution within the passband of the Butterworth filter. A higher $n$ would result in a flatter frequency response until the cutoff. However, as mentioned in previous sections, a higher $n$ would also make the transition slope closer to a hard cutoff like a "brick wall", causing the ringing artifacts in the spatial domain. In contrast, a lower order $n$ would make the transition slope gentler and sacrifice some spectral amplitude at spatial frequencies close to the cutoff. This choice in turn would require more iterations in R-L deconvolution. To empirically investigate this issue, we designed Wiener-Butterworth back projectors with different $n$ and tested them on an image acquired by iSIM. As shown in **Figure S2.3**, the deconvolution with $n = 8$ only needs 1 iteration, while using smaller $n$ requires more iterations for similar image quality.

We used $n$ in the range 4 - 10 for all datasets presented in our paper (**Table S2.1**). For the datasets acquired by single- or dual- view microscopes, $n = 8$ or 10 enables R-L deconvolution with 1 iteration. For the datasets acquired by quad-view light-sheet microscopy, reflective diSPIM and reflective lattice light-sheet microscopy, the deconvolution is more sensitive to the ringing caused by a higher filter order. We used $n = 5$, producing a resolution-limited result with only 2-5 iterations. We note that even using this relatively conservative choice, Wiener-Butterworth deconvolution is still 10-15x faster than traditional R-L deconvolution.

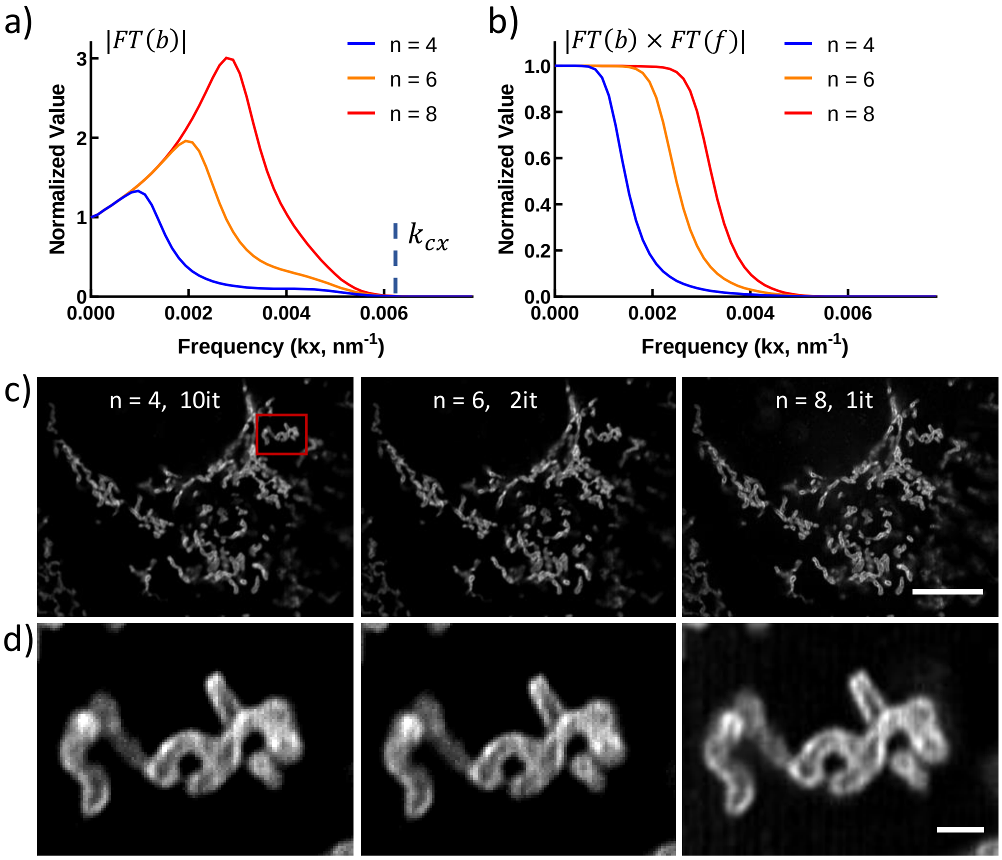


**Figure S2.3, Comparing Wiener-Butterworth back projectors with different filter orders. a)** Lateral (kx) line profiles through the central slice of Wiener-Butterworth back projector, for different filter orders. **b)** Lateral (kx) line profiles through the center of the spectral products $|DFT(b)DFT(f)|$ with filter orders as in **a**. **c)** R-L deconvolution results of images of U2OS cells immunolabeled by Tomm20 Alexa-488 and imaged with iSIM. Single planes from imaging stacks are shown, with filter orders and iteration number (it) as indicated. **d)** Higher magnification views, corresponding to the red rectangular region in **c**. Scale bars: **c**) 10 µm; **d**) 1 µm.

**Table S2.1, Parameters for all datasets used in Butterworth and Wiener-Butterworth Deconvolution**

| | | Butterworth | | | Wiener-Butterworth | | | |
|---|---|---|---|---|---|---|---|---|
| **Figures/Videos** | **Microscope** | $\boldsymbol{\beta}$ | $\boldsymbol{n}$ | **Iteration Number** | $\boldsymbol{\alpha}$ | $\boldsymbol{\beta}$ | $\boldsymbol{n}$ | **Iteration Number** |
| Fig 1 e, f,<br>Sup. Fig 4f<br>Sup. videos 1-3 | iSIM | 0.001 | 8 | 4 | 0.001 | 0.001 | 8 | 1 |
| Sup. Fig 5c, 6c | Widefield | 0.01 | 10 | 4 | 0.01 | 0.01 | 10 | 1 |
| Sup. Fig 6a | Confocal | 0.001 | 8 | 3 | 0.001 | 0.001 | 8 | 1 |
| Sup. Fig 6b | iSPIM | 0.01 | 10 | 5 | 0.05 | 0.01 | 10 | 1 |
| Figs 2a, j, 3a, d<br>Sup. Figs 7, 15<br>Sup. Videos 4, 5, 7, 10-14 | diSPIM | 0.01 | 10 | 3<br>(Sup. Fig 7) | 0.05 | 0.01 | 10 | 1 |
| Fig 2e<br>Sup. Fig 9<br>Sup. Video 6 | Quad-view light-sheet | -- | -- | -- | 0.01 | 0.05 | 5 | 5 |
| Fig 4c<br>Sup. Video 16 | Reflective diSPIM | -- | -- | -- | 0.01 | 0.02 | 5 | 2 |
| Fig 4f<br>Sup. Video 17 | Reflective LLS | -- | -- | -- | 0.01 | 0.02 | 5 | 4 |

### Supplementary Note 3, Deep learning for deconvolution

Our goal is to create deep learning models that simultaneously improve axial and lateral resolution, directly incorporating 3D information contained within the image stacks. A challenge when processing 3D image volumes is the associated computational burden. For example, processing a modestly sized 32-bit input image stack 40 MB in size, using only 8 convolution layers with 4 channels to output the result, implies that 40 x 4 x 8 = 1280 MB of memory is required. Recent evidence suggests that many complex tasks benefit from even deeper networks, with more layers[1-3]. Such increased depth and additional operations including activation functions and batch normalization[4] help the network to better represent[5] the underlying task and to stabilize the learning procedure[4], but at least double the computational memory required. Back-propagation, which is used to tune the network parameters according to the loss function, additionally doubles computational cost. One solution to the memory problem is to use down-sampling (pooling) in the learning model, e.g. as employed in the U-net architecture[6]. However, too much down-sampling results in a loss of detail unless the model contains enough parameters to compensate for the effect. Another solution is to crop the input data into multiple subimages. Selecting a random subset of these images for training can help the model learn the relationship between input and labels (ground truth) in a memory-efficient manner, but during the test procedure, stitching is required to combine the cropped images back to the original image size. Boundary artifacts[7], as well as differing local intensity/contrast in each restored sub-image can make this a nontrivial task. These issues motivated our search to find an efficient deep learning method for performing 3D deconvolution on the whole volumetric image input.

We adopted the concept of a densely-connected network[8] (Densenet) to establish this framework, terming our model 'DenseDeconNet'. The major components of our model are three dense blocks. These blocks use multiple dense connections between convolutional layers to extract relevant features from the image volumes, learning the deblurring necessary for image reconstruction. To efficiently process volumes of nearly 80 MB input size with a 14-layer model, another important concept we incorporated is the use of down-sampling and up-sampling operations, decreasing the computational memory required by at least 50%.

#### 1. Model architecture

The DenseDeconNet Framework is shown in **Figure S3.1**. Our DenseDeconNet adopts a fully convolutional architecture[9] for 3D image data, taking full advantage of the 3D geometric cues for effective volume-to-volume image restoration and enabling the processing of input data of different sizes. Our model also incorporates dense connections to create short paths from a layer to all its subsequent layers. These connections improve the usability of early layers' features, which decreases the number of parameters needed to learn redundant feature maps, thus making the model easier to train.

As shown in **Figure S3.1**, our network consists mainly of three dense blocks, with additional operations before and after the blocks. The convolutional layer before dense block 1 is used to increase the channel number (i.e., the number of feature maps), sampling the input data with dimension w x h x d for later skip connection after dense block 3. The convolution in this layer is performed with [3x3x3] kernel and [1x1x1] stride, and is followed by a batch normalization (BN) and rectified linear unit (Relu) activation function[10]. The number of output channels (denoted as $O_{11}$) is 4, resulting in an output of size of w x h x d x 4.

After $O_{11}$, dense block 1 contains three other convolutional layers with output denoted $O_{12}$, $O_{13}$, and $O_{14}$. The operations in each convolutional layer also include a convolution (Conv), a BN, and a Relu activation function. The convolution kernel size is set to [3x3x3] with stride [1x1x1]. These operations are used to extract higher-level features from previous layers. We denote the series of operation in each layer as $T(\bullet)$, thus $O_{12}$ can be computed from $O_{11}$ as:

$$O_{12} = T(O_{11})$$

We use $O_{12}$ to compute $O_{13}$ and to create a short path for delivering the feature map from $O_{11}$ to $O_{13}$ as follows. First, we concatenate the output channels $O_{12}$ and $O_{11}$, denoting this operation as $(O_{12} \oplus O_{11})$, where $\oplus$ represents the concatenate operation (Concat). Then we use this concatenation as the input of next layer so that $O_{13}$ is computed as:

$$O_{13} = T(O_{12} \oplus O_{11})$$

Similarly, short paths for $O_{11}$ and $O_{12}$ to $O_{14}$ are created and concatenated with $O_{13}$ as the input to $O_{14}$:

$$O_{14} = T(O_{13} \oplus O_{12} \oplus O_{11})$$

Concatenation helps subsequent layers to reuse previous feature maps, but it also causes the inclusion of redundant features and increases the number of parameters. For example, if the number of output channels in $O_{12}$ is 8 and the number of output channels for $O_{11}$ is 4, then the number of input channels for $O_{13}$ is 12, and the number of input channels for $O_{14}$ is 16. Because $O_{12}$ is derived from $O_{11}$, their feature maps may be similar. To lessen this redundancy, the number of output channels in $O_{13}$ and $O_{14}$ is limited to 4 by using the convolutional layers to select the most important features. This attribute also minimizes extraneous computational burden.

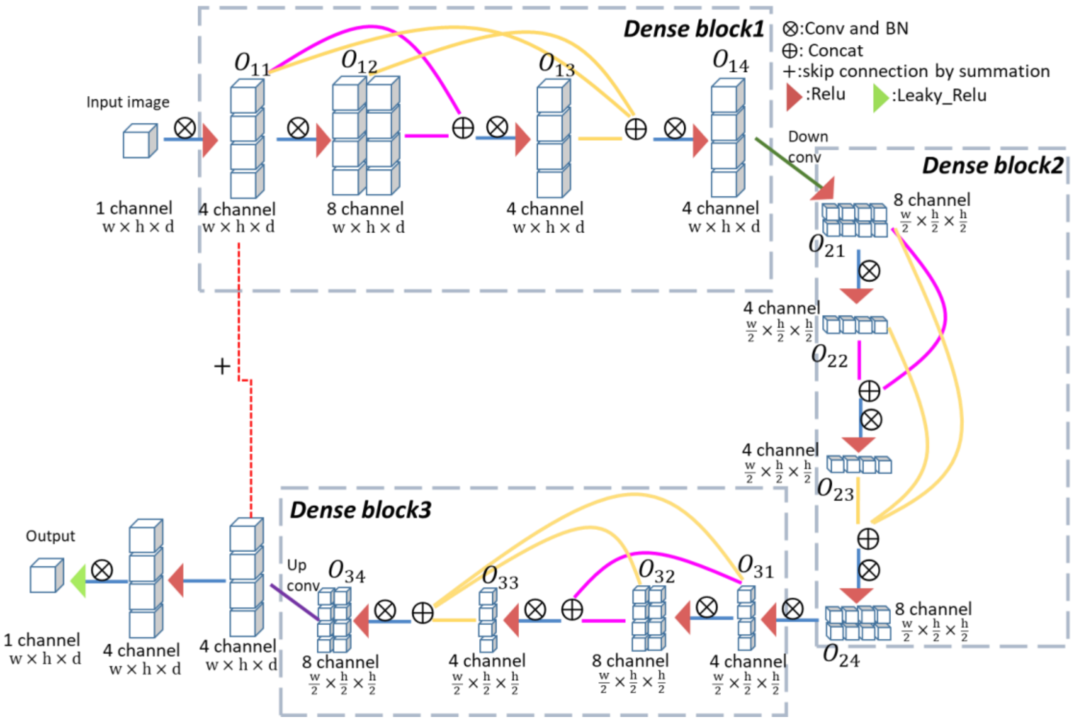


**Figure S3.1, The architecture of our DenseDeconNet neural network.** This fully convolutional network consists of three dense blocks (dashed rectangles), one down-sampling operation ('Down conv'), one up-sampling operation ('Up conv') and one skip connection (red dotted line). All operations are implemented on 3D data. Conv: convolution; BN: batch normalization; Relu: rectifying linear unit; Leaky_Relu: leaky rectifying unit; w: width; h: height; d: depth; Concat: concatenation. $O_{b1}$, $O_{b2}$, $O_{b3}$, $O_{b4}$, with $b$=1, 2, 3 represent the output of convolutional layers in each dense block. Blue lines signify the extraction and transfer of feature maps in all layers. Magenta lines signify the concatenation of the first two layers in each block, yellow lines signify the concatenation of the first three layers in each block.

After dense block 1, we use a down-sampling layer to decrease the size of the feature map $O_{14}$ by half to w/2 x h/2 x d/2, but include 8 channels. The result $O_{21}$ serves as the beginning of dense block 2. The down-sampling layer consists of a convolution with [2x2x2] kernel and [2x2x2] stride (denoted Down conv), a BN and a Relu activation. Constructing $O_{21}$ in this way enables efficient memory usage while also increasing the receptive field to incorporate more information surrounding each voxel.

Dense block 2 also contains other 3 convolutional layers with similar function to dense block 1, with outputs denoted $O_{22}$, $O_{23}$, $O_{24}$, respectively. Again, the number of output channels are limited to 4 or 8, and we rely on the network learning the most salient features to avoid redundancy.

After dense block 2, a convolutional layer connects dense block 3 to increase the depth and receptive field of the model. This layer consists of a convolution with [3x3x3] kernel and [1x1x1] stride, a BN and a Relu activation and compresses the number of feature maps from 8 in the last output of dense block 2 (i.e., $O_{24}$) to 4 in the input of dense block 3 (i.e., $O_{31}$).

Dense block 3 contains 3 other convolutional layers with outputs $O_{32}$, $O_{33}$, $O_{34}$, respectively, with characteristics shown in **Fig. S3.1**. After dense block 3, the network needs to recover the size of the original input image. Therefore, we use an up-sampling component composed of a transport convolution with [2x2x2] kernel and [2x2x2] stride (Up conv) and a BN. However, the transport convolution usually introduces checkerboard artifacts[7], which decreases output quality. To remedy this problem, we use a long skip connection that sums the output of $O_{11}$ and the up-sampled output of $O_{34}$. Next, we implement a nonlinear Relu operation. A final convolution layer is used to output the result. Because the gradient may vanish when the input of the Relu operation is negative, we used a Leaky-Relu operation[11] in the last convolution layer to maintain the gradient. The total number of learned parameters in our DenseDeconNet are approximately 18 thousand.

We designed our objective (loss) function with three terms: the mean square error (*MSE*), the structural similarity (*SSIM*) index[12] and the minimum value of the output (*MIN*). The *MSE* is widely used as a fidelity term to make sure the difference between network outputs and ground truths is as small as possible, but it may lead to blurred edges and a loss in resolution and fine image structure[12]. Thus, we also used the *SSIM* to preserve the global structural similarity between the network output and the ground truth. Because the Leaky_Relu in the last convolutional layer permits negative values, we monitor the *MIN* of the output to avoid negative values. The *MSE* and the *MIN* complement each other, as the objective function tends to make the *MIN* value larger, but the *MSE* acts to limits the *MIN*. Specifically, the objective function we aim to minimize is:

$$L(l,o) = MSE(l,o) - \ln\left(\frac{1 + SSIM(l,o)}{2}\right) - \lambda MIN(o) \tag{29}$$

where *l* represents the ground truth, *o* represents the model output and ln is the natural logarithm function. The *MSE* is defined as:

$$MSE(l,o) = \frac{1}{whd}\sum_{k=1}^{d}\sum_{j=1}^{w}\sum_{i=1}^{h}\left(l(i,j,k) - o(i,j,k)\right)^2 \tag{30}$$

where d, w, h represents the depth, width and height of the ground truth volume. The *SSIM* is defined as:

$$SSIM(l,o) = \frac{(2\mu_l\mu_o + C_1)(2\sigma_{lo} + C_2)}{(\mu_l^2 + \mu_o^2 + C_1)(\sigma_l^2 + \sigma_o^2 + C_2)} \tag{31}$$

where $\mu_l$, $\mu_o$ are the mean values of $l, o$; ${\sigma_l}^2$, ${\sigma_o}^2$ are the variances of $l, o$; $\sigma_{lo}$ is the covariance of $l$ and $o$; and $C_1$ and $C_2$ are small constants that prevent the denominator from becoming zero. In our implementation, we set $C_1 = 1e^{-4}$ and $C_2 = 9e^{-4}$. *SSIM* is in the range $(0,1]$. When *SSIM* equals 1, the ground truth and output are identical. Values for *MSE* and *SSIM* for all datasets are reported in **Table S3.2**. The $\ln(\cdot)$ operation is used to keep the objective function positive. The last term *MIN* is the minimum value of the output: $MIN(o) = \min(output)$.

We use a tunable parameter $\lambda$ to control the influence of *MIN*. If during training, the output tends to shift towards negative values, we set $\lambda > 1$, otherwise, we set $\lambda < 1$ or $\lambda = 0$.

The network is optimized using the backpropagation algorithm with the adaptive moment estimation (Adam) optimizer[13] and a starting learning rate $r_0$ which decays during the training procedure according to:

$$r = r_0 * k^{\frac{global\_step}{decay\_step}} \tag{32}$$

where *k* is the decay rate, *global_step* represents the number of training iterations (updated after each iteration), and *decay_step* determines the decay period. The values of these parameters are given in **Table S3.1**.

DenseDeconNet is implemented with the Tensorflow framework version 1.4.0 and python version 3.5.2 in the Ubuntu 16.04.4 LTS operating system. Training was performed on a workstation equipped with 32 GB of memory, an Intel(R) Core (TM) i7 – 8700K, 3.70 GHz CPU, and two Nvidia GeForce GTX 1080 Ti GPUs with 11 GB memory each. Kernels in the convolution layers were randomly initialized with a Gaussian distribution (mean= 0, standard deviation= 0.1). For an input image 70 MB in size, fully training the network with 10000 iterations took ~60 h.

**2. Results**

We tested DenseDeconNet on images of membranes and nuclei in live *C. elegans* embryos acquired with diSPIM, images of GCaMP3 expression in live *C. elegans* embryos acquired with reflective diSPIM, and images of α-actinin in live cells acquired with reflective lattice light-sheet (LLS) microscopy. Ground truth data consisted of traditional Richardson Lucy (R-L) joint deconvolution with 10 iterations for diSPIM data (conventional glass coverslips and reflective coverslips), and R-L deconvolution with the Wiener-butterworth back-projector with 1 iteration for reflective lattice light-sheet data. Additional parameters for datasets are shown in **Table S3.1**. All data are derived from volumetric time-series ('4D' data); 80% of volumes are used for training and the remaining 20% for testing. The test data are randomly chosen; all results shown in the following figures and in the main text figures display only the test data. Different datasets have different input sizes, but the training batch size is always 1. All the input data and ground truth data are normalized as follows:

$$X_N = \frac{X - \min(X)}{\max(X) - \min(X)} \quad X\ is\ input\ data \tag{31}$$

We first investigated the performance of deep learning on images of live *C. elegans* embryos expressing a pan-membrane mCherry label and pan-nuclear GFP label, acquired with diSPIM. We used the single-view images as network inputs and the dual-view joint deconvolution results as the ground truth. As shown in **Fig. S3.2**, the output of our DenseDeconNet network closely resembles the dual-view traditional deconvolution (e.g., the details indicated by yellow and pink arrows), although with worsened axial resolution. Interestingly we found that the network retained some details that are lost in the dual-view ground truth (e.g. the details indicated by the green arrows disappeared in the dual-view deconvolution but are retained in our method). These results are likely due to the highly dynamic nature of the membrane during embryo development, resulting in slight differences between the two raw views (which are acquired 1 s apart) and thus a loss of fine structure in the reconstruction.

Results were similar for relatively large structures like nuclei (**Fig. S3.3**) in late embryogenesis, when the embryo moves and twists frequently inside the eggshell. As before, the output from DenseDeconNet worsened axial resolution relative to the dual-view joint deconvolution (red arrows). However, this effect is counterbalanced by the effects of motion blur, which hinder the registration of the two volumetric views and compromise the resulting deconvolution (orange arrows).

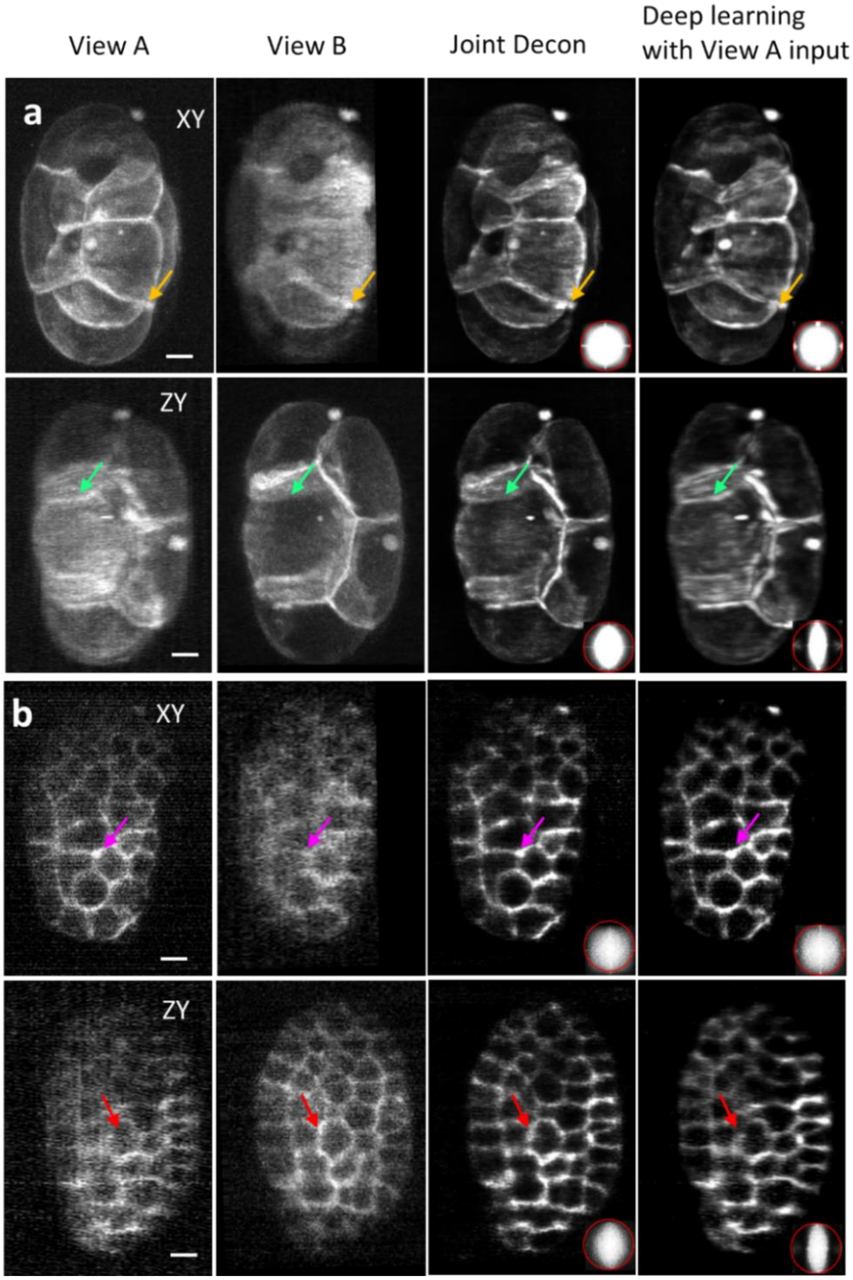


**Figure S3.2, Comparisons between raw single views, traditional joint deconvolution ('Joint Decon'), deep learning with single-view input, mCherry-membrane label in live *C. elegans* embryos. a)** Examples are from ~1 hour post fertilization and **b)** 4 hours post fertilization, XY and YZ maximum intensity projections are shown. In lateral XY views, both deep learning and traditional joint deconvolution produce similar reconstructions. The improved spatial resolution and contrast relative to the raw data reveal details that are otherwise obscured (magenta and yellow arrows). In axial YZ views, axial resolution in the learned result is not improved to to same extent as in the joint deconvolution (red arrows, see also OTF insets). However, the network results retain features evident in View A that are lost in the joint deconvolution (green arrows). OTFs averaged from all slices are shown at inset; red bounding box indicates a circle of 0.33 $\mu m^{-1}$. Scale bars: 5 µm.

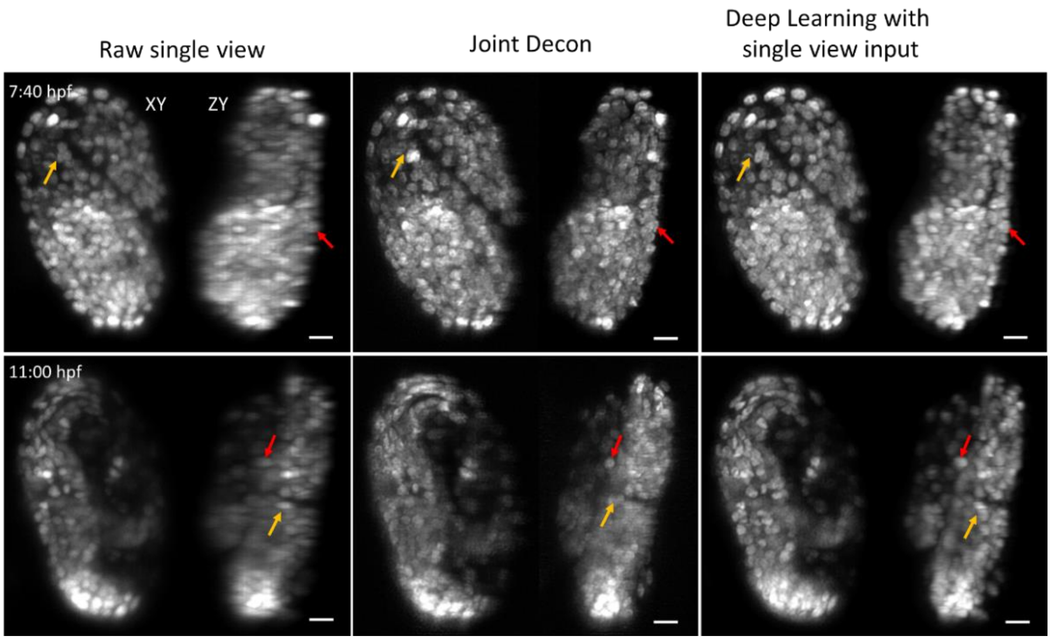


**Figure S3.3. Comparisons between raw single view, traditional joint deconvolution ('Joint Decon'), and deep learning, GFP-nuclear label in live *C. elegans* embryos.** Lateral (left) and axial (right) maximum intensity projections are shown for each condition; selected time points are shown both earlier in embryogenesis (top row) and later, post-twitching (bottom row). Note that the results shown for late twitching were obtained using training data from early twitching. Axial resolution is highest in the deconvolved result (red arrows). However, movement and twitching lead to degraded registration and deconvolution, resulting in improved reconstructions for many nuclei in the learned result (orange arrows). Scale bars: 5 μm.

We next investigated if we could further improve our reconstruction by using both orthogonal, registered volumes as input, and the joint deconvolution as the ground truth (**Supplementary Video 17**). The only change to the network is that the two inputs are concatenated, and the number of input channels is increased to two (**Fig. 4b**). For example, for two registered, orthogonally captured volumes in the reflected diSPIM each with dimensions [w x h x d], the input is now [w x h x d x 2]. However, like the single view case considered above, the output sizes of the first convolutional layer in both cases are [w x h x d x 4], and the following dense blocks and other operations are all the same as in the single view input case. Thus, operating the network with two inputs does not substantially increase computational burden. As shown in **Fig. S3.4** and **Supplementary Video 17**, using two inputs provides more information to the network, allowing resolution and resolution isotropy recovery, as well as the removal of epifluorescence contamination. With suitable training data, our DenseDeconNet thus facilitates the reconstruction of raw reflective diSPIM data.

Additional examples illustrating that the network performs better if two registered views are incorporated include registered dual-view GCaMP3 *C. elegans* embryo data acquired in the diSPIM (**Fig. S3.5, Supplementary Video 15**) and α-actinin data acquired by reflective lattice light-sheet (LLS) microscopy (**Fig. S3.6, Supplementary Video 16**).

For α-actinin data acquired by reflective LLS, the input data size for each view is 398 x 526 x 198 voxels, or 158 MB. This size exceeds the current memory limit for our network. Thus, in this case, we were forced to crop the data into multiple sub-volumes for training. Each subvolume then has size 210 x 271 x 198 voxels (42 MB), with a 22-pixel region in x and 16-pixel region in y overlap between neighboring subvolumes to avoid boundary artifacts. Note that when applying the training model to the test data, there is no need to crop the input data (i.e., we maintained the original size of the test data).

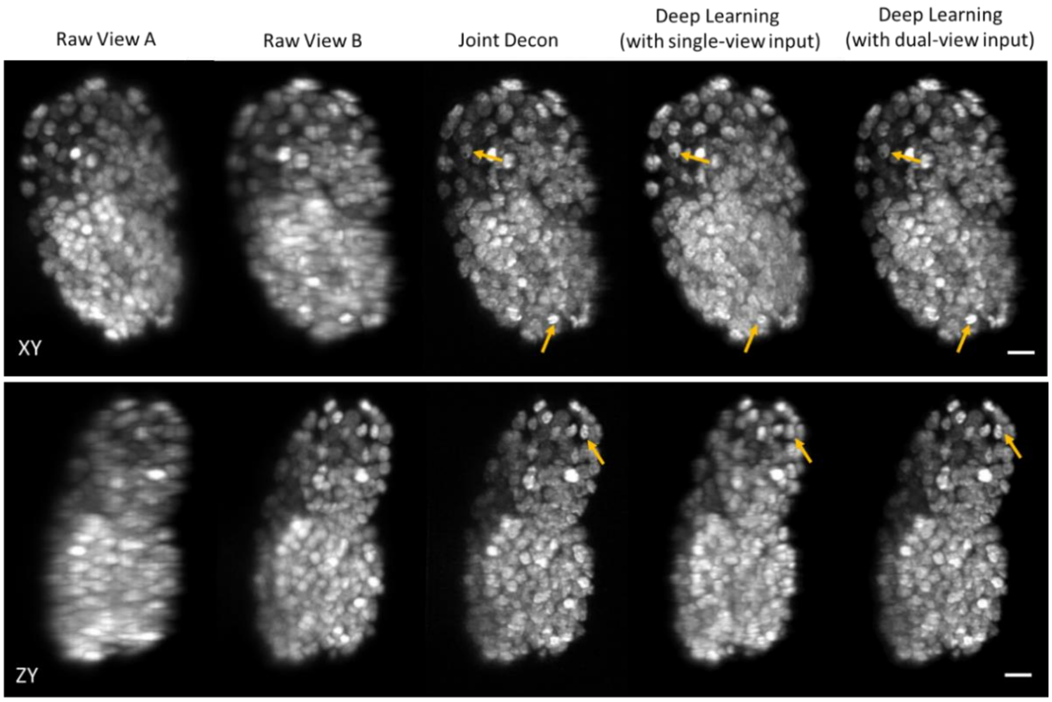


**Fig. S3.4, Comparing reconstructions of pan-nuclear *C. elegans* data with single- vs. dual-view input, acquired on the diSPIM.** Raw input data, traditional joint deconvolution ('Joint Decon'), deep learning with single-view input, and deep learning with dual-view input. Note the close resemblance (e.g. orange arrows) between the learned output with two inputs compared to the traditional deconvolution method. Scale bar: 5 µm. See also **Supplementary Video 17**.

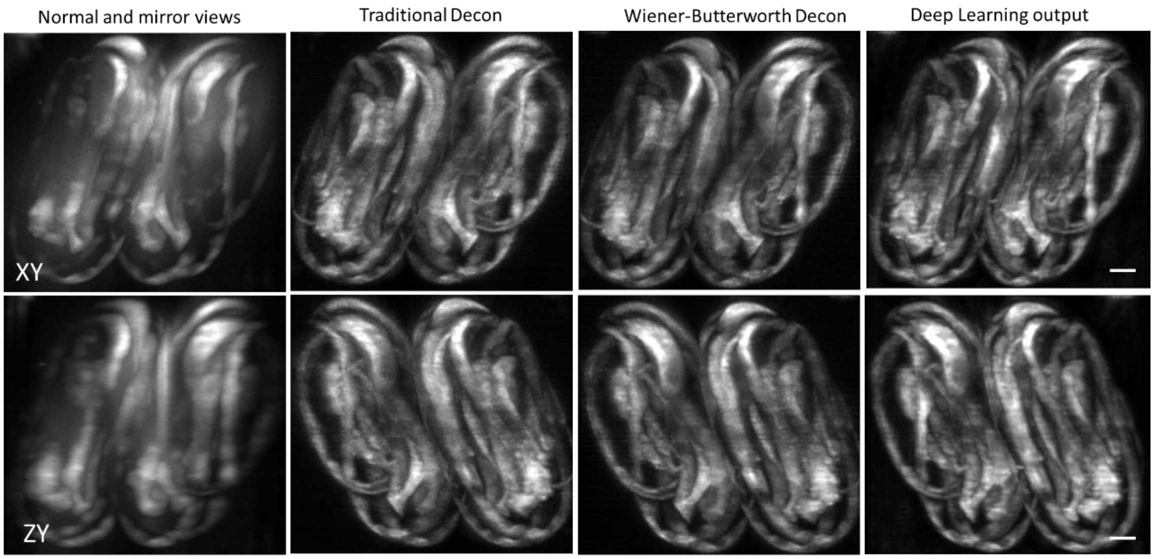


**Fig. S3.5, Comparisons between raw mirror data, traditional joint deconvolution ('Joint Decon'), Wiener-Butterworth joint deconvolution, and deep learning, GCaMP3 expression in twitching *C. elegans* embryos.** Lateral (top) and axial (bottom) maximum intensity projections are shown**.** Note the near equivalence of the deconvolved recontructions and the deep learning network output. Scale bars: 5 µm. See also **Supplementary Video 15**.

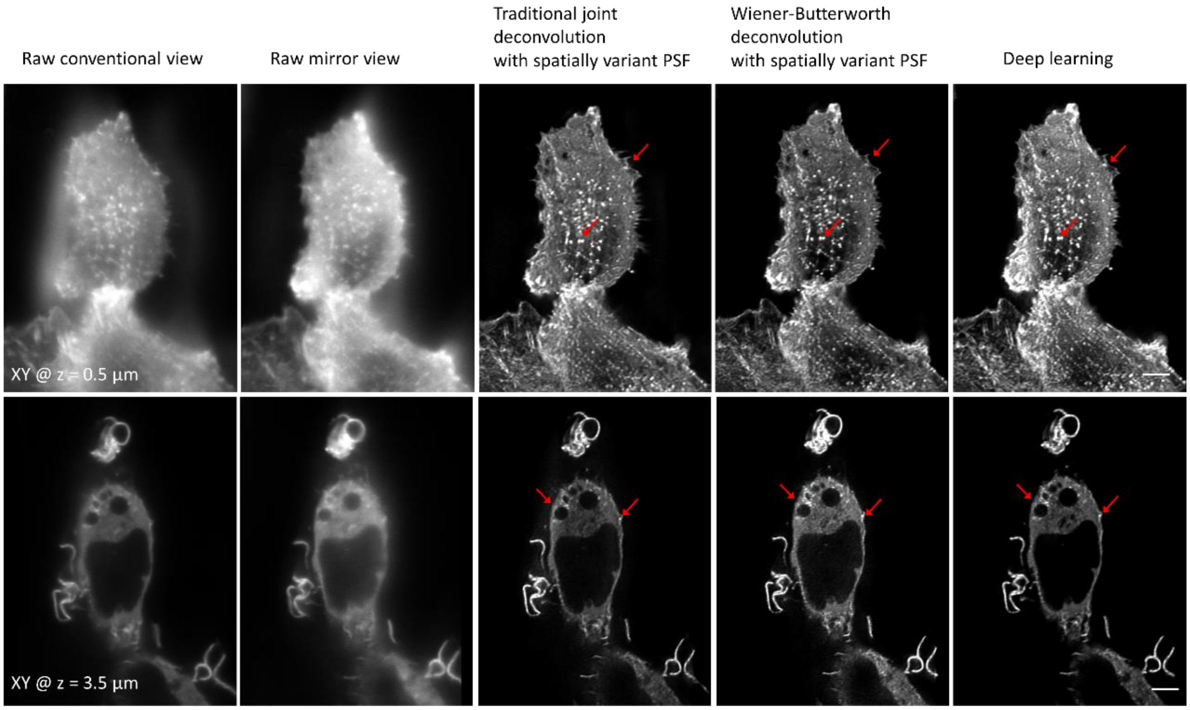


**Fig. S3.6, Post-processing mEmerald α-actinin data acquired by reflective LLS microscopy.** Two single-plane views at indicated axial depths from the coverlip are shown, columns indicate the two raw, registered inputs (conventional and mirror views), traditional joint deconvolution output, the Wiener-Butterworth deconvolution, and the output from DenseDeconNet deep learning (DL output). The deep learning output closely resembles the results obtained with traditional and Wiener-Butterworth deconvolution, including the retention of fine features (red arrows). Scale bar: 5 μm.

**Table S3.1, Parameters for all datasets used in deep learning**

| Samples | *C. elegans* embryo membrane | *C. elegans* embryo nuclei | | *C. elegans* embryos expressing GCaMP3 | U2OS cells expressing mEmerald-α-Actinin |
|---|---|---|---|---|---|
| **Microscope** | diSPIM | diSPIM | | reflective diSPIM | reflective LLS |
| **Input view number** | 1 | 1 | 2 | 2 | 2 |
| **Training volumes** | 160 | 180 | 180 | 100 | 80 |
| **Test volumes** | 40 | 110 | 110 | 55 | 20 |
| **Training volume size, voxels** | 240x340x246 (76.6 MB, 32 bit) | 240x360x240 (79 MB, 32 bit) | 240x360x240x2 (158 MB, 32 bit) | 340x170x340x2 (150 MB, 32 bit) | 210x271x198x2 (86 MB, 32 bit) |
| **Test volume size, voxels** | 240x340x246 (76.6 MB, 32 bit) | 240x360x240 (79 MB, 32 bit) | 240x360x240x2 (158 MB, 32 bit) | 340x310x340x2 (274 MB, 32 bit) | 398x526x198x2 (315 MB, 32 bit) |
| **Learning rate, $r_0$** | 0.07 | 0.04 | 0.004 | 0.004 | 0.005 |
| **Decay_step** | 200 | 150 | 600 | 600 | 400 |
| **Decay_rate, *k*** | 0.98 | 0.98 | 0.96 | 0.985 | 0.98 |
| **λ** | 1.5 | 1.2 | 1.3 | 1.3 | 1.3 |
| **Training iteration Number** | 8000 | 7000 | 7000 | 12000 | 13000 |
| **Training time** | ~57 h | ~56 h | ~56 h | ~10.8 h | ~8.2 h |
| **Test time for each volume** | ~1 s | ~0.81 s | ~1.23 s | ~1.68 s | ~2 s |

For data with dual-view inputs, the additional dimension (2) represents the concatenation/channel dimension. Training and model application were performed on a PC workstation equipped with 32 GB of memory, an Intel(R) Core (TM) i7 – 8700K, 3.70 GHz CPU, and two Nvidia GeForce GTX 1080 Ti GPU cards with 11 GB memory.

**Table S3.2, MSE and SSIM evaluation between joint deconvolution results (ground truth) and deep learning test outputs.**

| **Samples** | ***C. elegans* embryo membrane** | | ***C. elegans* embryo nuclei** | | | ***C. elegans* embryos expressing GCaMP3** | **U2OS cells expressing mEmerald-α-Actinin** |
|---|---|---|---|---|---|---|---|
| **Microscope** | diSPIM | | diSPIM | | | reflective diSPIM | reflective LLS |
| **Input view number** | 1 | | 1 | | 2 | 2 | 2 |
| **Time point** | @ 1:00 hpf | @ 4:00 hpf | @ 7:40 hpf | @ 11:00 hpf | @ 7:00 hpf | @ 6.3 s (#19 in the movie) | @ 5 s (#3 in the movie) |
| **MSE** | 3.0e-04 | 2.9e-04 | 6.6e-04 | 3.5e-03 | 0.6e-04 | 4.8e-04 | 5.0e-05 |
| **SSIM** | 0.684 | 0.923 | 0.805 | 0.440 | 0.986 | 0.923 | 0.965 |

MSE: mean square error; SSIM: structural similarity index. Except for the late twitching embryo with single view input, the MSE values are all < 0.001. We suspect this larger MSE value is due to the pronounced twitching late in embryogenesis. SSIM values show more variability, with single-view inputs for the membrane and late nuclear datasets resulting in lower SSIM values, presumably due to registration artifacts in the ground truth data. For datasets with two inputs, the similarity between the ground truth joint deconvolution and the deep learning output is very high. hpf: hours post fertilization.
